# Genomic underpinnings of lifespan allow prediction and reveal basis in modern risks

**DOI:** 10.1101/363036

**Authors:** Paul RHJ Timmers, Ninon Mounier, Kristi Läll, Krista Fischer, Zheng Ning, Xiao Feng, Andrew Bretherick, David W Clark, eQTLGen Consortium, Xia Shen, Tōnu Esko, Zoltán Kutalik, James F Wilson, Peter K Joshi

**Affiliations:** Centre for Global Health Research, Usher Institute of Population Health Sciences and Informatics, University of Edinburgh, Teviot Place, EH8 9AG Edinburgh, United Kingdom; Institute of Social and Preventive Medicine, University Hospital of Lausanne, 1010 Lausanne, Switzerland; Swiss Institute of Bioinformatics, 1015 Lausanne, Switzerland; Estonian Genome Center, University of Tartu, 51010 Tartu, Estonia; Institute of Mathematics and Statistics, University of Tartu, 50409 Tartu, Estonia; Department of Medical Epidemiology and Biostatistics, Karolinska Institutet, SE-171 77 Stockholm, Sweden; State Key Laboratory of Biocontrol, Guangdong Provincial Key Laboratory of Plant Resources, Key Laboratory of Biodiversity Dynamics and Conservation of Guangdong Higher Education Institutes, School of Life Sciences, Sun Yat-sen University, Guangzhou, China; MRC Human Genetics Unit, Institute of Genetics and Molecular Medicine, University of Edinburgh, Western General Hospital, Crewe Road, EH4 2XU Edinburgh, United Kingdom; Broad Inst of Harvard and MIT, Cambridge, 02142 MA, USA

## Abstract

We use a multi-stage genome-wide association of 1 million parental lifespans of genotyped subjects and data on mortality risk factors to validate previously unreplicated findings near *CDKN2B-AS1*, *ATXN2/BRAP*, *FURIN/FES*, *ZW10*, *PSORS1C3*, and 13q21.31, and identify and replicate novel findings near *GADD45G*, *KCNK3*, *LDLR*, *POM121C*, *ZC3HC1*, and *ABO*. We also validate previous findings near 5q33.3/EBF1 and FOXO3, whilst finding contradictory evidence at other loci. Gene set and tissue-specific analyses show that expression in foetal brain cells and adult dorsolateral prefrontal cortex is enriched for lifespan variation, as are gene pathways involving lipid proteins and homeostasis, vesicle-mediated transport, and synaptic function. Individual genetic variants that increase dementia, cardiovascular disease, and lung cancer –but not other cancers-explain the most variance, possibly reflecting modern susceptibilities, whilst cancer may act through many rare variants, or the environment. Resultant polygenic scores predict a mean lifespan difference of around five years of life across the deciles.

## Introduction

Human longevity is a highly complex trait, the product of myriad health, lifestyle, genetic, and environmental factors – alongside chance – and both individuals and society put much effort into its elongation. The extent to which lifespan can be explained by additive genetic variation in particular has been widely debated(1), with the most recent, and by far most well-powered study estimating heritability as 16.1% (SE = 0.4%)(2). Despite this modest heritability, extensive research, with some success, has gone into finding genetic variants influencing human survival, both in terms of age at death (3-6) and living to exceptional age (longevity) (6-12).

Studying the extremely long-lived, using a case-control design (7, 11-15) has the advantage of focusing on the truly remarkable, who also exhibit extreme healthspan and potentially unique genetic attributes (8, 16) whilst statistically focusing on those with most information, enhancing power at a given sample size, albeit from subjects that are hard to collect. However, although genome-wide association studies (GWAS) of mortality risk factors (such as cardiovascular traits and cancer) have had remarkable success (17-21), GWAS of longevity has proved more challenging, with only two robustly replicated, genome-wide significant associations (near *APOE*, *FOXO3*) having been made (7, 10).

This is a pity as understanding the effect of genetic variation on longevity or the overlapping but distinct (16) trait, lifespan, has the potential for fundamental understanding of the forces shaping how we age and our genome’s evolution as well as translational benefits. This was recognised almost fifty years ago when Lewontin speculated on a potential study of ABO blood groups and lifespan(22). However, he also identified two major obstacles: hundreds of thousands of lives would have to be recruited and an approximate follow-up of 35 years would be required. Recent developments, in particular the creation of a five hundred thousand subject population cohort, UK Biobank (23), and the use of parental lifespans and offspring genotypes(3) – an extension of the Wacholder’s kin-cohort method(24) – now enable researchers to discover genetic variants affecting survival from early middle age into the final decades of life without the long wait Lewontin prescribed, or the costly and often difficult recruitment of the extreme long-lived(7, 10, 15), albeit recognising the particular interest and statistical power of nonagenarians and centenarians.

The effectiveness of the kin-cohort approach was recently demonstrated by Pilling *et al*.(6), who increased the number of genome-wide significant associations with human survival from 4 to 25. Although their study is a major step forward in mapping the genetic architecture of lifespan, the design did not allow effect sizes to be readily interpreted or meta-analysed (for example as hazard ratios or years of life), and the novel genetic variants were not replicated, due to lack of an independent dataset.

Here, we leverage data from UK Biobank to carry out a genome-wide association study (GWAS) of parental survival beyond age 40, extending previous research by providing intuitive effect sizes and seeking replication in 26 independent European-heritage population cohorts (the LifeGen consortium(5)), yielding a combined sample of over 1 million parental lifespans. We then further supplement this with data from 58 GWAS on mortality risk factors to conduct a Bayesian prior-informed GWAS (iGWAS) and attempt a second round of replication in publicly available longevity studies.

We also examine association of lifespan-altering variants with diseases of subjects and their kin within UK Biobank (PheWAS) and an independent dataset(25) to provide insight into how genetic variants act to shorten or prolong lifespan. Finally, we implicate specific genes, biological pathways, and cell types in human survival, and use our findings to create and test predictions of lifespan, which, in theory, could have been made at birth.

## Result

### Genome-wide association analysis

We carried out GWAS of survival in a discovery sample of 635,205 parents (68% deceased) of unambiguously British ancestry subjects from UK Biobank, and a replication sample of 377,035 parents (47% deceased) of other European ancestry subjects from UK Biobank and 26 additional populations cohorts (LifeGen; Table S1). In each sample, we performed a sex-stratified analysis and then combined the allelic effects in fathers and mothers into a single parental survival association in two ways. First, we assumed genetic variants with common effect sizes (CES) for both parents, maximising power if the effect is indeed the same, secondly, we allowed for potentially different effect sizes (PDES), maximising power to detect sexually dimorphic variants, including those only affecting one sex.

We find fourteen genomic regions containing SNPs with genome-wide significant (P < 5×10^−8^) association in the discovery cohort, for one or both analyses (Fig. 1a). Ten of these loci have been previously reported using similar data (6), but only 4 have been successfully replicated so far (3-5, 10). We calculate the effects of previously replicated SNPs to be –1.06 (near *APOE*), –0.42 (*CHRNA3/5*), –0.76 (*LPA*), and +0.56 (*HLA-DQA1*) years of life gained per minor allele, estimated from the meta-analysis of discovery and replication cohorts. We also find evidence for replication (P < 0.05, one-sided test) for an additional 3 loci near *ATXN2/BRAP* (–0.28), *FURIN/FES* (–0.25), and *CDKN2B-AS1* (–0.25), which were previously identified at genome-wide significance by Pilling et al(6) (Table 1).

**Fig. 1:**
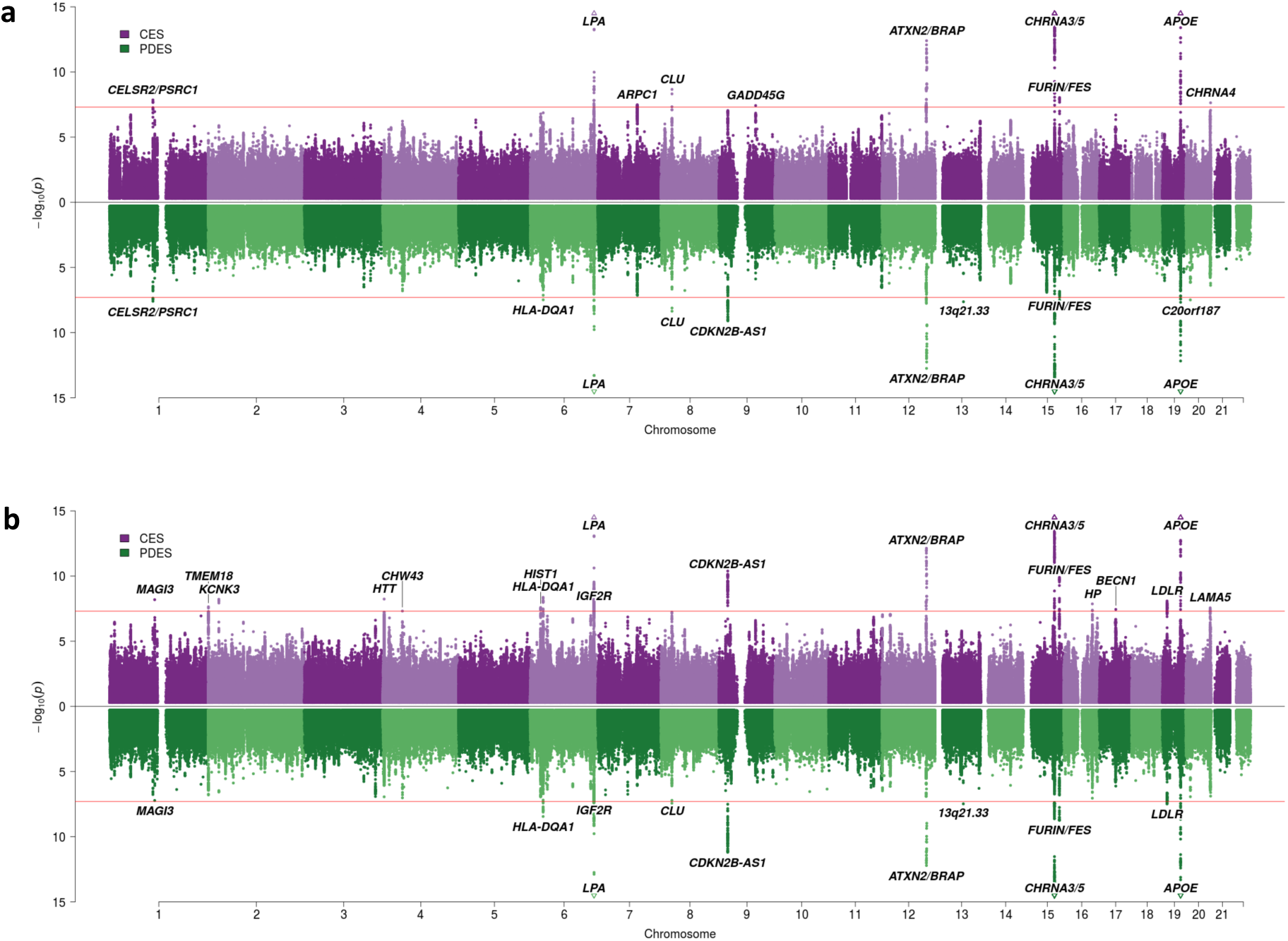
SNP associations with lifespan from the discovery cohort and discovery and replication meta-analysis cohorts, under common and potentially different assumptions of effects across sexes. (a) GWAS of UK Biobank discovery cohort, (b) GWAS of discovery and replication cohorts combined. In purple are the associations under the assumption of common SNP effects across sexes (CES); in green are the associations under the assumption of potentially different effects between sexes (PDES). P refers to the two-sided P values for association of allelic dosage on survival under the residualised Cox model. Annotated are the gene, cluster of genes, or cytogenetic band near the top SNP. The red line represents the genome-wide significance threshold (P = 5 x 10^-8^). P values have been capped at – log_10_(p) = 15 to better visualise associations close to genome-wide significance. SNPs with P values beyond this cap (near APOE, CHRNA3/5 and LPA) are represented by triangles.

**Table 1:**
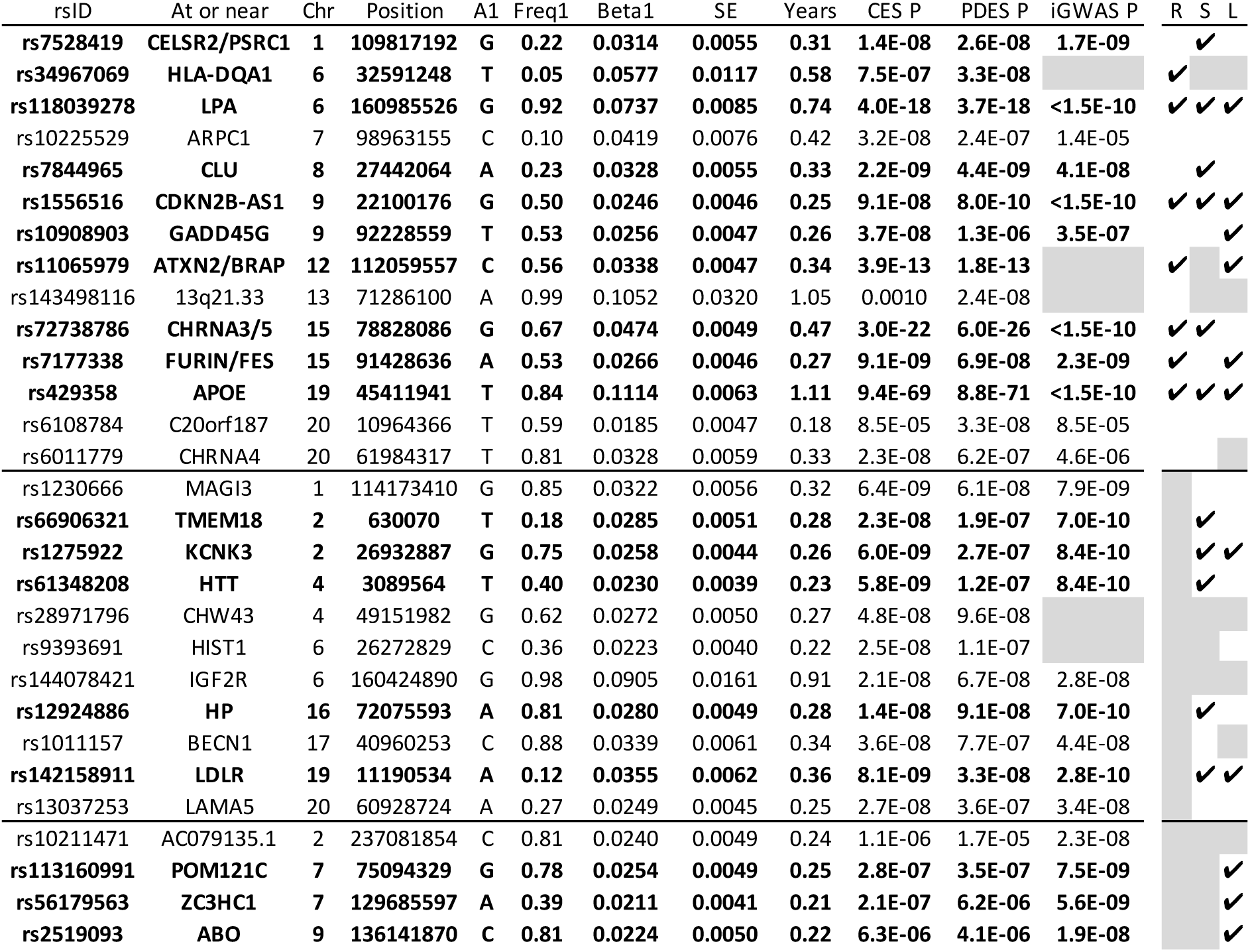
Twenty-nine genome-wide significant associations with lifespan in discovery, discovery and replication meta-analysis, or iGWAS. Top section contains SNPs reaching genome-wide significance in the discovery cohort, middle section contains SNPs reaching genome-wide significance in the meta-analysis of discovery and replication cohorts, bottom section contains additional SNPs reaching genome-wide significance in iGWAS. At or near – Gene, cluster of genes, or cytogenetic band in close proximity to lead SNP; Chr – Chromosome; Position – Base-pair position on chromosome (GRCh37); A1 – the effect allele, increasing lifespan; Freq1 – Frequency of the A1 allele; Betal – the log_e_(protection ratio) for carrying one copy of A1 under an additive dosage model which multiplied observed offspring genotype on parent effect by 2. For the top section, this is the effect reported in the discovery cohort, for the other two sections, this is the effect reported in the combined cohort; SE – Standard Error; Years – Years of life gained for a carrying one copy of the A1 allele; CES – Assumption of common effect size of A1 across sexes; PDES – Allowing for potentially different effect sizes of A1 across sexes; P – For the top section, the P value for the Wald test of association between imputed dosage and cox model residual in the discovery cohort. For the other sections, the same P value of the Wald test for the combined cohort; iGWAS P – The permutation P value of Bayes Factors against 7.2 billion null Bayes Factor distributions, hence limited to a minimum value of 1.4E-10; R – Replication P < 0.05 (CES one sided test, PDES two-sided test); S – Strengthened by iGWAS, i.e. iGWAS P value is lower than CES P value in the combined cohort or reaches minimum; L – associates with longevity (P<0.05, one-sided test) in external longevity studies. SNPs which replicate (lifespan or longevity) or gain additional support under iGWAS are in bold;. Grey – not applicable; for iGWAS this means the SNP was not included, for R (replication) this is not relevant when the replication data is used in discovery, S (iGWAS strengthening) as per iGWAS, L (longevity) – SNP or proxy not available in external dataset.

While we were unable to replicate the remaining seven loci, this may be due to lack of power for SNPs at or near 13q21.33 and *C20orf187*, where 95% confidence intervals (CIs) for replication effect overlap with both discovery effect Cis and zero. Conversely, lead SNPs near *CELSR2/PSRC1*, *ARPC1*, *CLU*, *GADD45G*, and *CHRNA4* do not appear to replicate due to small observed effects in the replication cohort rather than power (discovery and replication 95% CIs do not overlap and replication CI covers zero; Fig. 2a).

**Fig. 2.**
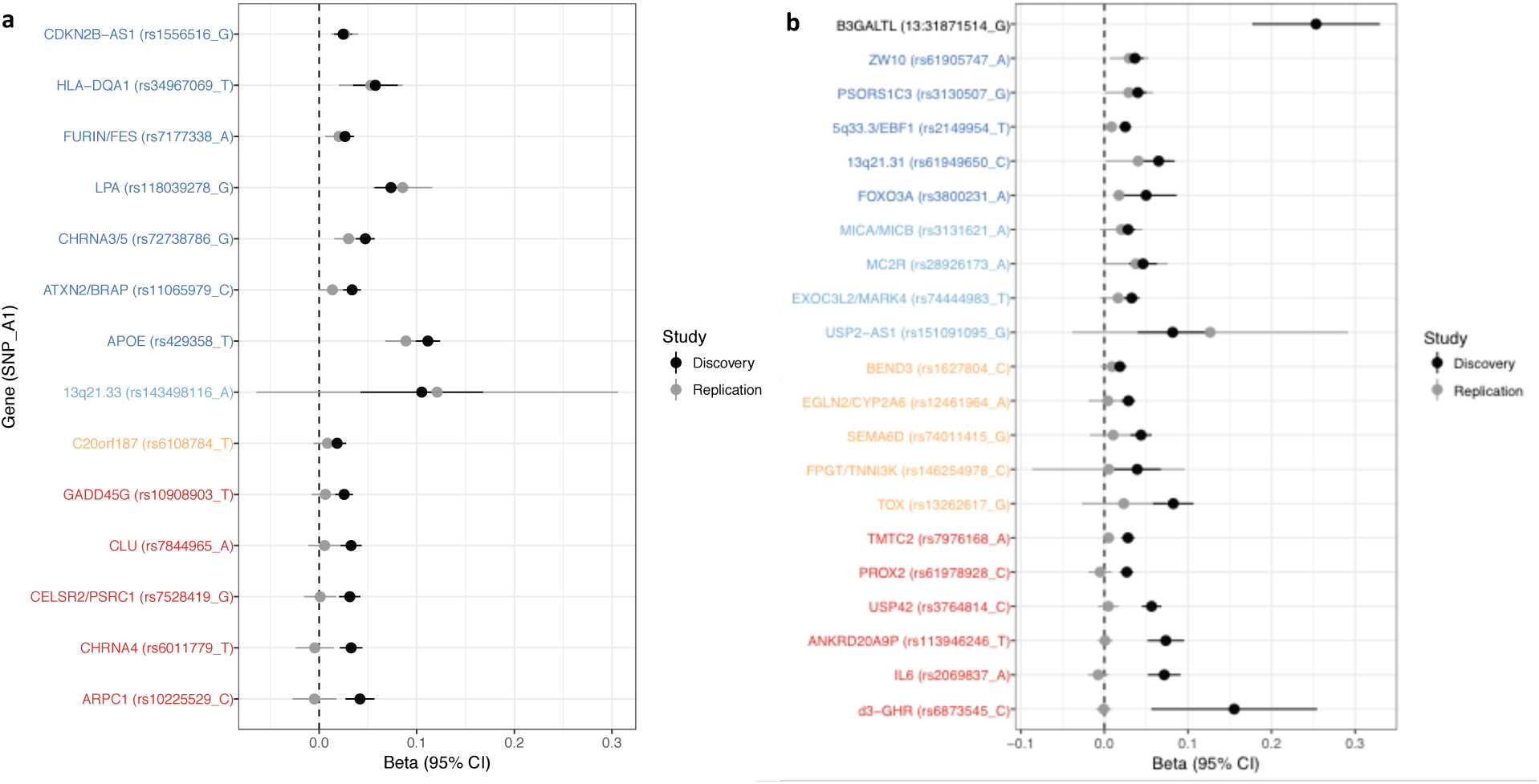
Validation of SNPs identified in our own and other studies using independent samples of European descent. Comparison of (inferred) effect sizes between discovery and replication cohorts. Panel A has discovery estimates taken from our own UKB Gen. British sample and replication estimates from the combined LifeGen + UKB European descent samples, other than those in discovery. Panel B has (sex-specific) discovery estimates inferred from other studies (6, 11-13, 15, 26) (see Methods and Table S3) and replication estimates from either LifeGen – to replicate Pilling et al. (6) – or the full dataset from Panel A (UKB discovery + replication combined). Gene names are as reported by discovery, and have been coloured based on overlap between confidence intervals (Cis) of effect estimates. Note, rs151091095 near USP2-AS1 is a proxy (r^2^ = 1.00) for rs139137459, the SNP reported by Pilling et al; rs113946246 near ANKRD20A9P is a proxy (r^2^ = 0.97) for rs2440012, the SNP reported by Zeng et al; no proxies could be found for 13:31871514_T_G. Dark blue – Nominal replication (P<0.05, one-sided test). Light blue – Cis overlap (P_different_>0.05) and cover zero, but replication estimate is closer to discovery than zero. Yellow – Cis overlap (P_different_>0.05) and cover zero, and replication estimate is closer to zero than discovery. Red – Cis do not overlap (P_different_<0.05) and replication estimate covers zero. Gene – Nearby gene(s) as reported by discovery. SNP – rsiD of SNP or proxy. A1 – Lifespan-increasing allele. Beta – the estimated log_e_(protection ratio) for one copy of the effect allele. Ci – Confidence interval.

Meta-analysis of our discovery and replication cohorts, totalling 1,012,240 parental lifespans, increased power further, with the P value for the lead SNP near *APOE* falling to 1.83×10^-85^. The analysis reveals 11 additional genome-wide significant loci at or near the following genes (and increase in lifespan per minor allele): *MAGI3* (–0.32), *TMEM18* (+0.28), *KCNK3* (–0.26), *HTT* (+0.23), *CHW43* (–0.27), *HIST1* (+0.22), *IGF2R* (–0.91), *HP* (–0.28), *BECN1* (–0.34), *LDLR* (+0.36), and *LAMA5* (+0.25) (Table 1, Fig. 1b).

The combined analysis has at least 50% power at genome-wide significance to detect any association between lifespan and frequent genetic variants (MAF > 0.3) with effect sizes of 0.25 years of life per minor allele or more, or common genetic variants (MAF > 0.1) with effect sizes of 0.56 years of life per minor allele or more.

We used the combined cohort to test six candidate SNPs previously reported at genome-wide significance to associate with longevity (11, 12, 15, 27) for association with lifespan. We find directionally consistent evidence of association (P < 0.05, one-sided test) for rs3800231 near *FOXO3* (+0.175) and rs2149954 near *5q33.3/EBF1* (+0.085) but find no effect on lifespan for SNPs near *IL6*, *ANKRD20A9P*, *USP42*, and *TMTC2*. We also tested a deletion, *d3-GHR*, reported to affect male lifespan by 10 years when homozygous(26), having converted the recessive effect into an expected apparent effect in our study for a truly recessive allele, but find no evidence of association with lifespan in our own male sample (Table S3;Fig. 2b).

We next attempted to validate additional survival SNPs found by Pilling *et al*.(6) using the independent replication cohort, LifeGen. We find evidence for replication (P < 0.05, one-sided test) for female-specific SNPs near *PSORS1C3* (–0.29 years per minor allele in LifeGen) and an intergenic region within 13q21.31 (+0.40), as well as one male-specific SNP near *ZW10* (–0.30). The remaining 10 SNPs for which LifeGen statistics were available did not replicate, primarily due to lack of power (95% CIs for LifeGen effect size overlap with both estimated discovery effect and zero), except for *PROX2*, where our independent result is not consistent with Pilling *et al.’s* discovery (95% CIs for effect estimates do not overlap and LifeGen CI covers zero) (Table S3; Fig. 2b).

### Mortality risk factor-informed GWAS (iGWAS)

We integrated 58 publicly available GWAS on mortality risk factors with our combined sample GWAS, creating Bayesian priors for each lifespan SNP effect based on causal effect estimates of independent risk factors on lifespan. This reveals an additional 4 genome-wide significant associations with lifespan (permutation P < 5 × 10^-8^) near *AC079135.1* (–0.24), *POM121C* (0. 25), *ZC3HC1* (+0.21) and *ABO* (–0.22) (Fig. 3, Table S10) where reported effects in brackets represent additional years of life per minor allele in the standard GWAS. A total of 82 independent SNPs associate with lifespan when allowing for a 1% false discovery rate (FDR) (Table S10).

**Fig. 3.**
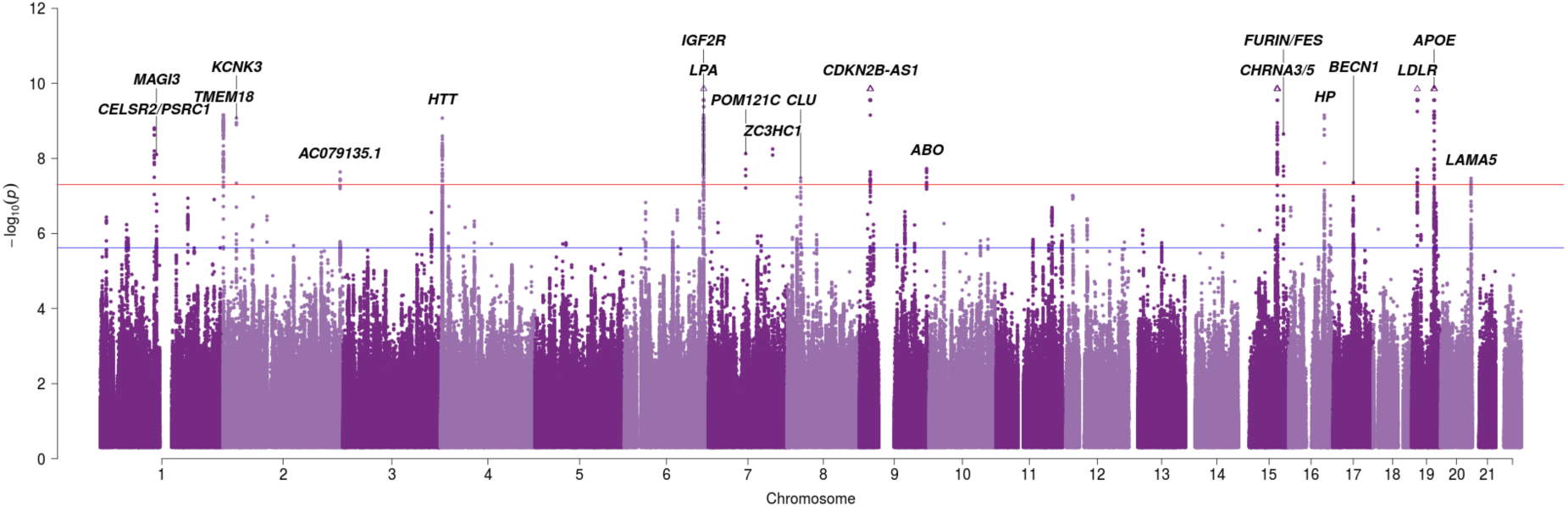
Manhattan plot of Bayesian associations of SNPs informed by risk factors with parental lifespan under the CES assumption. Bayesian iGWAS was performed using observed associations from the CES GWAS (discovery and replication sample combined) and priors based on 16 traits selected by an AIC-based stepwise model. As the P values were assigned empirically using a permutation approach, the minimum P value is limited by the number of permutations; SNPs reaching this limit are represented by triangles. Annotated are the gene, cluster of genes, or cytogenetic band in close proximity to the top SNP. The red line represents the genome-wide significance threshold (P = 5 x 10-8). The blue line represents the 1% FDR threshold.

Notably, 8 out of the 11 lead SNPs in discovery (for which we also had iGWAS data) show consistent evidence between replication and iGWAS (i.e. both weakened or strengthened the evidence together; Table 1).

We attempted to replicate the new hits and the rest of our genome-wide significant findings using publicly available summary statistics on extreme longevity (10, 15, 28), despite limited power. Remarkably, 23 out of 28 SNPs show directional consistency, and 9 SNPs or close proxies (r^2^ > 0.8) reach nominal significance in the replication sample (P < 0.05, one-sided test). Of these, SNPs near *ZC3HC1*, *ABO*, *GADD45G*, *LDLR*, *POM121C*, and *KCNK3* are replicated for the first time (Table S11), and thus appear to be lifespan and longevity SNPs. The overall, meta-analysed ratio of replication effect to discovery effect size – excluding *APOE*, which was predetermined as 1 to enable calibrations – is 0.37 (95% CI 0.26–0.48; P = 1.5×10^-11^), indicating that most of our lead lifespan SNPs are also longevity SNPs (i.e. the overall ratio is not zero; Fig. 4), but have an even greater effect on lifespan than longevity (relative to APOE).

**Fig. 4.**
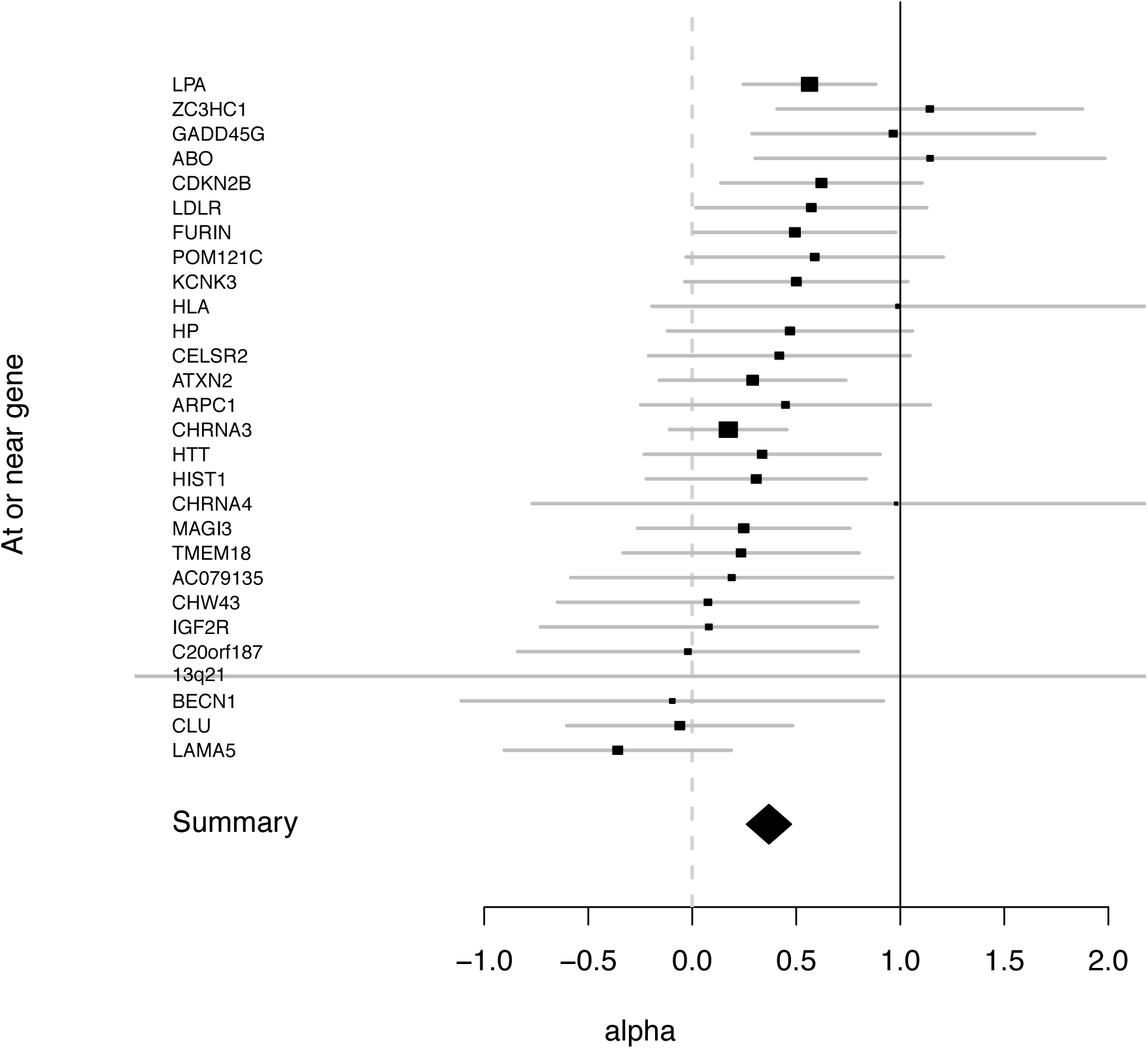
6 SNPs replicated for first time using 3 external GWAMAs of extreme long-livedness, whilst 23/28 show directional consistency. We attempted replicate the observed effect sizes for log hazard protection ratio in external GWAMAs of longevity (10, 15, 28),, having converted effect sizes to our scale (see Methods). At or near gene – the nearest gene to the lead SNP analysed (see Table S11). Alpha – ratio of replication to discovery effect sizes on the common scale and 95% CI (reflecting uncertainty in the numerator and denominator). A one-sided test was used for significance (nominal p<.05). A ratio of 1 indicates consistency with the relationship between the effect on 90+ longevity and lifetime hazard with that at APOE, a ratio of zero suggests no effect on replication. True (rather than estimated) alpha between 0 and 1 suggests the SNP has a greater effect on lifetime hazard than 90+ longevity, relative to APOE. SNPs where both 0 and 1 are covered are underpowered, although the result may be suggestive. APOE as the reference SNP (and thus, by definition, alpha=1) is excluded. The summary is the inverse variance meta-analysis of alpha over all SNPs 0.37 95% CI (0.26,0.48) p<1e-13 for H_0_ alpha <>0.

### Sex-and age-specific effects

We estimated SNP effects stratified by sex and age bands to identify age-and sex-specific effects. Although power was limited, as we sought contrasts in small effect sizes, we find 6 variants with age-specific effects on survival and 4 variants with sex-specific effects on survival (FDR 5% across the 49 putative lifespan and longevity variants considered). The lead variant at *APOE* shows stronger effects at older ages – the ε4 allele’s log hazard is about 2.5 times as strong in individuals in their 80s vs. 60s – whilst lead SNPs near *CLU*, *CHRNA3/5*, *ABO*, *CDKN2B-AS1*, and *EGLN2/CYP2A6*, show stronger effects at younger ages (Fig. 5a). Variants at or near 13q21.33 and *PSORS1C3* show stronger effects in women, while variants at or near *TOX* and *C20orf187* show stronger effects in men (Fig. 5b). Notably, the SNP at or near *ZW10*, which was identified by Pilling et al. (6) in men only and replicated by us in men, does not actually show statistically significant evidence of sex-specificity in our UK Biobank analysis (95% CI β_male_ –0.009 to 0.033; P = 0.266), although this could be due to lack of power (Table S20, Table S21).

**Fig. 5.**
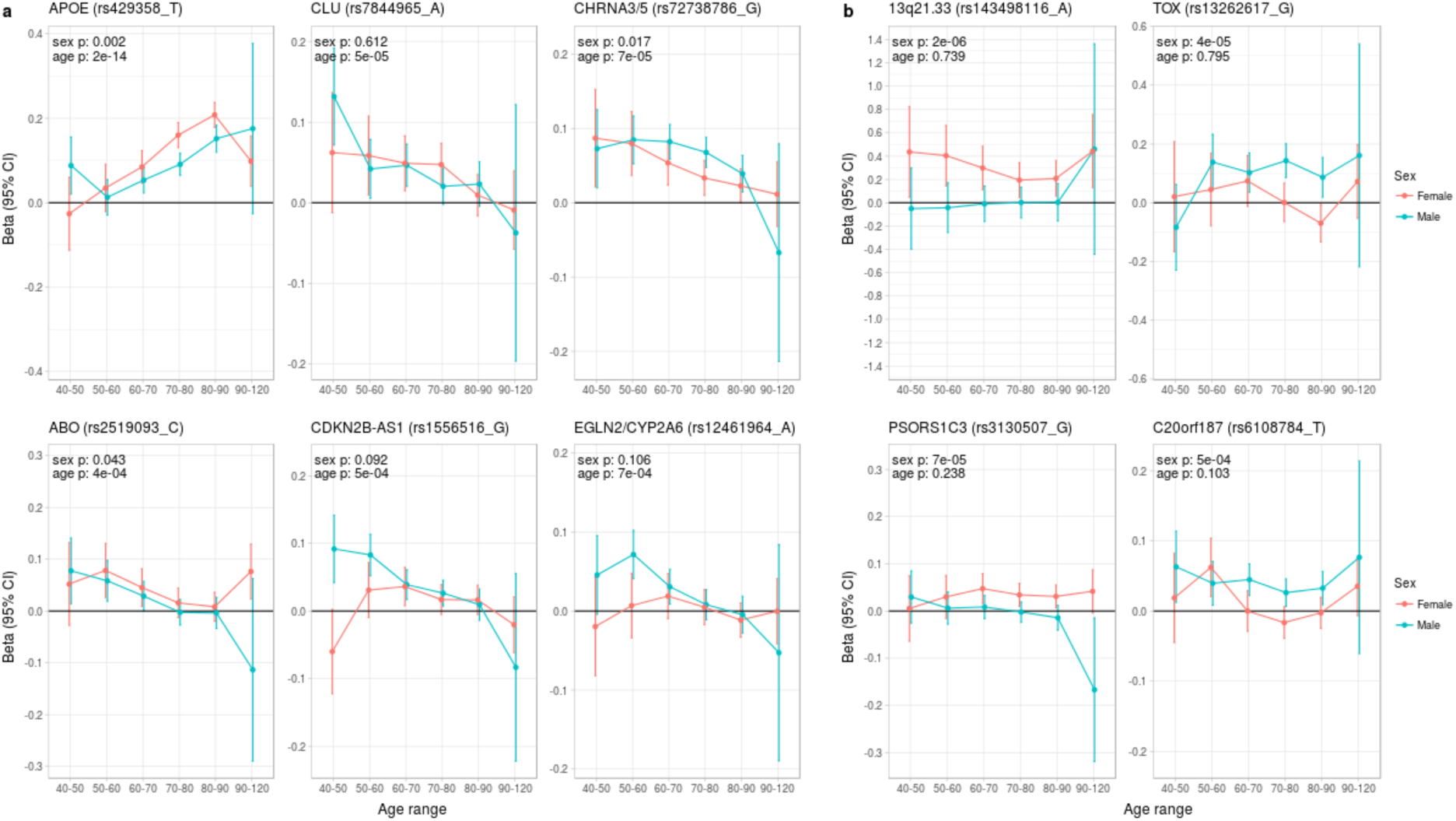
Age and sex specific effects on parent survival for 10 variants showing 5% FDR age-or sex-specificity of effect size from 49 lifespan-increasing variants. a) Variants showing age-specific effects; b) Variants showing sex-specific effects. Panel titles show the gene, cluster of genes, or cytogenetic band in close proximity to the lead lifespan variant, with this variant and lifespan-increasing allele in parentheses. Beta – log_e_(protection ratio) for 1 copy of effect allele in self in the age band (i.e. 2 x observed due to 50% kinship). Note the varying scale of y-axis across panels. Age range: the range of ages over which beta was estimated. Sex p – nominal P value for association of effect size with sex. Age p – nominal P value for association of effect size with age.

### Implication of causal genes and methylation sites

Combining gene expression and methylation data with our lifespan statistics, we identify causal roles for *FURIN* and *FES* within the *FURIN/FES* locus, *SH2B3* within the *ATXN2/BRAP* locus, and *SESN1* within the *FOXO3* locus. We also find causal CpG sites near *APOE*, *CHRNA3/5*, *HLA-DQA1*, *LPA*, *ATXN2/BRAP*, and 10 other loci at FDR 5%. (SI Appendix - section 1, Table S12, Table S13).

We performed conditional analysis on lead lifespan loci to find additional independent variants associated with lifespan. This increases out-of-sample predicted narrow-sense heritability by 79% (Table S14). As might be expected, *HLA-DQA1* appears to have high allelic heterogeneity, where at least 29 additional variants showed independent predictive causal effects, unable to be captured by only the top variant in the LD block.

### Disease and lifespan

We next sought to validate and understand how lifespan variants are linked to age-related disease by testing for disease association in UK Biobank and independently in PhenoScanner (25), recognizing in the former false positive associations with death might coincide with false positive associations with disease given the correlation between morbidity and mortality.

In UK Biobank, we find the lifespan-lengthening variants are protective against disease in 104 tests, but increase risk in 16 tests (at 5% FDR). Strikingly, the lifespan-lengthening variants are protective for cardiovascular disease (CVD) in 67 association tests and increased susceptibility only once (near *APOE*, although this SNP also shows two protective CVD associations, and is well known to be highly pleiotropic). For cancer, we see only 14 protective associations, all but four of which are related to lung cancer (SI Appendix - section 2, Table S6, Table S17). PhenoScanner associations are similar in character or even more pronounced (Table S7, Table S18).

Nonetheless, both analyses were subject to bias due to the structure of the sample as the numbers of disease cases (and thus power) differs by disease, a potential confounder, with cancer having been less studied and more heterogeneous than CVD. We therefore approached the question again, from the opposite end, identifying the most important loci for each disease category (neurological disease, CVD, diabetes, lung cancer, and other cancers) in large numbers (>20 associations in each category) from the GWAS catalog (29) and used our GWAS to see if the disease loci associate with lifespan. Our measure was lifespan variance explained (LVE, years^2^) by the locus, which balances effect size against frequency, and is proportional to selection response and the GWAS test statistic and thus monotonic for risk of false positive lifespan associations. Taking each independent disease variant, we ordered them by LVE, excluding any secondary disease where the locus was pleiotropic.

The Alzheimer’s disease locus *APOE* shows the largest LVE (0.23 years^2^), consistent with its most frequent discovery as a lifespan SNP in GWAS(3, 6, 7, 15). Of the 20 largest LVE SNPs, 12 and 4 associate with CVD and smoking/lung cancer, respectively, while only 2 associate with other cancers (near *ZW10* and *NRG1;* neither in the top 15 LVE SNPs). Cumulatively, the top 20/45 LVE SNPs explain 0.33/0.43 years^2^ through CVD, 0.13/0.15 years^2^ through smoking and lung cancer, and 0.03/0.11 years^2^ through other cancers (Fig. 6).

**Fig. 6:**
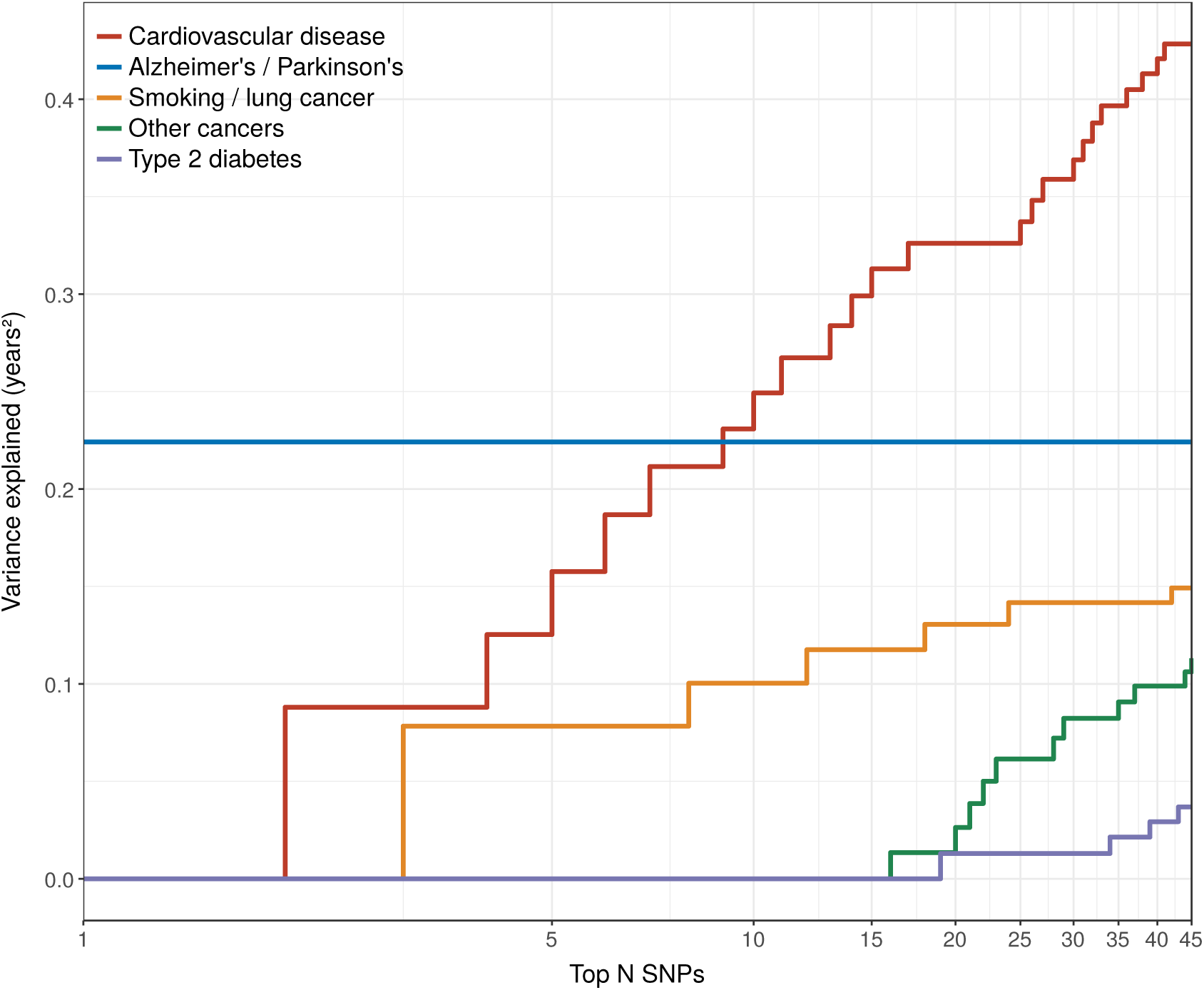
Disease loci explaining the most lifespan variance are primarily associated with neurological disease, cardiovascular disease, and lung cancer. SNPs reported as genome-wide significant for disease in European population studies, ordered by their lifespan variance explained (LVE), show the cumulative effect of disease SNPs on variation in lifespan. An FDR cut-off of 1.55% is applied simultaneously across all diseases, allowing for 1 false positive association with lifespan among the 45 independent loci. Note the log scale on the X axis. Cardiovascular disease – SNPs associated with cardiovascular disease or myocardial infarction. Alzheimer’s / Parkinson’s – SNPs associated with Alzheimer’s disease or Parkinson’s disease. Smoking / lung cancer – SNPs associated with smoking behaviour, chronic obstructive pulmonary disease and lung adenocarcinomas. Other cancers – SNPs associated with cancers other than lung cancer (see Table S19 for a full list). Type 2 diabetes – SNPs associated with type 2 diabetes.

Strikingly, two of the three largest LVE loci for non-lung cancers (at or near *ATXN2/BRAP* and *CDKN2B-AS1*), show **increased** cancer associating with **decreased** lifespan (due to antagonistic pleiotropy with CVD), while the third (at or near *MAGI3*) also shows evidence of pleiotropy, having an association with CVD three times as strong as breast cancer, and in the same direction. In addition, 6 out of the 11 remaining cancer-increasing/lifespan-decreasing loci passing FDR (near *ZW10*, *NRG1*, *C6orf106*, *HNF1A*, *C20orf187*, and *ABO*) also show significant associations with CVD but could not be tested for pleiotropy as we did not have data on the relative strength of association of every type of cancer against CVD, and thus (conservatively from the point of view of our conclusion) remain counted as cancer SNPs (Fig. 7, Table S19). Visual inspection also reveals an interesting pattern in the SNPs that did not pass FDR correction for affecting lifespan: cardio-protective variants associate almost exclusively with increased lifespan, while cancer-protective variants appear to associate with lifespan in either direction (grey dots often appear below the × axis for other cancers).

**Fig. 7.**
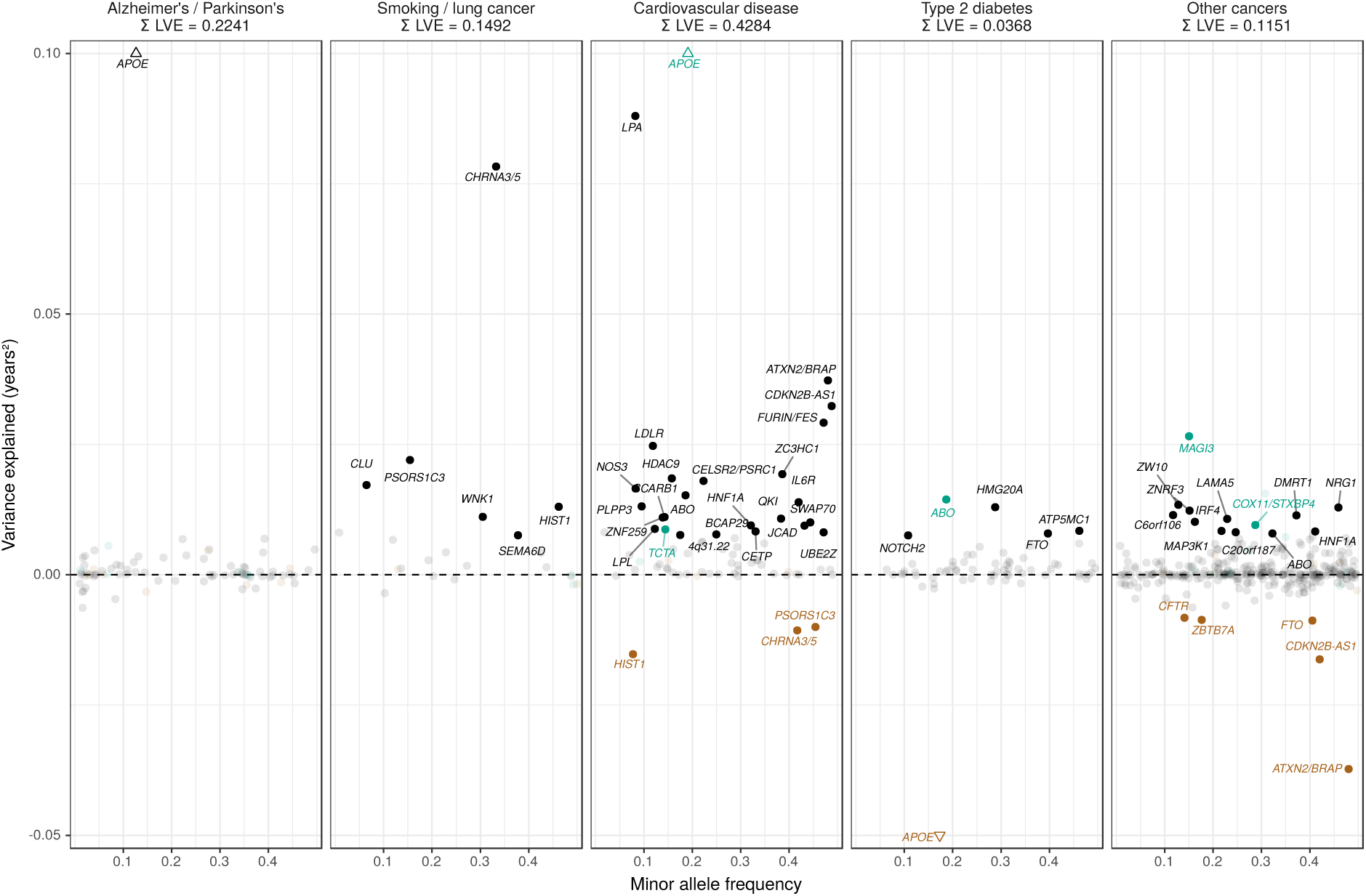
Lifespan variance explained by individual genome-wide significant disease SNPs within disease categories. Genome-wide significant disease SNPs from the GWAS catalog are plotted against the amount of lifespan variance explained (LVE), with disease-protective alleles signed positively when increasing lifespan and signed negatively when decreasing lifespan. SNPs with limited evidence of an effect on lifespan are greyed out: an FDR cut-off of 1.55% is applied simultaneously across all diseases, allowing for 1 false positive among all significant SNPs. Secondary pleiotropic SNPs (i.e. those associating more strongly with another one of the diseases, as assessed by PheWAS in UK Biobank) are coloured to indicate the main effect on increased lifespan seems to arise elsewhere. Of these, turquoise SNPs show one or more alternative disease associations in the same direction and at least twice as strong (double Z statistic – see Detailed Methods) as the principal disease, while brown SNPs show one or more significant associations with alternative disease in the opposite direction that explains the negative association of the disease-protective SNP with lifespan. Of specific interest is the SNP near MAGI3, which is reported as a breast cancer SNP but associates more strongly with CVD in UK Biobank and shows no evidence of sex-specific effects on lifespan. However, we do not classify it as a CVD SNP as its main effect on lifespan is likely due to protection from autoimmune disease by a nearby missense variant (rs6679677_C, r^2^>0.6, 95% CI log OR type 1 diabetes –0.74 to –0.46(30); rheumatoid arthritis –0.66 to –0.50(31), and carrying these diseases can reduce life expectancy up to 13 years (32, 33)). Similarly, the HLA-DQA1 locus also associates most strongly with autoimmune disease and is therefore absent from the analysis. The variance explained by all SNPs in black is summed (∑LVE) by disease. Annotated are the gene, cluster of genes, or cytogenetic band near the lead SNPs. The Y axis has been capped to aid legibility of SNPs with smaller LVE: SNPs near APOE pass this cap and are represented by triangles. Alzheimer’s/Parkinson’s – SNPs associated with Alzheimer’s disease or Parkinson’s disease. Smoking / lung cancer – SNPs associated with smoking behaviour, chronic obstructive pulmonary disease and lung adenocarcinomas. Cardiovascular disease – SNPs associated with cardiovascular disease or myocardial infarction. Type 2 diabetes – SNPs associated with type 2 diabetes. Other cancers – SNPs associated with cancers other than lung cancer (see Table S19). ∑LVE – Total Lifespan Variance Explained by non-pleiotropic SNPs passing FDR, in years^2^.

### Cell type and pathway enrichment

At FDR 5%, we find enrichment in SNP heritability in five categories: two histone and two chromatin marks linked to male and female foetal brain cells, and one histone mark linked to the dorsolateral prefrontal cortex of the brain. Despite testing other cell types, such as heart, liver, and immune cells, no other categories were statistically significant after multiple testing correction (Table S22).

Next, we determined which biological pathways could explain the associations between our genetic variants and lifespan, using three different methods. VEGAS highlights 33 gene sets at FDR 5%, but neither PASCAL nor DEPICT (with SNP thresholds at P < 5 × 10^−8^ and P < 1 × 10^−5^) finds any significant gene sets that passed multiple testing correction. The 33 gene sets highlighted by VEGAS are principally for blood lipid metabolism (21), with the majority involving lipoproteins (14) or homeostasis (4). Other noteworthy gene sets are neurological structure and function (5) and vesicle-mediated transport (3). Enrichment was also found for organic hydroxy compound transport, macromolecular complex remodelling, signalling events mediated by stem cell factor receptor (c-kit), and regulation of amyloid precursor protein catabolism (Table S23)

Finally, we performed an analysis to assess whether genes that have been shown to change their expression with age(34) are likely to have a causal effect on lifespan itself. Starting with a set of independent SNPs affecting gene expression (eQTLs), we created categories based on whether gene expression was age-dependent and whether the SNP was associated with lifespan in our study (at varying levels of significance).

We find eQTLs associated with lifespan are 1.78 to 3.45 times more likely to have age-dependent gene expression, dependent on the P value threshold used to define the set of lifespan SNPs (Table S24, Fig. S4).

### Out-of-sample lifespan predictions

We calculated polygenic risk scores (PRS) for lifespan for two subsamples of UK Biobank (Scottish individuals and a random selection of English/Welsh individuals), and one sample from the Estonian Biobank, using (recalculated) lifespan GWAS summary statistics that excluded these samples.

When including all independent markers, we find an increase of one standard deviation in PRS increases lifespan by 0.8 to 1.1 years, after doubling observed parent effect sizes to compensate for the imputation of their genotypes (see Table S25 for a comparison of performance of different PRS thresholds).

Correspondingly – again after doubling for parental imputation – we find a difference in median predicted survival for the top and bottom decile of PRS of 5.6/5.6 years for Scottish fathers/mothers, 6.4/4.8 for English & Welsh fathers/mothers and 3/2.8 for Estonian fathers/mothers. In the Estonian Biobank, where data is available for a wider range of subject ages (i.e. beyond median survival age) we find a contrast of 3.5/2.7 years in survival for male/female subjects, across the PRS tenth to first decile (Table 2, Fig. 8).

**Table 2:** Polygenic scores for lifespan are predictive of out-of-sample parent and subject lifespans. A polygenic risk score was made for each subject using GWAS results that did not include the subject sets under consideration. Subject or parent survival information (age entry, age exit, age of death, if applicable) was used to test the association between polygenic risk score and survival as (a) a continuous score and (b) by dichotomising the top and bottom decile scores. Sample – Population sample of test dataset, where E&W is England and Wales; Kin – individuals tested for association with polygenic score; N – Number of lives used for analysis; Deaths – Number of deaths; Beta – Effect size in log_e_(protection ratio), doubled in parents to reflect the expected effect in cohort subjects. SE – Standard error, doubled in parents to reflect the expected error in cohort subjects; Years – Estimated effect size in years of life; P – P value of two-sided test of association; Median age at death – difference in years between the median lifespan of individuals in the top decile of the score and the bottom decile (again raw observed parent effects are doubled).

| Sample | Kin | N | Deaths | Effect per longevity score SD |  |  |  | Contrast in top vs. bottom decile |  |  |  | Median age at death |  |
| --- | --- | --- | --- | --- | --- | --- | --- | --- | --- | --- | --- | --- | --- |
|  |  |  |  | Beta | SE | Years | P | Beta | SE | Years | P | Men | Women |
| Scotland | Parents | 46,936 | 33,196 | 0.107 | 0.011 | 1.1 | 4.2E-22 | 0.483 | 0.049 | 4.8 | 9.3E-23 | 5.6 | 5.6 |
| Scotland | Subjects | 24,059 | 941 | 0.085 | 0.033 | 0.8 | 1.0E-02 | 0.510 | 0.143 | 5.1 | 3.7E-04 | - | - |
| E&W | Parents | 58,070 | 39,347 | 0.133 | 0.010 | 1.3 | 7.3E-39 | 0.507 | 0.045 | 5.1 | 1.4E-29 | 6.4 | 4.8 |
| E&W | Subjects | 29,815 | 760 | 0.098 | 0.037 | 1.0 | 7.1E-03 | 0.067 | 0.161 | 0.7 | 6.8E-01 | - | - |
| Estonia | Parents | 61,728 | 29,660 | 0.099 | 0.012 | 1.0 | 2.5E-17 | 0.296 | 0.052 | 3.0 | 1.2E-08 | 3 | 2.8 |
| Estonia | Subjects | 24,800 | 2,894 | 0.087 | 0.019 | 0.9 | 2.6E-06 | 0.313 | 0.082 | 3.1 | 1.3E-04 | 3.5 | 2.7 |

**Fig. 8:**
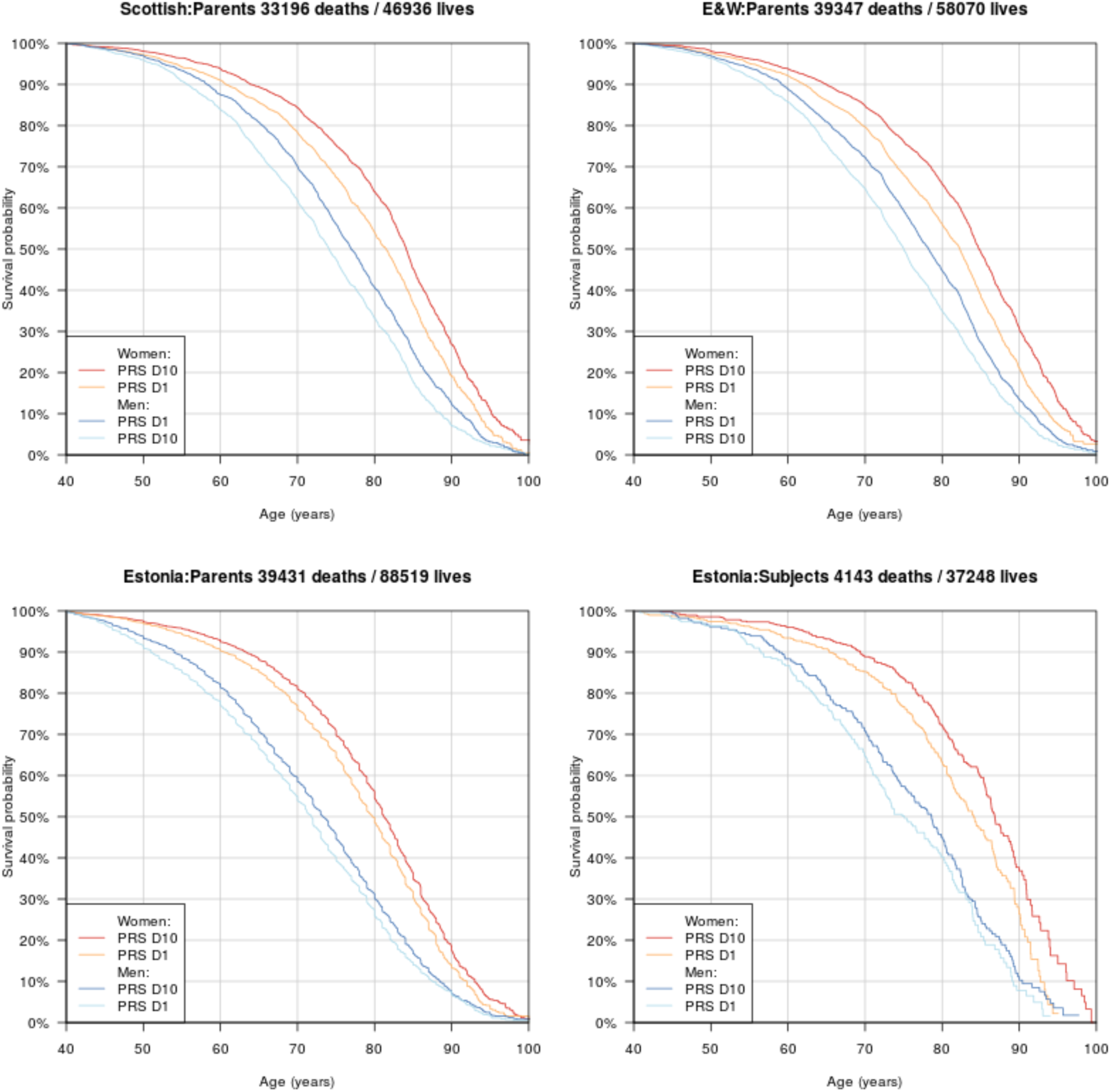
Survival curves for highest and lowest deciles of lifespan polygenic risk score. A polygenic risk score was made for each subject using GWAS results that did not include the subject sets under consideration. Subject or parent survival information (age entry, age exit, age of death (if applicable) was used to create Kaplan-Meier curves for the top and bottom decile of score. In this figure (only) no adjustment has been made for the dilution of observed effects due to parent imputation from cohort subjects. Effect sizes in parent, if parent genotypes had been used, are expected to be twice that shown. E&W – England and Wales; PRS – polygenic risk score.

Finally, as we did for individual variants, we looked at the age– and sex-specific nature of the PRS on parental lifespan and tested for associations with (self-reported) age-related diseases in subjects and their kin. We find a high PRS has a larger protective effect on lifespan for mothers than fathers in UK Biobank subsamples (P = 0.0071), and has a larger protective effect of lifespan in younger age bands (P = 0.0001) (Table S26,Fig. 9), although in both cases, it should be borne in mind that women and younger people have a lower baseline hazard, so a larger improvement in hazard ratio does not necessarily mean a larger absolute protection.

**Fig. 9.**
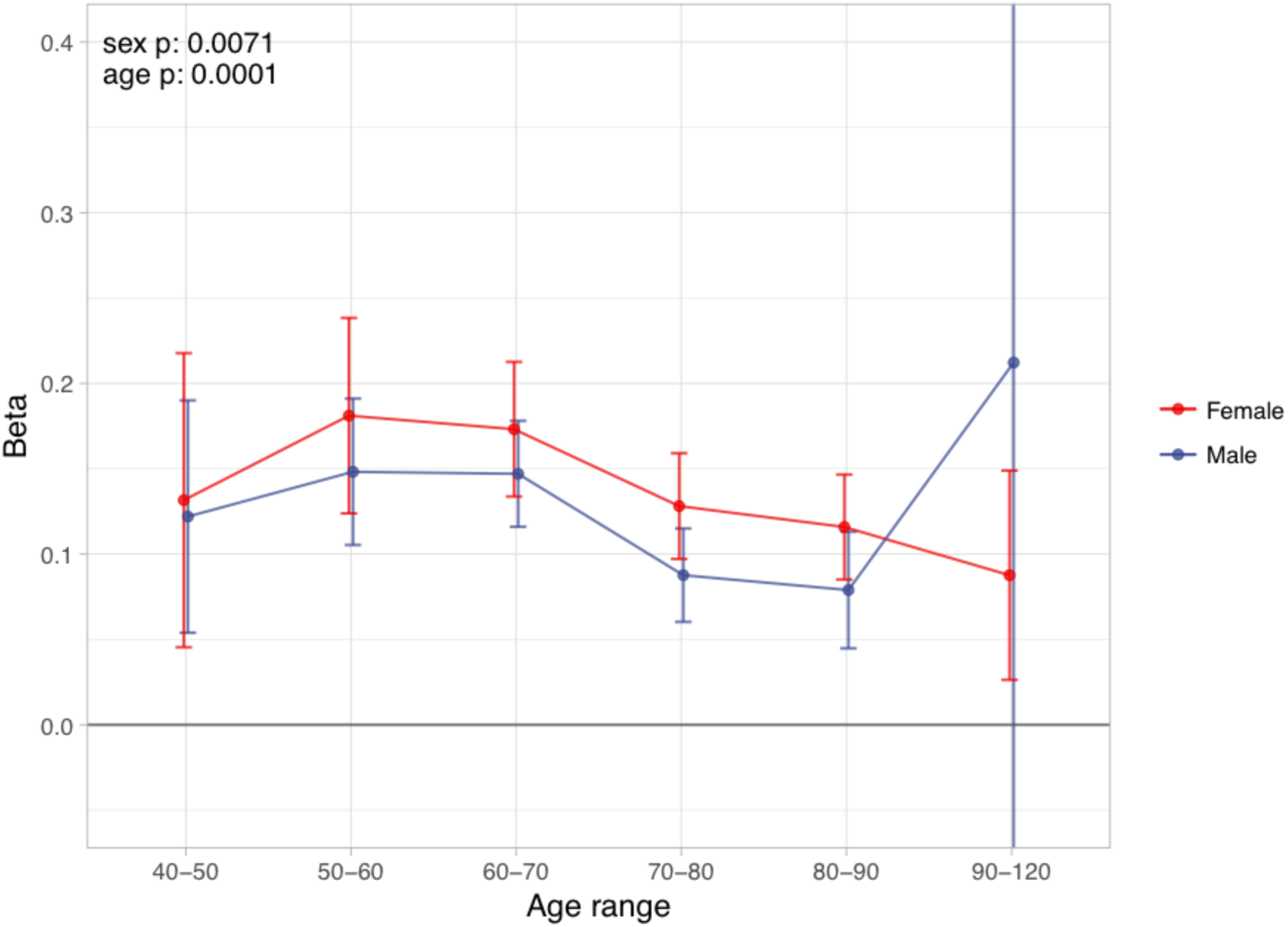
Sex and age specific effects of polygenic survival score (PRS) on parental lifespan in UK Biobank. The effect of out-of-sample PRS on parental lifespan stratified by sex and age was estimated for Scottish and English/Welsh subsamples individually (see Fig. S5) and subsequently meta-analysed. The estimate for the PRS on father lifespan in the highest age range has very wide confidence intervals (CI) due to the limited number of fathers surviving past 90 years of age. The beta 95% CI for this estimate is –0. 15 to 0.57. Beta – log_e_(protection ratio) for 1 standard deviation of PRS for increased lifespan in self in the age band (i.e. 2 x observed due to 50% kinship), bounds shown are 95% CI; Age range – the range of ages over which beta was estimated; sex p – P value for association of effect size with sex; age p – P value for association of effect size with age.

We find that overall, higher PRS scores (i.e. genetically longer life) are associated with less heart disease, diabetes, hypertension, respiratory disease and lung cancer, but increased prevalence of Alzheimer’s disease, Parkinson’s disease, prostate cancer and breast cancer, the last three primarily in parents. We find no association between the score and prevalence of cancer in subjects. (Table S27, Fig. S6).

## Discussion

Applying the kin-cohort method in a GWAS across UK Biobank and the LifeGen consortium, we replicated associations with lifespan for 6 loci discovered previously and 6 discovered here. We identified a further 12 novel lifespan variants at genome-wide significance, without replication. Further discussion of the loci, their mechanism, diseases and power are contained in SI Appendix - section 3.

Genetic variants affecting lifespan were enriched for pathways involving the transport, homeostasis and metabolism of lipoprotein particles, validating previous reports(35). We also identified new pathways including vesicle transport, metabolism of acylglycerol and sterols, and synaptic and dendritic function. We discover histone marks associated with foetal brain cells and the adult dorsolateral prefrontal cortex are enriched for human lifespan heritability.

We described multiple disease associations of life-lengthening variants and whole-genome polygenic risk scores with protection from cardiovascular disease, diabetes, COPD, and lung cancer, but found very few associations with other forms of cancer, suggesting cancer causes death more through (perhaps many) rarer variants or the enviroment. Finally, we showed that, using our GWAS results, we can construct a polygenic risk score making 3 to 5 year distinctions in life expectancy at birth between individuals from the score’s top and bottom decile.

Despite replicating long-standing longevity SNPs near *APOE*, *FOXO3*, and 5q33.3/EBF1 – albeit with smaller effect sizes in the latter two cases – we do not find evidence of association with lifespan for more recently published longevity SNPs near *IL6*, *ANKRD20A9P*, *USP42*, and *TMTC2*. Although this could be due to the original findings arising from chance, differences in ancestry and sample structure might also be the cause, but more intriguingly there remains the possibility that we were unable to replicate the effects because they uniquely act through mechanisms slowing the ageing process which only become apparent in the extremely long-lived.

At the same time, our analysis comparing lifespan and longevity effect sizes suggests that lifespan SNPs often associate with extreme long-livedness, consistent with the genetic correlation between the two traits (r_g_ = 0.73; SE = 0.11(35)). Again, the remaining 27% clearly leaves room for SNPs affecting lifespan and longevity in distinct ways.

Much work has been done implicating *FOXO3* as an aging gene in model organisms(36, 37), however we found the association in humans at that locus may be driven by expression of *SESN1* (admittedly a finding restricted to peripheral blood tissue). *SESN1* is a gene connected to the *FOXO3* promoter via chromatin interactions and involved in the response to reactive oxygen species and mTORC1 inhibition(38). This contrasts with fine-mapping studies which found common genetic variation within the locus increases expression of *FOXO3* itself (39, 40)

The magnitude of the distinctions our genetic lifespan score is able to make (5 years of life between top and bottom decile) is meaningful socially and actuarially: the implied distinction in price (14%; Methods) being greater than some recently reported annuity profit margins (8.9%)(41). However, the legal and ethical frameworks (at least in the UK(42)) surrounding genomic testing and commercial applications, such as life insurance, have yet to regulate the use of genome-wide measures (rather than single markers), especially for annuities. This needs to be urgently addressed.

Although clearly meaningful in isolation, our lifespan predictions may only have practical clinical or actuarial meaning if, rather than at birth, distinctions in lifespan can be drawn in middle age, and include independent information beyond that readily available using existing risk measures (e.g. occupation, smoking and blood pressure) in middle age. Such an assessment has been beyond the scope of this work; in part as such risk measures are not readily available for the parents (rather than offspring) studied.

The analysis of parent lifespans has enabled us to probe mortality for a generation whose lives have often been complete and attain unprecedented power in a survival GWAS, but changes in the environment (and thus the relative importance of each genetic susceptibility, for example following the smoking ban) inevitably mean we have less certainty about associations with prospective lifespans for the present generation of middle aged people, or the next.

The diseases which we found were associated with lifespan SNPs mirror the causal effect estimates from mortality risk factors(35) and are some of the leading causes of death across the world(43). However, importantly, the identified effects of lifespan-associated SNPs are not simply those of the risk factors. Surprisingly, although cancer is a major source of mortality, common genetic variation associated with increased lifespan does not appear to arise from a protection from cancers (other than lung cancer mediated through smoking), despite reported heritabilities for myocardial infarction and cancers being similar: around 30%(44, 45) All this suggests genetic variation for non-lung cancer alleles affecting lifespan is either rarer (e.g. BRCA1(46)), has smaller effects, or exhibits antagonistic pleiotropic effects, either due to linkage or biological compromise. Curiously, we find little evidence of SNPs of large deleterious effect on lifespan acting with antagonistic pleiotropy on other fitness and developmental component traits, despite long-standing theoretical suggestions to the contrary (47). However, we did not examine mortality before the age of 40, or mortality of individuals without offspring (by definition as we were examining parental lifespans), which may well have exhibited this feature.

In conclusion, recent genomic susceptibility to death in the normal age range seems rooted in modern diseases: Alzheimer’s, lung cancer and CVD; in turn arising from our modern – long-lived, obesogenic and tobacco-laden – environment, however the keys to the distinct traits of aging and extreme longevity remain elusive. At the same time, genomic information alone can now make material predictions of variations in expected length of life, although the accuracy of the predictions is far from supporting genetic determinism of that most (self-) interesting of traits - your lifespan.

## Methods - Summary

### GWAS

For each European ethnicity in UK Biobank, association analysis was performed between unrelated subjects’ genotypes (MAF > 0.005; HRC imputed SNPs only; ~9 million markers) and parent survival using age and alive/dead status in residualised Cox models, as described in (5) (5). To account for parental genotype imputation, effect sizes were doubled, yielding log hazard ratios for the allele in carriers themselves. These values were inverted to obtain a measure of log protection ratio, where higher values indicate longer life.

Mother and father survival information was combined in two separate ways, essentially assuming the effects were the same in men and women (common effects between sexes; CES), or allowing for sex-specific effect sizes (potentially different effects between sexes; PDES), with appropriate allowance for the covariance amongst the traits. For the second analysis we used MANOVA, implemented in MultiABEL (48). (48)

For LifeGen, where individual-level data was not available, parent survival summary statistics were combined for CES using conventional fixed-effects meta-analysis, adjusted to account for the correlation between survival traits (estimated from summary-level data). For PDES, the same procedure was followed as for the UK Biobank samples, with correlation between traits again estimated from summary-level data.

CES discovery and replication statistics were combined with inverse-variance meta-analysis. Both the discovery GWAS and combined cohort GWAS showed acceptable inflation, as measured by their LD-score regression intercept (<1.06, Table S4).

### Candidate SNP replication

Effect sizes from longevity studies were converted to our scale using an empirical conversion factor, based on the observed relationships between longevity and hazard ratio at the most significant variant at or near APOE, observed in the candidate SNPs study and our data (5).

Estimates reported in Pilling et al. (6) were based on rank-normalized Martingale residuals, unadjusted for the proportion dead, which – for individual parents – could be converted to our scale by multiplying by sqrt(c)/c, where c is the proportion dead in the original study (see Detailed Methods for derivation). Combined parent estimates were converted using the same method as the one used for longevity studies.

The deletion reported by Ben-Avrahim et al. (26) is perfectly tagged by a SNP that we used to assess replication. Assuming a recessive effect and parental imputation, we derived the expected additive effect to be 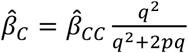, where *β̂*_*c*_ is the additive effect we expect to observe, *β̂*_*cc*_ is the homozygous effect reported in the original study, *q* is the C allele frequency, and *p* is 1 – *q*. (see Detailed Methods for derivation)

### iGWAS

58 GWAS on mortality risk factors were used to create Bayesian priors for the SNP effects observed in the combined cohort CES study, as described in (35). In short, Mendelian randomisation was used to estimate causal effects of independent risk factors on lifespan, and these estimates were combined with the risk factor GWAS to calculate priors for each SNP. Priors were multiplied with observed Z statistics and used to generate Bayes factors. Observed Z statistics were then permuted, leading to 7.2 billion null Bayes factors (using the same priors), which were used to assess significance.

### Sex and age stratified analysis

Cox survival models, adjusting for the same covariates as the standard GWAS, were used to test SNP dosage against father and mother survival separately. The analysis was split into age bands, where any parent who died at an age younger than the age band was excluded and any parent who died beyond the age band was treated as alive. Using the R package “metafor", moderator effects of sex and age on hazard ratio could be estimated while taking into account the estimate uncertainty (see Detailed Methods for formula).

### Causal genes and methylation sites

SMR-HEIDI (49) tests were performed on our combined cohort CES statistics to implicate causal genes and methylation sites. Summary-level data from two studies on gene expression in blood (50, 51) and data on gene expression in 48 tissues from the GTEx consortium (52) were tested to find causal links between gene expression and lifespan. Similarly, data from a genome-wide methylation study (53) was used to find causal links between CpG sites and lifespan. All results from the SMR test passing a 5% FDR threshold where the HEIDI test P > 0. 05 were reported.

### Conditional analysis

SOJO (54) was used to fine-map the genetic signals in 1 Mb regions around each top SNP identified in the discovery GWAS, combined cohort GWAS, and iGWAS. The analysis was based on UK Biobank CES discovery statistics, using the CES replication cohort to optimise the LASSO regression tuning parameters. For each parameter, a polygenic score was built and the proportion of predictable variance from the regional polygenic score in the validation sample was calculated.

### Disease-wide association analysis

Logistic regression, adjusted for subject sex; subject age; genotyping batch and array; and 40 principal components, was used to test diseases against SNP dosages from lead SNPs from our discovery and combined cohort GWASs, and external survival GWAS. Diseases were self-reported by 325,292 unrelated, genomically British subjects from UK Biobank about themselves, their siblings, and each parent. Before analysis, subject diseases were grouped to match pre-existing disease categories for family members (see Table S16 for grouping). Results were corrected for multiple testing using Benjamini-Hochberg with an FDR threshold of 5% and then further grouped into broad disease categories.

PhenoScanner was used to look up known associations with the same SNPs and close proxies (r^2^ > 0.8) (25). All associations passing a 5% FDR threshold were divided using keywords into broad disease categories, which were then further curated (see Table S18 and Detailed Methods for grouping criteria).

### Lifespan variance explained by disease SNPs

The GWAS catalog was checked for disease associations discovered in European ancestry studies, which were grouped into broad disease categories based on keywords and manual curation (see Table S19 and Detailed Methods). Associations were pruned by distance (500kb) and LD (r^2^ < 0.1), keeping the SNP most strongly associated with lifespan in the combined cohort CES GWAS. Where possible this SNP was tested against diseases in UK Biobank subjects and their family, as described above, to test for pleiotropy. Significance of associations with lifespan was determined by setting an FDR threshold that allowed for 1 false positive among all independent SNPs tested (q ≤ 0.022). Lifespan variance explained (LVE) was calculated as 2pqa^2^, where p and q are the frequencies of the effect and reference alleles in our lifespan GWAS, and a is the SNP effect size in years of life (55).

### Cell type enrichment

Stratified LD-score regression (56) was used to test for cell type-specific enrichment in lifespan heritability in the CES discovery cohort GWAS statistics, which had the highest SNP heritability as measured by LD-score regression (57). The statistics were analysed using the procedure described in (56), and P values were adjusted for multiple testing using the Benjamin-Hochberg procedure.

### Pathway enrichment

VEGAS2 v2.01.17 (58) was used to calculate gene scores using SNPs genotyped in UK Biobank, based on summary statistics of the combined cohort CES GWAS and the default software settings. VEGAS2Pathway was then used to check for pathway enrichment using those gene scores and the default list of gene sets (59).

DEPICT (60) was also used to map genes to lifespan loci and check for pathway enrichment in the combined cohort CES GWAS. Default analysis was run for regions with genome-wide significant (P < 5e-8) variants in the first analysis, and genome-wide suggestive (P<1e-5) variants in the second analysis, excluding the MHC in both cases.

PASCAL (61) was used with the same summary statistics and gene sets as DEPICT, except the gene probabilities within the sets were dichotomized (Z>3) as described in (62).

For each software independently, pathway enrichment was adjusted for multiple testing using the Benjamin-Hochberg procedure.

### Age-related eQTL enrichment

Combined cohort CES lifespan statistics were matched to eQTLs associated with the expression of at least one gene (P<1e-4) in a dataset provided to us by the eQTLGen Consortium (14,155 individuals). Data on age-related expression (34) allowed eQTLs to be divided into 4 categories based on association with age and/or lifespan. Fisher’s exact test was used check if age-related eQTLs were enriched for associations with lifespan.

### Polygenic score analysis

Polygenic risk scores (PRS) for lifespan were calculated for two subsamples of UK Biobank (24,059 Scottish individuals and a random 29,815 English/Welsh individuals), and 36,499 individuals from the Estonian Biobank, using combined cohort CES lifespan summary statistics that excluded these samples. PRSice 2.0.14.beta (63) was used to construct the scores from genotyped SNPs in UK Biobank and imputed data from the Estonian Biobank, pruned by LD (r^2^ = 0.1) and distance (250kb). Polygenic scores were Z standardised.

Cox proportional hazard models were used to fit parental survival against polygenic score, adjusted for subject sex; assessment centre; genotyping batch and array; and 10 principal components. Parental hazard ratios were converted into subject years of life as described in the GWAS method section.

Logistic regression models were used to fit polygenic score against the same self-reported UK Biobank disease categories used for individual SNPs. Effect estimates of first degree relatives were doubled to account for imputation of genotypes and then meta-analysed using inverse variance weighting, adjusting for trait correlations between family members.

## Data availability

The results that support our findings, in particular, the GWAS summary statistics for >1 million parental lifespans in this study are available to bona-fide researchers from the corresponding author upon request or from UK Biobank(5, 34, 64, 65) eQTLGen Consortium results will be made available after that manuscript is published.

## Methods - Details

### Data sources

Our discovery cohort consisted of 409,700 genomically British individuals from UK Biobank. Details on genotyping marker and sample QC are described in (23). Subjects completed a questionnaire which included questions on adoption status, parental age, and parental deaths. For our analysis, we excluded individuals who were adopted or otherwise unclear about their adoption status (N = 5,829), individuals who did not report their parental ages (N = 2,650), and individuals both of whose parents died before the age of 40 and which were therefore more likely due to accident or injury (N = 3,927). We further excluded one of each pair from related individuals (N = 71,288) from every relative pair reported by UK Biobank, leaving 326,006 individuals for the final analysis. Although exclusion of relatives reduces sample size, we were concerned that linear mixed modelling to account for relatedness might not be fully appropriate under the kin-cohort model. Consider the parental phenotypic correlation for two full sibling subjects (r^2^ = 1) or the maternal genetic covariance amongst two subjects who are the offspring of two brothers (r^2^ = 0): the heritability/GRM implied covariance if incorrect for both cases (although in the sibling case, it may be correct on average). Individuals passing QC reported a total of 312,260 paternal and 322,945 maternal lifespans, ranging from 40 to 107 years of life, i.e. 635,205 lives in total (Table S1).

Our replication dataset was LifeGen, a consortium of 26 population cohorts investigating genomic effects on parental lifespans(5). LifeGen had included results from UK Biobank, but the UK Biobank GWAS data were removed here, giving GWAS summary statistics for 160,461 father and 160,158 mother lifespans in the form of log hazard ratios. This dataset was then supplemented by 56,416 parental lifespans of UK Biobank individuals of self-reported British (but not identified as genomically British), Irish, and other white European descent, not included in the discovery. Cohorts were combined using inverse-variance meta-analysis, giving a total replication set of 377,035 lifespans, and over 1 million lives across discovery and replication combined.

### UK Biobank Genome-Wide Association Study

In the discovery and UK Biobank portions of the replication cohorts separately for each self-declared ethnicity, we carried out association analysis between genotype (MAF > 0.005; HRC imputed SNPs only; ~9 million markers) and parent age and alive/dead status, effectively analysing the effect of genotype in offspring on parent survival, given survival to at least age 40, using Cox Proportional Hazards models. The following model was assumed to hold:

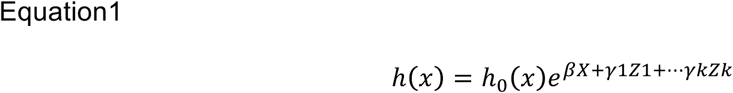

Where × is (parent) age, h0 the baseline hazard and X the offspring genotype(coded 0,1,2), beta the log_e_(hazard ratio) associated with X and Z_1_-Z_k_ the covariates, with corresponding effect sizes y_1_-y_k_. The covariates were genotyping batch and array, the first forty principal components of relatedness, as provided by UK Biobank, and subject sex (but not age, as we were analysing parent age).

To facilitate practical runtimes, the Martingale residuals of the Cox model were calculated for father and mothers separately and multiplied by 1/proportion dead to give estimates of the hazard ratio(66) giving a residual trait suitable for GWAS (for more details of the residual method see Joshi et al.(5)). Effect sizes observed under this model, for a SNP in offsprining are half that of the actual effect size in the parent carrying the variant(3). Reported effect sizes (and their SE) have therefore been doubled to give the effect sizes in carriers themselves, giving an estimate of the log hazard ratios (or often, with sign reversed, log protective ratios). These estimates are suitable for meta-analysis and allow direct comparison with the log hazard ratios from LifeGen.

Analysis of association between genotype and survival across both parents was made under two contrasting assumptions and associated models, which had to adjust for the covariance amongst parent traits, preventing conventional unadjusted inverse-variance meta-analysis. Firstly, we assumed that the hazard ratio was the same for both sexes, i.e. a common effect size across sexes (CES). If there were no correlation amongst parents’ traits, this could have been done by straightforward inverse variance meta-analysis of the single parent results. However, to account for the covariance amongst father and mother lifespans, we calculated a total parent residual, the sum of individual parent residuals, for each subject (i.e. offspring). Under the common effect assumption, the combined trait’s effect size is twice that in the single parent, and the variance of the combined trait, automatically and appropriately reflects the parents’ covariance, amongst the two parents, giving a residual trait suitable for GWAS, with an effect size **equal** to that in a carrier of the variant, and correct standard error. Secondly, we assumed that, there might be potentially different effect sizes across sexes (PDES) in fathers and mothers. Under the PDES assumption, individual parental GWAS were carried out, and the summary statistics results were meta-analysed using MANOVA, accounting for the correlation amongst the parent traits and the sample overlap (broadly complete), but agnostic as to whether the effect size was similar or different in each parent, giving a P value against the null hypothesis that both effect sizes are zero, but, naturally, no estimate of a single common beta. This procedure was carried out using the R package MultiABEL(48) and used summary-level data for the analysis(67). The procedure requires an estimate of the correlation amongst the traits (in this case parent residuals), which was measured directly (r = 0.1). The procedure automatically estimates the variance of the traits from summary level data (Mother residuals σ^2^ = 6.74; Father residuals σ^2^ = 5.25)

For the replication cohort, the PDES procedure to combine results was identical to discovery (Mother residuals σ^2^ = 14.12; Father residuals σ^2^ = 18.75), except the trait correlation was derived from summary level data instead of measured directly (r = 0.1). This was done by taking the correlations in effect estimates from independent SNPs from the summary statistics of the individual parents, which equals the trait correlation, assuming full sample overlap (which is slightly conservative). Similarly, since we did not have access to individual level (residual) data, it was not possible to carry out a single total parent residual GWAS under the CES assumption. Instead we meta-analysed the single parent effect sizes using inverse variance meta-analysis, but adjusted the standard errors to reflect the correlation amongst the traits (r) as follows:

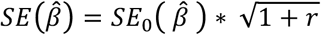

Where *SE*_0_(*β̂*) is the usual (uncorrected) inverse-variance weighted meta-analysis standard error, ignoring the correlation amongst the estimates and *SE*(*β̂*) is the corrected estimate used.

This is slightly conservative as

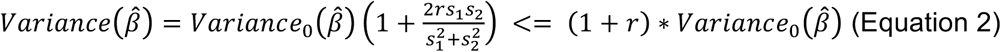

which follows straightforwardly from 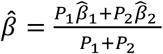.

Where *s*_1_ and *s*_2_ are the standard error of the individual estimates and *P*_1_, *P*_2_ their associated precisions (i.e. reciprocal of the variance). Equation (2) is always conservative, but exact if *s*_1_ = *s*_2_. In practice *s*_1_ and *s*_2_ were similar, as the sample sizes, allele frequencies and variance in the traits for the two parents were very similar.

As we were using unrelated populations and fitting forty principal components to control for population structure, material inflation of test statistics due to structure or relatedness was not to be expected. This was confirmed using the intercept of LD-score regression(57) for the summary statistics as shown in Table S4. We have tried to use a consistent approach to the direction of lifespan altering effects: positive implies longer life, consistent with previous studies of long-livedness(15). Our base measure was thus a protection ratio, directly mirroring the cox hazard ratio. Effect sizes (betas) are typically –log_e_(cox hazard ratio), which we denote the log_e_(protection ratio). Years of life gained were estimated as 10 * log protection ratio, in accordance with a long-standing actuarial rule of thumb and recently verified(5).

### Candidate SNP replication

We sought to reproduce and replicate genome-wide significant associations reported by Pilling et al.(6), who recently published a GWAS on the same UK Biobank data, but using a slightly different method. Rather than excluding relatives, Pilling et al. used BOLT-LMM and the genomic relationship matrix in subjects, to approximately account for covariance amongst parental phenotype. Pilling at al. also analysed parents separately as well as jointly, using a last survivor phenotype. Despite these factors, reproduction (obtaining the same result from almost the same data) was straightforward and consistent, once effect sizes were placed on the same scale (see below and **Fig. S1**), confirming our re-scaling was correct. To try to independently replicate their results, we used the consortium, LifeGen, excluding individuals from UK Biobank.

Pilling et al.(6) performed multiple parental survival GWAS in UK Biobank, identifying 14 loci using combined parent lifespan and 11 loci using individual parent lifespan. Their study design involved rank-normalising Martingale residuals before regressing against genotype, which does not give an estimate of the loge(hazard ratio), nor, we believe, another naturally interpretable scale of effects, as the scale is now dependent on the proportion dead. Simulations (not shown) suggested 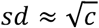 for some Martingale residual distributions, where sd is the standard deviation of the distribution and c is the proportion dead. As multiplying the untransformed Martingale residual distribution by 1/c gives an estimate of the hazard ratio (5, 66)), for individual parents, we could convert Pilling et al’s effect sizes by multiplying them by sqrt(c) to return them to the Martingale residual scale (which still depends on the study structure) and then by 1/c to place them on the log HR scale, using the proportion dead from Pilling et al.’s study descriptives. Further multiplication by 2 allows conversion from a subject-parent effect to an effect in self. The cumulative scale conversion allowing for all three of these effects was to multiply Pilling et al’s effect sizes by 2.5863/2.2869 in mothers/fathers, respectively, placing them on a log_e_ HR scale for effects in male/female subjects. The joint life parent phenotype does not appear to have a straightforward conversion to log_e_ HR in self. Instead, we used an empirical estimate derived from effect sizes comparison of the APOE allele between Pilling’s discovery sample and our own UK Biobank Gen. British discovery sample (both parents combined), which were largely overlapping: to get from Pilling et al.’s effect size to log_e_ HR, we had to multiply their effect sizes by 1.9699 for APOE and used this ratio for other alleles, which should be completely valid under the proportional hazard assumption. Whilst this scheme may appear a little *ad hoc* (the use of simulation and APOE), it was confirmed empirically: visual inspection indeed showed hazard ratios from our own UK Biobank discovery sample calculations and inferred hazard ratios from Pilling were highly concordant (**Fig. S1**, noting one concordance – for joint life at APOE, which was pre-defined to be perfectly concordant by our procedure, is not, of itself, evidence).

Flachsbart et al.(13) and Deelen et al.(15) tested extreme longevity cases (95–110 years, ≥85 years, respectively) against controls (60–75 years, 65 years, respectively), identifying SNPs at or near *FOXO3* and 5q33.3/EBF1. As done previously(5), we estimated the relationship between longevity log odds ratio and log hazard ratio empirically using APOE variant rs4420638_G (reported log OR –0.33 (15), our log HR –0.086), assuming increased odds of surviving to extreme age is due to a reduction in lifetime mortality risk. Inverting the sign to give log_e_(protection ratio) estimates, the conversion estimate used was –3.82.

Ben-Avrahim et al.(26) reported a deletion in Growth Hormone Receptor exon 3 (*d3-GHR*) associated with an increase of 10 years in male lifespan when homozygous. This deletion is tagged by rs6873545_C(68), which is present in our combined discovery and replication sample at a frequency of 26.9% (*q*). Considering the association is recessive and we are imputing father genotypes, we converted the reported effect size into expected years of life per allele as follows:

If the subject genotype is CT, the parent contributing the C allele has 50% chance of being the father and 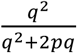 chance of being homozygous. If the subject genotype is CC, the father has 100% chance of contributing the C allele and again has 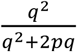 chance of being homozygous. We therefore expect the relationship to be 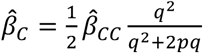, where *β̂*_*c*_ is the observed effect per subject allele on father lifespan and *β̂*_*CC*_ is the reported effect of the homozygous deletion in the father. As before, doubling the allele effect gives an estimate of the effect of the allele on subject lifespan, which finally yielded a converted estimate of 0.155: i. e. under Ben-Avrahim et al.’s assumptions on inheritance patterns, if their estimate of effect size in minor homozygotes is correct, we should see under the additive model an effect size of 0.155 years, or a loge(hazard ratio) of –0.015, and correspondingly scaled standard errors (note we are assuming that the effect is actually recessive, and estimating how that effect should appear if an additive model is fitted).

Standard errors were calculated from inferred betas and reported P values, assuming a twosided test with a normally distributed estimator. Confidence interval overlap was then assessed using a two-sided test on the estimate difference (P_diff_), using a Z statistic from the difference divided by the standard error of the difference.

### iGWAS

We performed a Bayesian Genome-Wide Association Study using the CES discovery-replication meta-analysis results and summary statistics on 58 risk factor GWASs (imputed, leading to 7.2 million SNPs in common between all the studies), as described by McDaid et al. (35). To calculate our prior for SNPs on a given chromosome, first we used a multivariate Mendelian Randomization (masking the focal chromosome) to identify the risk factors significantly influencing lifespan and estimate their causal effect. This identified 16 risk factors independently causally contributing to lifespan (see Table S9 for the causal effect estimates). Next, assuming that a SNP affects lifespan through its effects on the 16 risk factors, prior effects estimates were estimated as the sum of the products of the causal effect estimates of the 16 significant risk factors on lifespan and the effect of the SNP on each risk factor. We added 1 to the prior effect variance formula described in McDaid et al. (35) to account for the fact that prior effects are estimated using observed Z-scores, and not true Z-scores, with *Z*_*obs*_~***N***(*Z*_*true*_, 1).

We computed Bayes factors by combining the prior effects and the observed association Z statistics. The significance of the Bayes factors was assessed using a permutation approach to calculate P values, by comparing observed Bayes factors to 7.2 billion null Bayes factors. These null Bayes factors were estimated using 1000 null sets of Z statistics combined with the same priors. These empirical P values were then adjusted for multiple testing using the Benjamini-Hochberg procedure.

### Replication in extreme long-livedness

Published summary statistics for 3 GWAMAs of extreme long-livedness were used from Deelen *et al*. (age > 90)(15), Broer *et al*. (10), Walker *et al*. (28). Effect sizes were given (or could be estimated from P value, effect direction and N, as well as the SNPs MAF). These effect sizes were rescaled to hazard ratios, using the effect size in the GWAMA concerned at the reference SNP (rs2075650 near APOE), compared to the hazard ratio in our own GWAS, giving an assumed hazard ratio of 0.0822 in all GWAMA at rs2075650 and proportionate effect sizes at all other SNPs. This method assumes a stable relationship between the lifespan hazard ratio and effect on longevity, as is true under the proportionate hazards assumption (5) Having recalibrated the 3 published GWAMAs to a common scale, effect sizes were meta-analysed using fixed-effect inverse variance meta-analysis. Test of the hypothesis that the effect was zero, was one sided, with alternate hypothesis that the effect had the same sign as in discovery. Effect sizes in discovery and replication were then compared by calculating the ratio (alpha) of replication effect sizes to discovery effect sizes:

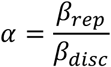

and its standard error using the following formula, reflecting the Taylor series expansion of the denominator for SE:

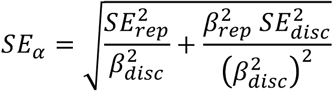

where *rep* and *disc* are replication and discovery, respectively. Alpha was then inverse-variance meta-analysed across all SNPs to test for collective evidence that the discovery SNPs influence longevity.

### Age and sex-stratified effects

Calculation of age and sex stratified effect sizes was done using the full Cox model (Equation 1), imputed dosages and the package “Survival” in R. We split the full analysis into age decades from 40 to 90 and a wider band, 90-120, beyond that, excluding any parent who died at an age younger than the age band and treating any parent who died beyond the age band as alive at the end of the age band. We thus had, across independent periods of life, estimates of the hazard ratio by decade of age and parent sex, along with standard errors. This gave estimates of the hazard ratio beta(age band, sex) in each age band and sex.

We tested the effect of age and sex, by fitting the linear model beta(age band, sex) = intercept + beta1 x ageband + beta2 × sex + e, where e is independent, but with known variance (the square of the SE in the age/sex stratified model fit) and using the rma function from the R package “metafor” which uses **known** variances of dependent variables. The process is more easily understood by examining the age and sex related effect size graphs, and recognising we are fitting an age and sex as explanatory variables, considering the standard error of each point shown.

### Causal gene prediction

In order to more accurately implicate causal genes and methylation sites from the detected loci associated with human lifespan, Summary-level Mendelian Randomisation (SMR) and HEterogeneity In InDependent Instruments (HEIDI) tests(49) were performed on our combined discovery and replication CES statistics. Three separate analyses were performed. First, cis-eQTL scan results from peripheral blood tissue from two previous studies, the Westra data(50) and CAGE data (51), were used to prioritize causal genes. Second, cis-eQTL signals (SNPs with FDR < 0.05) for 48 tissues from the GTEx consortium (52) were used to prioritize causal genes in multiple tissues. Third, genome-wide methylation QTL (mQTL) scan signals in blood tissue from the Brisbane Systems Genetics Study and Lothian Birth Cohort (53) were used to predict causal CpG sites associated with human lifespan loci. All results from SMR test passing a 5% FDR threshold where the HEIDI test P > 0.05 were reported.

### Fine-mapping using LASSO regression

Selection Operator for JOint multi-SNP analysis (SOJO) (54) was used to perform conditional fine-mapping analysis of the lifespan loci. The SOJO procedure implements LASSO regression for each locus, which outperforms standard stepwise selection procedure (e.g. GCTA-COJO), based on summary association statistics and the European-ancestry 1000 Genomes samples for LD reference. We based the SOJO analysis on our CES summary association statistics from the discovery population and used the replication cohort as validation sample to optimise the LASSO tuning parameters for each locus. Loci were defined prior to analysis as 1Mb windows centred at each top variant from the GWAS. For each locus, based on discovery data, we recorded the first 30 variants entering the model and the tuning parameters for these entering points along the LASSO path, as well as the LASSO results at the tuning parameters. For each recorded tuning parameter, we then built a polygenic score and computed the proportion of predictable variance from the regional polygenic score in the validation sample. The best out-of-sample R squared is reported, together with the selected variants per locus.

### Identification of disease traits underpinning variation in lifespan

For the lead lifespan SNPs, we used UK Biobank and PhenoScanner(25) to see if we could identify disease pathways underpinning the longevity effects and also provide supporting evidence for the plausibility of the lifespan effects. Because the UK Biobank analysis is reusing the same samples, there is a risk of chance associations with lifespan being caused by chance associations with disease (due to correlation between the two phenotypes), but for true associations, the data is more comparable across SNPs. We tested lead SNPs and candidates from Table 1 but loci that were only significant under iGWAS were precluded from this analysis due the to the potential circularity arising from the prior focus on disease-inducing SNPs in building of the prior for iGWAS.

### UK Biobank Disease-Wide Association Study

SNPs reaching genome-wide significance in the discovery cohort GWAS, combined discovery and replication GWAS, as well as candidate SNPs, were tested for association with self-reported diseases in the discovery sample of 325,292 unrelated, genomically British subjects, their siblings, and each parent separately. Diseases of subject relatives were already coded into broad disease categories by UK Biobank. For offspring, ICD codes had been recorded which we grouped into similar categories (hypertension, cerebral infarction, heart disease, diabetes, dementia, depression, stress, pulmonary disease, and cancer, in accordance with Table S16, although cancer in subjects was more directly taken as the trait of reporting number of cancers >0). The trait of reporting these diseases (separately for each relative and the subject themselves) was then tested for association with genotypic dosage for our candidate SNPs. The model fitted was a logistic regression of **not** reporting the disease, using the same covariates as the main analysis with the addition of subject age, and estimated the log odds ratio of protection from disease for each copy of the lifespan protective allele. Statistically significant results were corrected for multiple testing using Benjamini-Hochberg with an FDR threshold of 5%. Diseases were then grouped into broader categories, and number of protective (+) or deleterious (–), corresponding to the sign of the log odds ratio, associations were counted for each variant. Each lead SNP could associate many times with a disease category, as repeated associations across relatives, or diseases within the category (or both) might be found (Table S17).

Lead SNPs from the discovery GWAS, combined discovery and replication GWAS, as well as candidate SNPs and close proxies (r^2^ > 0.8), were scanned for known associations using PhenoScanner(25). All associations passing a 5% FDR threshold were divided using keywords into broad disease categories or “other”, which were then further curated. These categories were CVD – Cardiovascular diseases and risk factors, such as myocardial infarction, aortic valve calcification, hypertension, and cholesterol and triglyceride levels; IMMUNE – autoimmune and chronic inflammation disorders, such as type 1 diabetes, rheumatoid arthritis, systemic lupus erythematosus, inflammatory bowel diseases, and autoimmune liver and thyroid disease; PULMONARY – pulmonary function and disease (exc. cancer): asthma, chronic pulmonary obstructive disorder, respiratory function and airflow obstruction; DIABETES – type 2 diabetes and risk factors including glucose, HbA1c, and insulin levels; OBESITY – Anthropometric measures such as BMI, body fat percentage, waist/hip circumference, weight, and obesity; NEURO – Neurological disorders, such as Alzheimer’s, Parkinson’s, and Huntington’s disease, as well as depression, smoking addiction, and neuroticism. CANCER – Any association with cancer. Identical traits (such as “LDL”, “cholesterol LDL”, and “Serum LDL C”) and similar traits (such as “Childhood BMI”, “BMI in females”, and “BMI in males”) were grouped, keeping only the strongest association. In total, there were 131 unique traits of which 61 could be classified into one of the categories (15 CVD, 7 OBESITY, 14 NEURO, 4 DIABETES, 15 IMMUNE, 4 PULMONARY, 2 CANCER) (Table S18).

### Lifespan variance explained

As our tests of association with disease of lifespan SNPs in UK Biobank depended on power, reflecting the UK Biobank sample structure and (parental) disease prevalence, or the study designs underpinning PhenoScanner, we sought an independent, large set of disease-associated SNPs with strong effects on lifespan. A large number of SNPs per disease category, especially other cancers, were used to ensure that diseases were not under-represented when testing for association with lifespan. The latest, genome-wide significant disease SNPs from European ancestry studies were retrieved from the GWAS catalog (14 March 2018), based on string matching within reported trait names. For Alzheimer’s / Parkinson’s disease, these were “alzh” and “parkin”; for CVD, these were “myocard”, “cvd”, “cardiovascular”, “coronary”, and “artery disease"; for Type 2 diabetes this was “type 2 diabetes”; for cancers, this was “cancer”, “noma”, “ioma”, “tumo[u]r”, and “leukemia”. Cancers were then divided in Lung cancer and Other cancers based on the presence or absence of the keyword “lung”. The Smoking / Lung cancer category was created by adding traits containing the keywords “smoking” and “chronic obstructive” to the lung cancers. Each category was manually checked to include only associations with the diseases themselves or biomarkers of the diseases. Although some throat cancers are often caused by smoking and alcohol consumption, we did not treat these as smoking loci; in practice, this choice had no effect as the only significant throat cancer locus (oesophageal cancer near CFTR) was discounted as secondary pleiotropic – see below.

SNPs missing from the CES meta-analysis summary statistics were imputed from the closest proxy (min. r^2^ > 0.9) or averaged from multiple proxies if equally close. SNPs without effect sizes, SNPs matching neither our reference nor effect alleles, and SNPs with reported frequencies differing by more than 0.3 from our own were excluded. The remaining SNPs were subdivided into independent (r^2^ < 0.1) loci 500kb apart, keeping the SNP most strongly associated with lifespan in the CES meta-analysis – thus proportional to the lifespan GWAS test statistic rather than disease structure in UK Biobank. Lastly, where possible, loci were tested for association with their disease category in UK Biobank parents and siblings (using the same models as our disease-wide association study). Effects reported in the GWAS catalog for which we found the pooled estimate from our association study was in the opposite direction were flipped (if P < 0.05) or discarded (if P ≥ 0.05).

Our final dataset consisted of 555 disease SNPs (81 neurological, 72 cardiovascular, 65 diabetes, 22 smoking/lung cancer, and 315 other cancers). Lifespan variance explained (LVE) was calculated as 2pqa^2^, where p and q are the frequencies of the effect and reference alleles in our lifespan GWAS, and a is the SNP effect size in years of life(55). To assess pleiotropy, SNPs were tested against other disease categories, and where possible, the relative strengths of standardised associations between disease categories were compared. SNPs associating more strongly with another disease, as defined by a Z statistic more than double that of the original disease, were marked as pleiotropic and secondary. Whilst strength of association would not normally be perceived as appropriately measured in this way (odds ratio being more conventional and independent of prevalence), here we are interested in the excess number of disease cases in the population due to the variant, so any locus with a moderate OR for a highly prevalent disease is judged more causative of that disease than a locus with a (somewhat) higher OR for a very rare disease, as the number of attributable cases will be lower. The Z statistic captures this – given that p and q are obviously the same (same SNP, same population). Correspondingly, for diseases only present in one sex, the other sex was treated as all controls. Whilst this halves the apparent effect size, the required measure is the amount of disease caused across the whole population. A SNP conferring similar attributable counts of CVD and breast cancer in women, but also CVD in men, is causing CVD more than cancer across the population. Correspondingly selection pressure on the breast cancer effect is half that for a matching effect in both sexes. SNPs conferring both an increase in disease and an increase in lifespan were marked as antagonistically pleiotropic. Unsurprisingly, in practice, there were one or more other diseases reduced by the SNP and therefore the reported disease-increasing association was considered secondary. Total LVE per disease category was calculated by summing SNPs not marked as secondary and with significant effects on lifespan, where significance was determined by setting an FDR threshold that allowed for 1 false positive among all SNPs tested (q ≤ 0.016, 60 SNPs). To compare the cumulative LVE of the top LVE loci, all non-secondary association SNPs from the disease categories were pooled and again subdivided into independent loci (r^2^ ≤ 0.1) 500kb apart. Applying an FDR threshold with the same criteria (q ≤ 0.022), a total of 45 (1 neurological, 23 cardiovascular, 4 diabetes, 6 smoking/lung cancer, and 11 other cancer) independent loci remained and their LVE was summed by disease category.

### Tissue and pathway enrichment

LD-score regression (57) indicated that between the CES and PDES discovery and discovery-replication meta results, the CES discovery sample had the highest SNP heritability, plausibly due to its uniformity of population sample. Stratified LD-score regression(56) partitions SNP heritability into regions linked to specific tissues and cell types, such as super-enhancers and histone marks, and then assesses whether the SNPs in these regions contribute disproportionately to the total SNP heritability. The CES statistics were selected and analysed using the procedure described by Finucane et al. (56), which included limiting the regressions to HapMap3 SNPs with MAF > 0.05 to reduce statistical noise. Results from all cell types were merged and then adjusted for multiple testing using Benjamini–Hochberg (FDR 5%).

The CES discovery-replication meta-analysis dataset was subjected to gene-based tests, which used up to 10^6^ SNP permutations per gene to assign P values to 26,056 genes, as implemented by VEGAS2 v2.01.17(58) Only directly genotyped SNPs from the UK Biobank array were used to facilitate practical runtimes. Using the default settings, all SNPs located within genes (relative to the 5’ and 3’ UTR) were included. Scored genes were then tested for enrichment in 9,741 pathways from the NCBI BioSystems Database with up to 10^8^ gene permutations per pathway using VEGAS2Pathway(59). Pathway enrichment P values were automatically adjusted for pathway size (empirical P) and further adjusted for multiple testing using Benjamini-Hochberg (FDR 5%).

DEPICT was also used to create a list of genes; however, this method uses independent SNPs passing a P value threshold to define lifespan loci and then attempts to map 18,922 genes to them. Gene prioritization and subsequent gene set enrichment is done for 14,461 probabilistically-defined reconstituted gene sets, which are tested for enrichment under the self-contained null hypothesis (60). Two separate analyses were performed on the combined CES discovery-replication summary statistics, using independent SNPs (>500kb between top SNPs) which were present in the DEPICT database. The first analysis used a genome-wide significance threshold (GW DEPICT analysis) and mapped genes to 10 loci, automatically excluding the major histocompatibility complex (MHC) region. The second used a suggestive significance threshold (P < 10^-5^), which yielded 93 loci and mapped genes to 91 of these, again excluding the MHC region. To test if pathways were significantly enriched at a 5% FDR threshold, we used the values calculated by DEPICT, already adjusted for the non-independence of the gene sets tested.

PASCAL was used with the same summary statistics and gene sets as DEPICT, except the gene probabilities within the sets were dichotomized (Z>3) (62), leading to the analysis of the same 14,461 pathways. PASCAL transformed SNP P values into gene-based P values (with default method “––genescoring=sum”) for 21,516 genes (61). When testing the pathways for overrepresentation of high gene scores, the P values are estimated under the competitive null hypothesis (69). These pathway empirical P values were further adjusted for multiple testing using Benjamini-Hochberg procedure.

### Age-related eQTLs enrichment

We identified SNPs in our GWAS (discovery plus replication combined CES) that were eQTLs i. e. associated with the expression of at least one gene with P <10^−4^ in a dataset provided to us by the eQTLGen Consortium (n=14,155 individuals). A total of 2,967 eQTLs after distance pruning (500kb) were present, of which 500 were associated with genes differentially expressed with age (34). We used Fisher’s exact test to determine, amongst the set of eQTLs, if SNPs which were associated with lifespan (at varying thresholds of statistical significance) were enriched for SNPs associated with genes whose expression is age-related.

### Polygenic lifespan score associations

We used the combined discovery and replication CES GWAS, excluding (one at a time) all Scottish populations (whether from Scottish UK Biobank assessment centres or Scottish LifeGen cohorts), Estonian populations and a random 10% of UK Biobank English and Welsh subjects to create polygenic risk scores using PRSice (63), where the test subjects had not been part of the training data. As we find polygenic risk scores developed using all (P ≤ 1) independent (r^2^ ≤ 0.1) SNPs (PRSP1), rather than those passing a tighter significance threshold are most predictive (highest standardised effect size; see Table S25 for comparison between thresholds), these were used in the prediction analysis.

To make cross-validated lifespan predictions using polygenic scores, our unrelated, genomically British sample was partitioned into training and test sets. The first test set consisted of Scottish individuals from UK Biobank, as defined by assessment centre or northings and eastings falling within Scotland (N = 24,059). The second set consisted of a random subset of the remaining English and Welsh population, reproducibly sampled based on the last digit of their UK Biobank identification number (#7, N = 29,815). The training set was constructed by excluding these two populations, as well as excluding individuals from Generation Scotland, from our GWAS and recalculating estimates of beta on that subset.

A third independent validation set was constructed by excluding the EGCUT cohort from the LifeGen sample and using the remaining data to predict lifespan in the newly genotyped EGCUT cohort(70), using unrelated individuals only (N = 36,499).

Polygenic survival scores were constructed using PRSice 2.0.14.beta(63) in a two-step process. First, lifespan SNPs were LD-clumped based on an r^2^ threshold of 0.1 and a window size of 250kb. To facilitate practical run times of PRSice clumping, only directly genotyped SNPs were used in the Scottish and English/Welsh subsets. The Estonian sample was genotyped on four different arrays with limited overlap, so here imputed data (with imputation measure R2>0.9) was used and clumped with PLINK directly (r^2^ = 0.1; window = 1000kb). The clumped SNPs (85,539 in UK Biobank, 68,234 in Estonia) were then further pruned based on several different P value thresholds, to find the most informative subset. For all individuals, a polygenic score was calculated as the sum of SNP dosages (of SNPs passing the P value threshold) multiplied by their estimated allele effect. These scores were then standardised to allow for associations to be expressed in standard deviations in polygenic scores.

Polygenic scores of test cohorts were regressed against lifespan and alive/dead status using a cox proportional hazards model, adjusted for sex, assessment centre, batch, array, and 10 principal components. Where parental lifespan was used, hazard ratios were doubled to gain an estimate of the polygenic score on own mortality. Scores were also regressed against diseases using a logistic regression adjusted for the same covariates plus subject age. As with previous disease associations, estimates were transformed so positive associations indicate a protective or life-extending effect, and effect estimates of first degree relatives were doubled. Meta-analysis of estimates between cohorts was done using inverse variance weighting. Where estimates between kin were meta-analysed, standard errors were adjusted for correlation between family members. This involved multiplying standard errors by 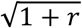 for each correlation (r) with the reference kin (Equation 2), which appears slightly conservative. As correlations between family member diseases were very low (range 0.0005 to 0.1048), in practice, this adjustment had no effect.

### Sensitivity of annuity prices to age

Market annuity rates for life annuities in January 2018 written to 55, 60, 65, and 70 year olds were obtained from the sharing pensions website http://www.sharingpensions.com/annuity_rates.htm (accessed 22 January 2018) and were £4158, £4680, £5476, £6075, £7105 respectively per year for a £100,000 purchase price. The arithmetic average increase from one quinquennial age to the next is 14 percent.

### URLs

MultiABEL: https://github.com/xiashen/MultiABEL/1

LDSC: https://github.com/bulik/ldsc

SMR/HEIDI: https://cnsgenomics.com/software/smr/

SOJO: https://github.com/zhenin/sojo/

DEPICT: https://www.broadinstitute.org/mpg/depict/

PASCAL: https://www2.unil.ch/cbg/index.php?title=Pascal GTEx: https://gtexportal.org/home/datasets

## Acknowledgements

We thank the UK Biobank Resource, approved under application 8304; we acknowledge funding from the UK Medical Research Council Human Genetics Unit, Wellcome Trust PhD Training Fellowship for Clinicians, the Edinburgh Clinical Academic Track (ECAT) programme (204979/Z/16/Z), the Medical Research Council Doctoral Training Programme (MR/N013166/1) and the AXA research fund. We thank Tom Haller of the University of Tartu, for tailoring RegScan so we could use it with compressed files(65). We would also like to thank the researchers, funders and participants of the LifeGen consortium(5).

## Contributions

PRJHT, NM, KL, KF, ZN, XF, AB, DC performed analyses.

TE, eQTLGen contributed data

XS, TE, KF, ZK, PKJ designed the experiments

PRJHT, NM, KL, KF, AB, XS, TE, ZK, JFW, PKJ wrote the manuscript

## Additional Information

eQTLGen Consortium

Agbessi M^1^, Ahsan H^2^, Alves I^1^, Andiappan A^3^, Awadalla P^1^, Battle A^4^, Beutner F^5^, Bonder MJ^6^, Boomsma D^7^, Christiansen M^8^, Claringbould A^6^, Deelen P^6^, van Dongen J^7^, Esko T^9^, Favé M^1^, Franke L^6^, Frayling T^10^, Gharib SA^11^, Gibson G^12^, Hemani G^13^, Jansen R^7^, Kalnapenkis A^9^, Kasela S^9^, Kettunen J^14^, Kim Y^4^, Kirsten H^15^, Kovacs P^16^, Krohn K^17^, Kronberg-Guzman J^9^, Kukushkina V^9^, Kutalik Z^18^, Kähönen M^19^, Lee B^3^, Lehtimäki T^20^, Loeffler M^15^, Marigorta U^12^, Metspalu A^9^, van Meurs J^21^, Milani L^9^, Müller-Nurasyid M^22^, Nauck M^23^, Nivard M^7^, Penninx B^7^, Perola M^24^, Pervjakova N^9^, Pierce B^2^, Powell J^25^, Prokisch H^26^, Psaty BM^27^, Raitakari O^28^, Ring S^29^, Ripatti S^14^, Rotzschke O^3^, Ruëger S^18^, Saha A^4^, Scholz M^15^, Schramm K^22^, Seppälä I^20^, Stumvoll M^30^, Sullivan P^31^, Teumer A^32^, Thiery J^33^, Tong L^2^, Tönjes A^30^, Verlouw J^21^, Visscher PM^25^, Võsa U^6^, Völker U^34^, Yaghootkar H^10^, Yang J^25^, Zeng B^12^, Zhang F^25^

1 Ontario Institute for Cancer Research Toronto Canada

2 Department of Public Health Sciences University of Chicago Chicago United States of America

3 Singapore Immunology Network Agency for Science, Technology and Research Singapore Singapore

4 Department of Computer Science Johns Hopkins University Baltimore United States of America

5 Heart Center Leipzig Universität Leipzig Leipzig Germany

6 Department of Genetics University Medical Centre Groningen Groningen The Netherlands

7 Vrije Universiteit Amsterdam The Netherlands

8 Cardiovascular Health Research Unit University of Washington Seattle United States of America

9 Estonian Genome Center University of Tartu Tartu Estonia

10 Exeter Medical School University of Exeter Exeter United Kingdom

11 Department of Medicine University of Washington Seattle United States of America

12 School of Biological Sciences Georgia Tech Atlanta United States of America

13 MRC Integrative Epidemiology Unit University of Bristol Bristol United Kingdom

14 University of Helsinki Helsinki Finland

15 Institut für Medizinische InformatiK, Statistik und Epidemiologie, LIFE – Leipzig Research Center for Civilization Diseases Universität Leipzig Leipzig Germany

16 IFB Adiposity Diseases Universität Leipzig Leipzig Germany

17 Interdisciplinary Center for Clinical Research, Faculty of Medicine Universität Leipzig Leipzig Germany

18 Lausanne University Hospital Lausanne Switzerland

19 Department of Clinical Physiology Tampere University Hospital and Faculty of Medicine and Life Sciences, University of Tampere Tampere Finland

20 Department of Clinical Chemistry Fimlab Laboratories and Faculty of Medicine and Life Sciences, University of Tampere Tampere Finland

21 Department of Internal Medicine Erasmus Medical Centre Rotterdam The Netherlands

22 Institute of Genetic Epidemiology, Helmholtz Zentrum München, German Research Center for Environmental Health, Neuherberg, Germany

23 Institute of Clinical Chemistry and Laboratory Medicine University Medicine Greifswald Greifswald Germany

24 National Institute for Health and Welfare University of Helsinki Helsinki Finland

25 Institute for Molecular Bioscience University of Queensland Brisbane Australia

26 Institute of Human Genetics Helmholtz Zentrum München München Germany

27 Cardiovascular Health Research Unit, Departments of Epidemiology, Medicine, and Health Services University of Washington Seattle United States of America

28 Department of Clinical Physiology and Nuclear Medicine Turku University Hospital and University of Turku Turku Finland

29 School of Social and Community Medicine University of Bristol Bristol United Kingdom

30 Department of Medicine Universität Leipzig Leipzig Germany

31 Department of Medical Epidemiology and Biostatistics Karolinska Institutet Stockholm Sweden

32 Institute for Community Medicine University Medicine Greifswald Greifswald Germany

33 Institute for Laboratory Medicine, LIFE – Leipzig Research Center for Civilization Diseases Universität Leipzig Leipzig Germany

34 Interfaculty Institute for Genetics and Functional Genomics University Medicine Greifswald *SI Appendix*

## SI Appendix - section 1: Implication of causal genes and methylation sites

SMR-HEIDI implicates specific causal genes within identified gene regions by finding similar patterns in SNP effects on gene expression and the trait in question(49). Cis-eQTL scan results from two previous studies on peripheral blood tissue, Westra (50) and CAGE (51), were used to prioritize causal genes within loci reaching genome-wide significance in our discovery GWAS, combined cohort GWAS, or iGWAS. We also included *d3-GHR*, *5q33.3/EBF1*, and *FOXO3* in the analysis.

At a 5% FDR threshold for SMR and a P > 0.05 threshold for HEIDI test, 11 genes (*PSRC1*, *ARPC1B*, *SH2B3*, *PSMA4*, *FES*, *FURIN*, *OCIAD1*, *BECN1*, *ATP6V0A1*, *KANK2*, and *SESN1*) are implicated in 9 separate gene regions (Table S5).

In order to expand the SMR-HEIDI analysis to other human tissues, we extracted cis-eQTL (SNPs with FDR < 5%) signals for 48 tissues from the GTEx consortium(52). At a 5% FDR threshold for SMR and a P > 0.05 threshold for HEIDI test, 27 unique genes from 11 loci across 25 tissues are implicated as causal (Table S12). Of these, the six most statistically robust associations (FDR < 1%) are (tissue:gene) Muscle Skeletal:CELSR2, Liver:PSRC1, Cells Transformed fibroblasts:FES, Liver:CELSR2, Esophagus Mucosa:PSRC1, Cells Transformed fibroblasts:BECN1.

We also extended the SMR-HEIDI test to genome-wide methylation QTL (mQTL) scan signals (blood tissue)(53) from BSGS and LBC studies, so that causal CpG sites associated with the human lifespan GWAS signals can be predicted. All results from SMR test with a 5% FDR threshold where the HEIDI test P > 0.05 were reported (Table S13). 57 sites at 16 loci were implicated as to having causal effects on the lifespan including some within loci with established biological relevance to lifespan (*CHRNA3/5*, *APOE*, and *LPA*) which the eQTL dataset may not have had sufficient power to reveal. The 9 most statistically significant results (FDR < 0.1%) were for (gene region: CpG probe) CHRNA3/5:cg04882995, CHRNA3/5:cg04140906, CLU:cg26027576, FURIN/FES:cg05469396, LAMA5:cg24112000, ARPC1:cg04083712, BECN1:cg04987362, *HLA-DQA1*:cg18060330, and *HLA-DQA* 1:cg06871764.

## SI Appendix - section 2: Disease associations of lead lifespan SNPs

We see three variants (at or near *ATXN2/BRAP*, *CDKN2B-AS1* and *LPA*) with multiple protective associations against CVD associating with a risk increase in self-reported cancer. Additionally, 8 loci (at or near 13q21.33, *CLU*, *HLA-DQA1*, *HIST1*, *LAMA5*, *PSORS1C3*, *MICA/MICB*, *EGLN2/CYP2A6*) show a protective effect on cancer and no statistically significant association with CVD. Of these, all but two associations are protective from lung cancer, the exceptions being *HLA-DQA1*, which associates with reduced respiratory disease and cancer in general, and *LAMA5*, which associates with reduced bowel cancer. For diabetes, we see lifespan-increasing variants with a protective effect at or near *CDKN2B-AS1*, *ATXN2/BRAP*, *MAGI3*, *TMEM18*, *CHW43*, *HP*, *BECN1*, *PSORS1C3*, and *EXOC3L2/MARK4*, but with deleterious associations on diabetes at or near *CELSR2/PSRC1*, *LDLR* and *APOE*. For neurological diseases, we see 4 protective associations, all from the life-lengthening variant near *APOE*, and 5 deleterious associations from SNPs at or near *CLU*, *FURIN/FES*, *KCNK3* and *EXOC3L2/MARK4*. Finally, for pulmonary diseases, we see lifespan increasing variants with a protective effect in 9 analyses (in loci at or near *HLA-DQA1*, *CHRNA3/5*, *CHRNA4*, *TMEM18*, *MICA/MICB*, *FOXO3*, and *SEMA6D*), but with deleterious effects for the life lengthening allele at or near *LPA* and *APOE*. Conversely, when considering the disease associations SNP-by-SNP, we found that 16 out of our 25 genome-wide significant longevity variants had more than one protective association with the diseases considered. Of the 11 SNPs for which we found replication, 9 SNPs have more than one protective association with disease, *ZW10* and 13q21.31 being the exceptions. Furthermore, lead SNPs at or near *CELSR2/PSRC1*, *LPA*, *CDKN2B-AS1*, *ATXN2/BRAP*, *CHRNA3/5*, *FURIN/FES*, *APOE*, *KCNK3* and *IGF2R* showed 5 or more protective associations. Three variants (at or near *LPA*, *CLU*, and *APOE*) with life-lengthening associations also showed more than one deleterious association with disease.

Lookup of genome-wide significant SNPs and candidate SNPs on PhenoScanner (25), together with their closest proxies (r2 > 0.8), show very similar patterns of association, but now in independent data (again loci that were only significant under iGWAS were precluded). We grouped phenotypes by broad disease category or obesity as summarised in Table S7 (for definition of the disease categories see Methods and see Table S18 for full detail). In particular, of the 210 unique PhenoScanner disease associations (FDR < 5%) found for the 45 SNPs, 80 associations were with CVD (especially at or near CELSR2/PSRC1, ATXN2/BRAP, LDLR, LPA, CDKN2B-AS1, and HP), and 48 were with obesity traits (especially at or near TMEM18, ATXN2/BRAP, FOXO3, and KCNK3). Of the remaining 82 associations, 33 were for neurological disorders and addiction (primarily at or near FURIN/FES and CHRNA3/5), 33 were for immune disorders (almost half near ATXN2/BRAP, and a third near MAGI3 and MICA/MICB), 8 were for type 2 diabetes risk factors, 6 were for pulmonary disorders (mostly near CHRNA3/5), and two were for cancers (one lung cancer, one glioma). Conversely when considering PhenoScannner associations SNP-by-SNP, we found 8 or more statistically significant disease associations for each of the lead SNPs at or near ATXN2/BRAP, CELSR2/PSRC1, CHRNA3/5, FOXO3, HIST1, MICA/MICB, TMEM18, FURIN/FES, and HP, mostly concordant with our own PheWAS and supporting their putative roles in lifespan. At the same time, we found 2 or fewer statistically significant disease associations for lead SNPs at or near 13q21.31, 13q21.33, *ANKRD20A9P*, *BECN1*, *CHRNA4*, *CHW43*, *FPGT/TNNI3K*, *LAMA5*, *SEMA6D*, *USP2-AS1*, *USP42*, *BEND3*, *d3-GHR*, *MC2R*, *TMTC2*, *ZW10*, *ARPC1*, *EXOC3L2/MARK4*, *IL6*, *TOX*.

These disease associations are broadly consistent with mortality risk factor associations from our iGWAS. When performing a lookup of 1% FDR iGWAS SNPs in the mortality risk factor studies underpinning the iGWAS, we find the lead genome-wide significant loci either show strong clustering of blood lipids and cardiovascular disease, moderate clustering of metabolic and neurological traits, or weak but highly pleiotropic clustering amongst many of the traits considered (Fig. S9).

We next looked up the 82 FDR < 1% iGWAS SNPs within the 16 risk factor GWAMAs used to form the iGWAS prior. As the iGWAS had enriched SNPs with disease associations, this was not an unbiased sample; nonetheless, the pattern of association is still of interest. Using a Bonferroni corrected (16 traits, 82 SNPs) threshold of 3.81 × 10^-5^, we find 52 associations between our identified lifespan SNPs and risk factors. Education Level (years of schooling - 10), LDL Cholesterol (9) BMI (8), Coronary Artery Disease (8) show the highest number of statistically significant associations. Conversely SNPs near *CELSR2/PSRC1* and *BUD13/APOA5* show evidence of pleiotropy, with associations with three or more risk factors (Table S10).

## SI Appendix - section 3: Extended Discussion

The functions, in the context of lifespan and longevity, of the newly replicated loci *CDKN2B-AS1*, *ATXN2/BRAP*, and 5q33.3/EBF1 have been described in detail previously (6, 15). Briefly, variants within these loci are known to play a role in cardiovascular disease, hypertension, and autoimmune disorders, matching disease associations in our own study. Pilling et al. (6) suggest a causal role for *CDKN2B-AS1* lncRNA expression on lifespan, and we identify a causal role for *SH2B3* gene expression in the *ATXN2/BRAP* locus, admittedly a highly pleiotropic region. The causal element for lifespan near 5q33.3/EBF1 remains unclear.

The *FURIN/FES* variant associates with extended lifespan, decreased *FURIN* and increased *FES* expression(71). The gene product, Furin, is a pro-protein convertase, linked to dysregulation of lipid levels(72) and progression of atherosclerosis(73), matching the protective role of the lifespan-increasing variant observed against heart disease and hypertension. Despite both proteins playing a role in cancer development – Furin promoting metastasis(74, 75) and Fes displaying both pro and anti-tumorigenic effects(76) – we did not observe any significant associations between the lead variant and cancer.

The lifespan-increasing variant rs61905747_A near *ZW10* is associated with decreased expression of *USP28* and increased expression of two uncharacterised pseudogenes(71). *USP28* is a deubiquitinase that stabilises the oncogene *MYC(77*) and is upregulated in multiple cancers(78-81). While we find no disease associations beyond protection from CVD for our lead variant (95% CI logOR –0.10 to –0.03), Law et al. find a proxy in moderate LD (rs61904987_C, r^2^ ~ 0.6) is associated with decreased chronic lymphocytic leukaemia (95% CI logOR –0.28 to –0.15)(82), suggesting decreased *USP28* expression may extend lifespan by conferring a protection from CVD and some cancers.

The variant near *PSORS1C3* is located within a known psoriasis susceptibility locus near *HLA-C* (83, 84), but the complex LD structure complicates identification of a causal gene. Psoriasis is linked to higher risk of myocardial infarction, and life expectancy of men and women with severe psoriasis is decreased by an average of 3.5 and 4.4 years, respectively (85). The same region has also been linked to lung cancer, follicular lymphoma, and multiple myeloma susceptibility (86-88), although we only find an association between the SNP and protection against lung cancer.

The variant within cytogenetic band 13q21.31 is located in a gene desert. However, somatic deletion of the 13q21-q22 region is frequently observed in *non-BRCA1/BRCA2* breast cancer, suggesting the region contains functional elements involved in tumour suppression(89). Altered breast cancer susceptibility would match the female-specific effects on lifespan we observed for both intergenic lifespan SNPs 13q21.31 and 13q21.33 within this region.

Despite the mixed evidence for association we found with lifespan, *CELSR2* is known to affect cardiovascular disease in diverse populations (17, 90-92) and the locus is known to have sex-specific effects on lipid metabolites (93, 94). It is likely *CELSR2* affects lifespan and in a sex-specific way, with our CES replication failing due to sex specificity and the PDES replication being (very) slightly underpowered at a 5% significance level (P = 0.0565).

Our findings validate the results of a previous Bayesian analysis (iGWAS) performed on a subset (n=116,279) of the present study’s discovery sample (35), which highlighted two loci which are now genome-wide significant in conventional GWAS in the present study’s larger sample. iGWAS thus appears to be an effective method able to identify lifespan-associated variants in smaller samples than standard GWAS, albeit relying on known biology.

Lipid metabolites – particularly cholesterol metabolites – have well-established effects on atherosclerosis, type-2-diabetes, Alzheimer’s disease, osteoporosis, and age-related cancers (95). It is therefore no surprise that lifespan genetics are enriched for lipid metabolism genes, considering the mortality risk associated with these diseases. Lipid levels also play a role in brain function, with cholesterol being necessary for dendrite differentiation and synaptogenesis (96) and altered lipid metabolism causing multiple neurodegenerative diseases (97). Pilling et al. (6) implicated nicotinic acetylcholine receptor pathways in human lifespan, which we detect at nominal significance (P = 2×10^-4^), but not quite at 5% FDR correction (q = 0.0556). Instead we highlight more general synapse and dendrite pathways, and identify the dorsolateral prefrontal cortex (DLPC) as an ageing-related tissue. Indeed, the DLPC, which is involved in smoking addiction (98) and dietary self-control (99), has been found to be especially vulnerable to age-related synaptic death (100).

Whilst it has previously been shown that transcriptomic age calculated based on ARGs is meaningful in the sense that its deviation from the chronological age is associated with biological features linked to aging (101), the role of ARGs in ageing was unclear. A gene might be an ARG because (i) it is a biological clock (higher expression tracking biological ageing, but not influencing ageing or disease); (ii) it is a response to the consequences of ageing (e.g. a protective response to CVD); (iii) it is an indicator of selection bias: if low expression is life-shortening, older people with low expression tend to be eliminated from the study, hence the average expression level of older age groups is higher. However, our results now show that the differential expression of many of these genes with age is not only a biomarker of aging, but the genes identified by Peters et al(101) are enriched for direct effects on lifespan.

The strengths of our study undoubtedly include its size with over one million lifespans being considered and the partition into independent discovery and replication, albeit mitigated by the power-reducing effects of using (two) parents rather than genotyped subjects. Meta-analysing under two contrasting assumptions of sexual dimorphism, also avoided the loss of power associated with making the wrong assumption for a locus. However, despite its size, definitive identification of ageing pathways beyond disease remains elusive in humans. In addition, our replication was not sufficiently powered to allow for a multiple testing adjustment across discovered alleles, the risk of some false positives is thus increased, albeit mitigated by the consistency with other lines of evidence, for example strong and previously well-known disease susceptibilities at many of the loci.

We also show how disease informed lifespan GWAS (iGWAS) scan can improve statistical power and identification of novel loci. Its main limitation is the inability to discover any new mechanisms that are not acting through disease predisposition. Interestingly, the observed lifespan-modulating effects of the discovered SNPs and the expected effects based on their known disease associations are still quite far apart, indicating that there are many heritable life-shortening conditions have not yet (sufficiently) studied by GWAS.

Our observation, that despite a larger dataset, we consider our study only moderately powered, whereas Pilling et al (6) stated “indeed the power (>99% to detect an allele of 1% minor allele frequency accounting for 0.1% of phenotype variance) is sufficient to suggest that we have identified all moderate to larger effect common genotyped or imputed variants in our studied population” can perhaps be reconciled by recognising the phenotype to which Pilling et al’s statement related: Martingale residuals of parent survival, not subject lifespan. Variance explained by an allele for the former will be several times less than for the latter (more pertinent) trait, due to the mitigating effects of parent imputation and the proportion still alive.

---

**Fig. S1:**
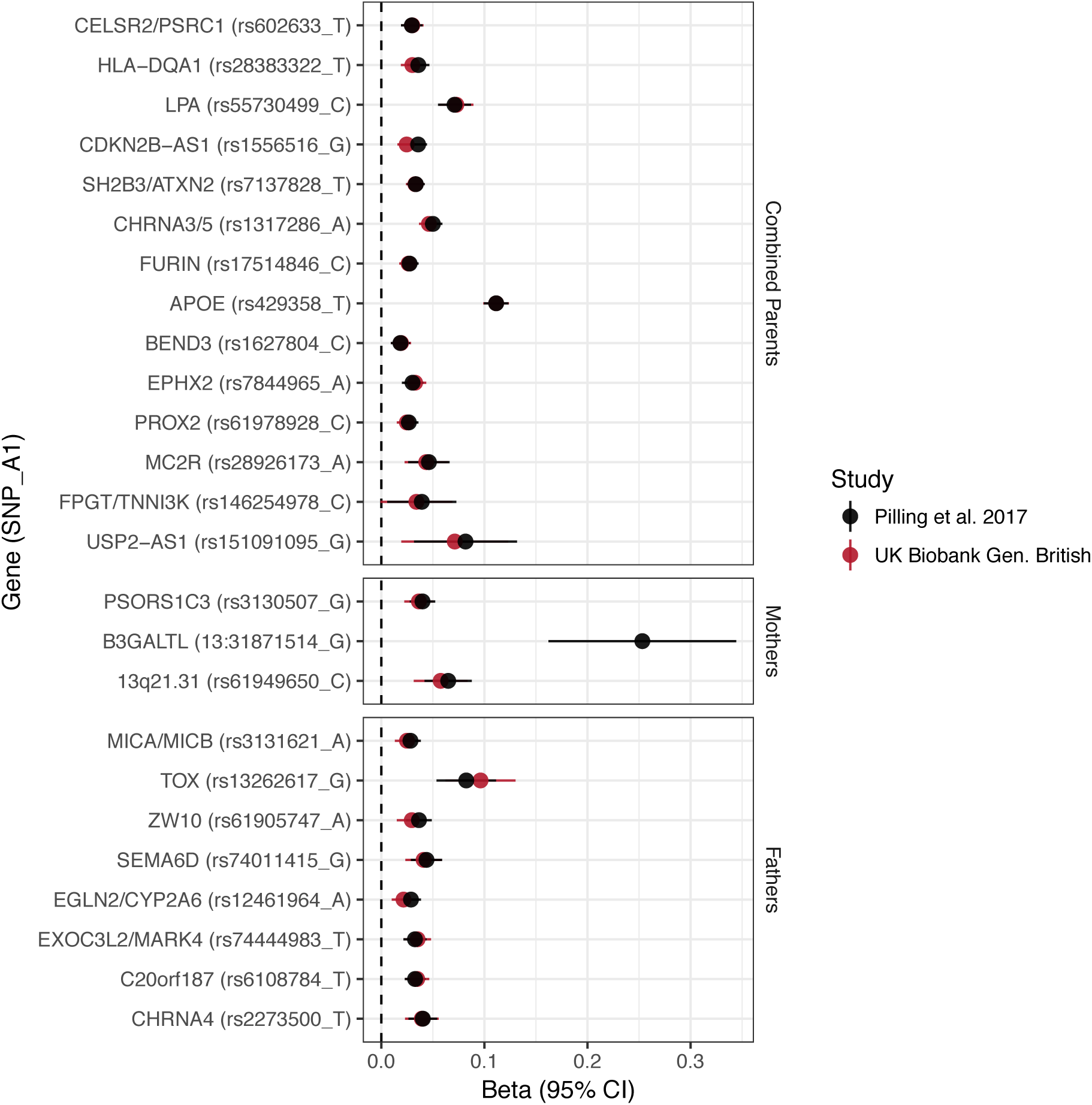
Concordance between inferred effect sizes from Pilling 2017 and our estimated effect sizes in a largely overlapping UK Biobank sample. Effect estimates from Pilling et al.(6) were converted to log_e_(protection ratio) based solely on the proportion dead in individual parental samples, or (for combined parents results) based on an empirical conversion factor from APOE (see Methods). By definition, the inferred effect estimate for APOE in combined parents is identical between the studies; all other estimates provide a measure of concordance between inferred and calculated effects for each locus. Gene names are as reported by discovery. Note, rs161091095 near USP2-AS1 is a proxy (r^2^ = 1.00) for rs139137459, the SNP reported by Pilling et al. No proxies could be found for 13:31871514_T_G. Gene – Nearby gene(s) as reported by discovery. SNP – rsID of SNP or proxy. A1 – Longevity allele. Beta – the estimated log_e_(protection ratio) for one copy of the effect allele. CI – Confidence Interval

**Fig. S2:**
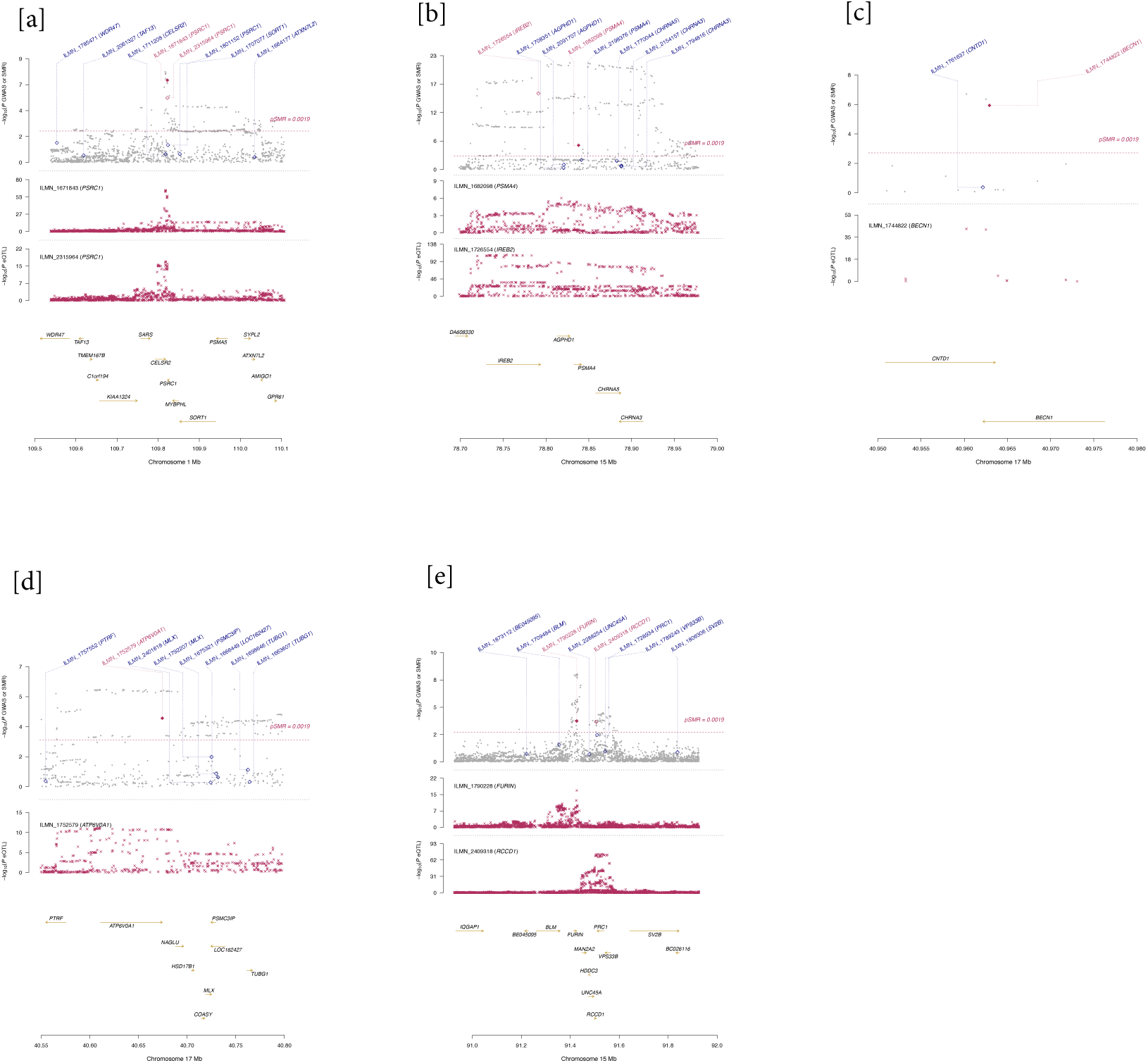
Loci with significantly predicted candidate genes using SMR-HEIDI test and the CAGE eQTL dataset (blood tissue) Lifespan GWAS and eQTL signals are plotted and compared. Gene expression probe names are provided with the corresponding gene names in brackets. The pSMR threshold corresponds to a significance level of FDR < 5%, and the gene expression probes that have SMR signal passing this threshold are displayed as red diamonds, otherwise blue. Filled diamonds indicate that the corresponding probes also pass the p > 0.05 threshold for the HEIDI test, i.e. the expressions of the particular genes possibly share causal variants with the lifespan GWAS signals. Panels are genomic loci around a) CELSR2/PSRC1; b) CHRNA3/5; c) BECN1; d) ARPC1; e) FURIN/FES

**Fig. S3:**
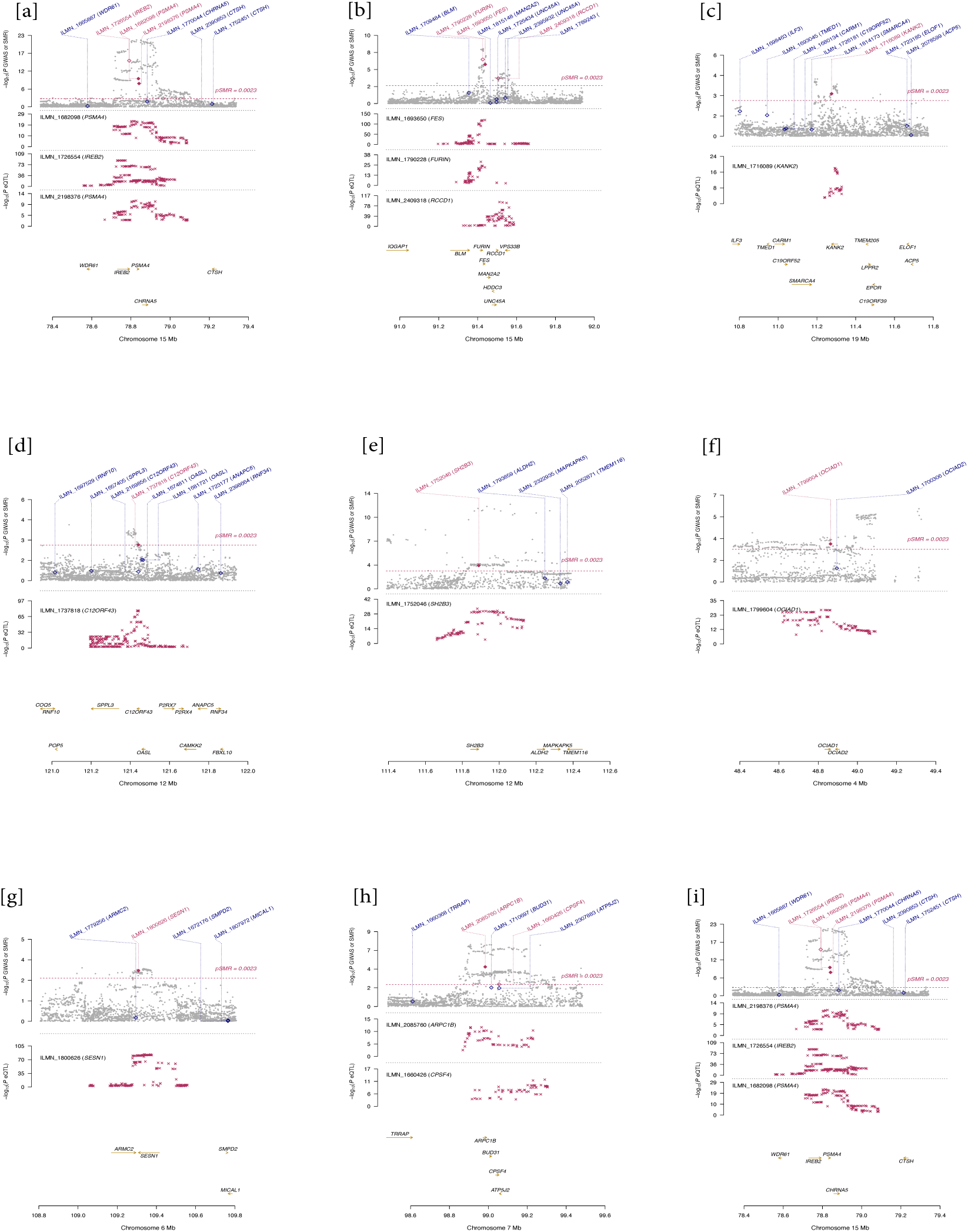
Loci with significantly predicted candidate genes using SMR-HEIDI test and the Westra eQTL dataset (blood tissue) Lifespan GWAS and eQTL signals are plotted and compared. Gene expression probe names are provided with the corresponding gene names in brackets. The pSMR threshold corresponds to a significance level of FDR < 5%, and the gene expression probes that have SMR signal passing this threshold are displayed as red diamonds, otherwise blue. Filled diamonds indicate that the corresponding probes also pass the p > 0.05 threshold for the HEIDI test, i.e. the expressions of the particular genes possibly share causal variants with the lifespan GWAS signals. Panels are genomic loci around a) CHRNA3/5; b) FURIN/FES; c) KANK2; d) C12Orf43; e) ATXN2/BRAP; g) FOXO3; h) ARPC1; i) CHRNA3/5

**Fig. S4:**
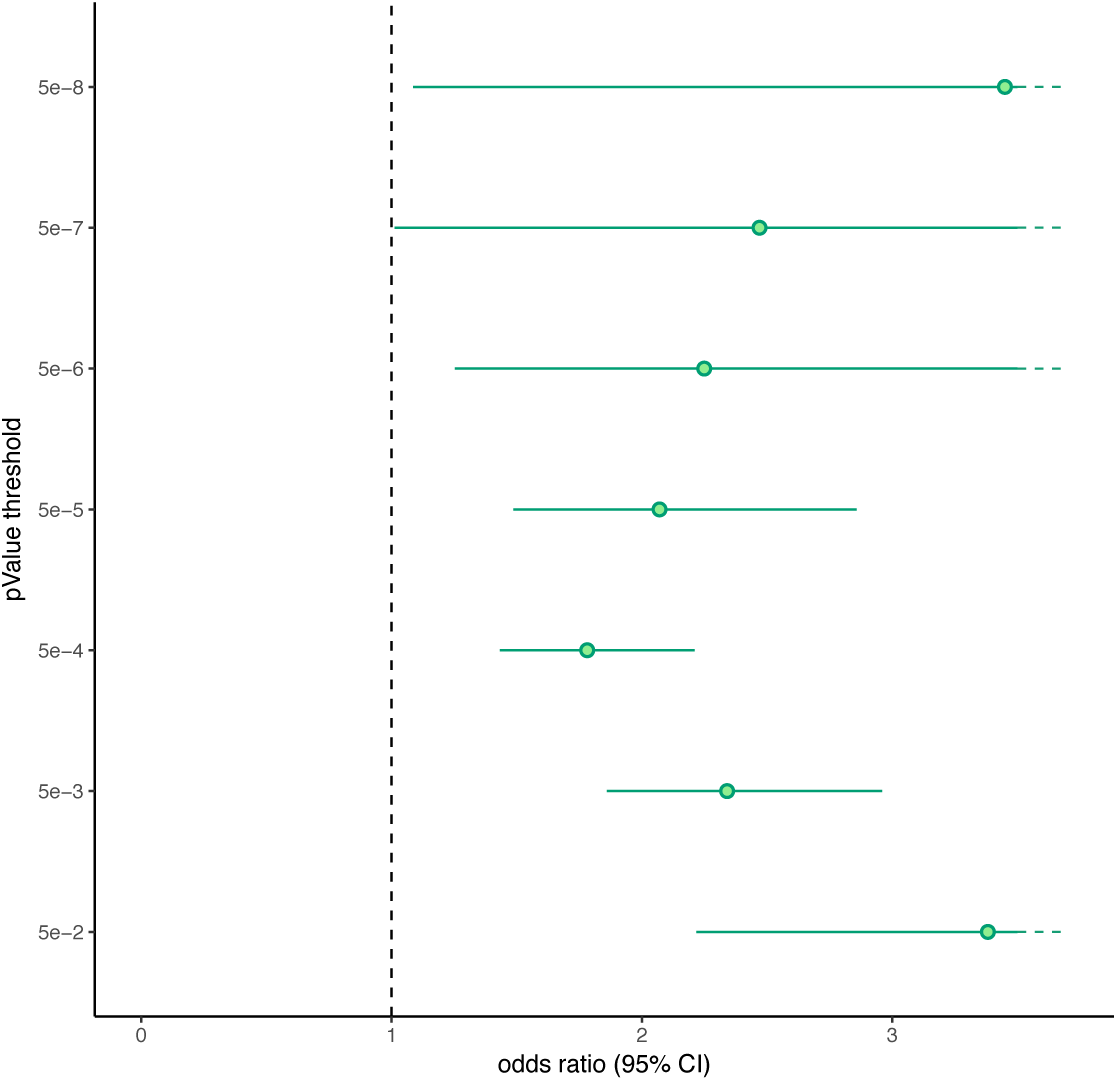
eQTL SNPs associated with lifespan for genes whose expression varies with age. We identified SNPs in our CES GWAS (discovery and replication combined) that were also eQTLs i.e. associated with the expression of at least one gene with P < 10^-5^ in a dataset provided to us by the eQTLGen Consortium. A total of 2,967 eQTLs after distance pruning (500 kb) were present, of which 500 were associated with genes differentially expressed with age(101). We used Fisher’s exact test to determine, amongst the set of eQTLs, if SNPs which were associated with lifespan (at varying thresholds of statistical significance) were enriched for SNPs associated with genes whose expression is agerelated. Odds ratio and 95% confidence intervals from Fisher’s exact test are represented for different thresholds of statistical significance. Upper bounds of confidence interval higher than 3.5 are represented by a dotted line (respectively 10.46, 5.68 and 5.37 for significance thresholds of 5e-8, 5e-7 and 5e-2). We see a significant enrichment (P < 0.05) in age-related eQTLs for all thresholds pointing out that age-related eQTLs, modulating the expression of genes differentially expressed with age, are enriched for lower than expected P value in our lifespan GWAS.

**Fig. S5:**
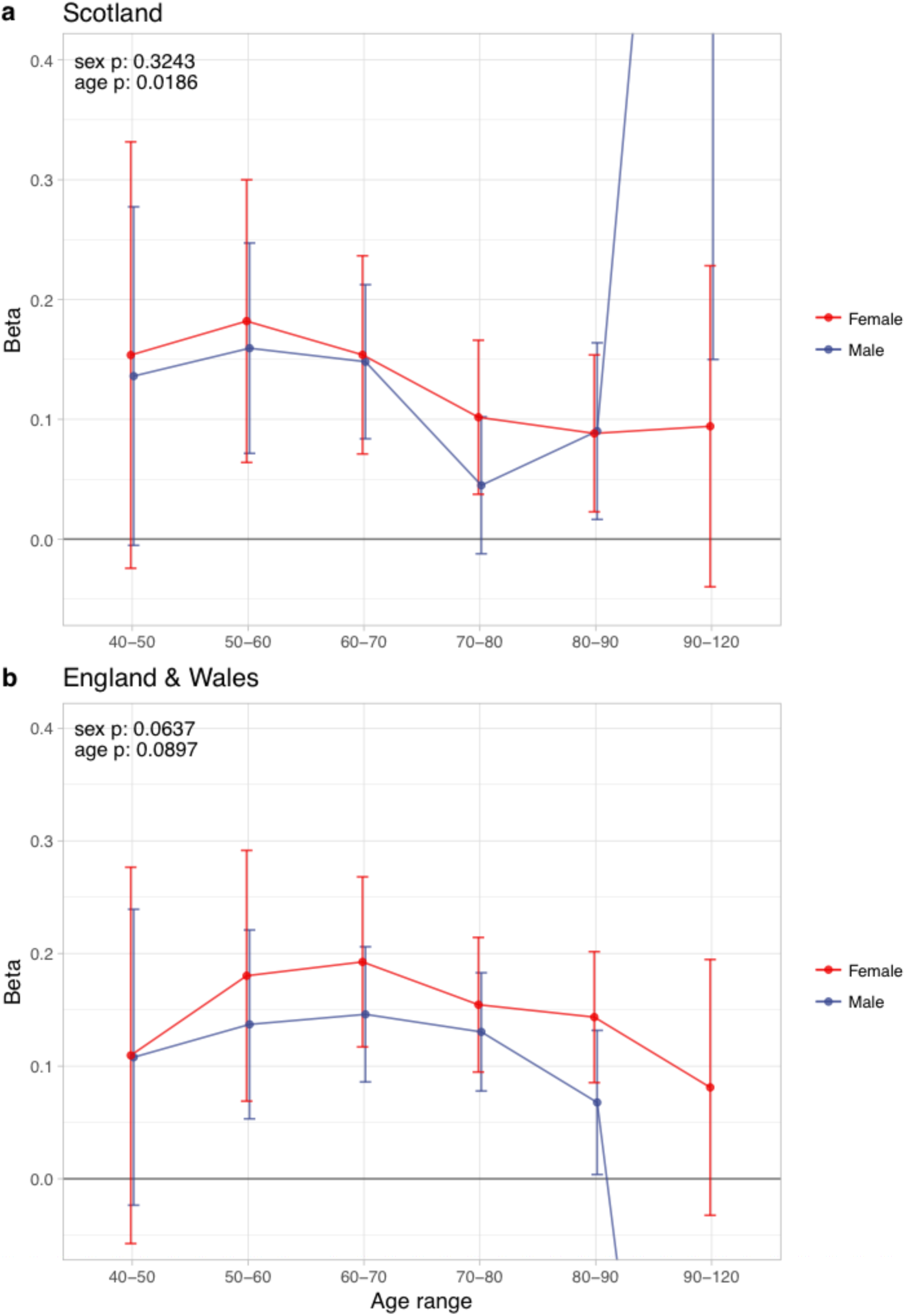
Sex and age specific effects of polygenic survival score (PRS) on parental lifespan of Scottish and English/Welsh subsamples of UK Biobank. a) Out of sample Scottish subset of UK Biobank; b) Out of sample English and Welsh subset of UK Biobank; Estimates for the PRS on father lifespan in the highest age range have very wide confidence intervals (CI) due to the limited number of fathers surviving past 90 years of age. The beta 95% CI for these estimates are 0.15 to 2.20 for Scottish subsamples and –1.34 to –0.16 for English & Welsh subsamples. Beta – loge(protection ratio) for 1 standard deviation of PRS for increased lifespan in self in the age band (i.e. 2 x observed due to 50% kinship), bounds shown are 95% CI; Age range – the range of ages over which beta was estimated; sex p – P value for association of effect size with sex; age p – P value for association of effect size with age

**Fig. S6:**
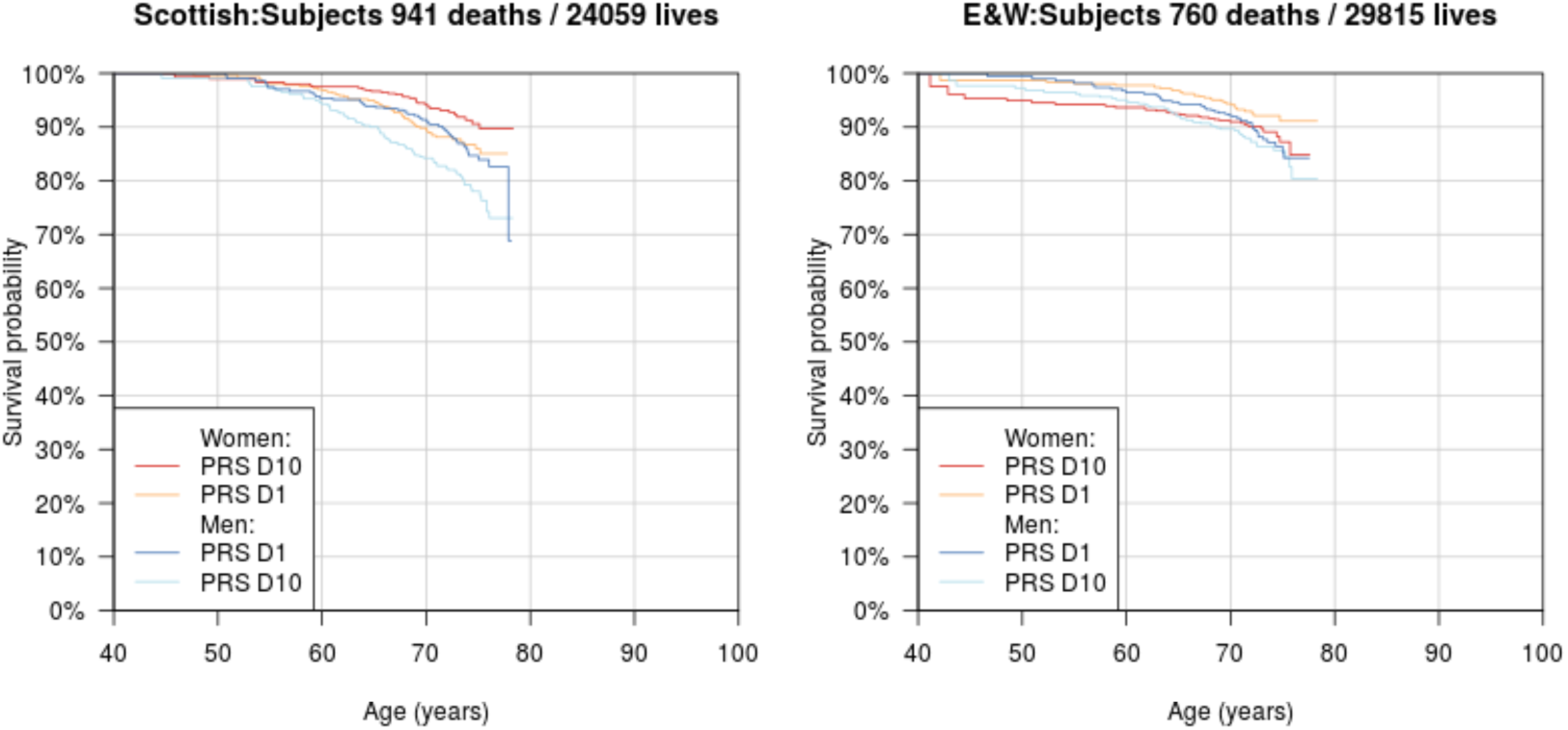
Survival Curves for highest and lowest deciles of lifespan polygenic risk score in UK Biobank subjects. A polygenic risk score was made for each subject using GWAS results that did not include the subject sets under consideration. Subject survival information (age entry, age exit, age of death (if applicable) was used to create Kaplan-Meier curves for the top and bottom decile of score. The narrow range of ages and short time since inception means that UK Biobank subject curves are subject to greater uncertainty, particularly at each end, and only cover a shorter interval. E&W – England and Wales; PRS – polygenic risk score.

**Fig. S7:**
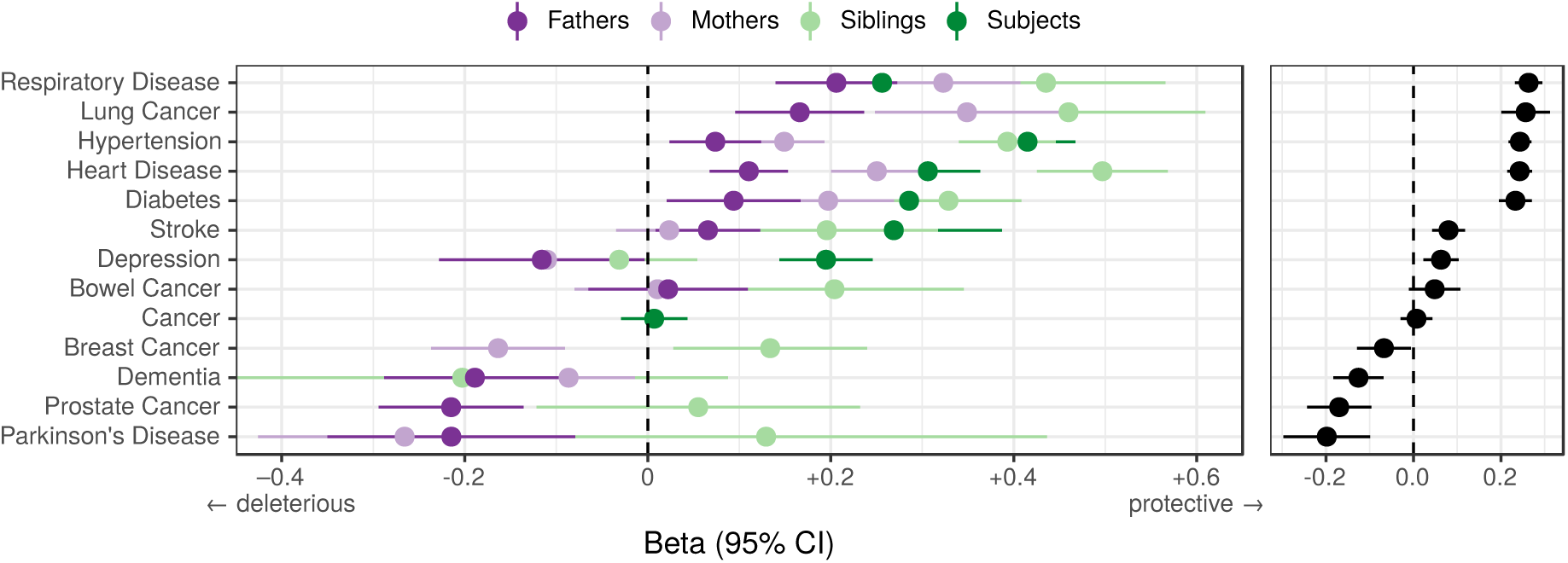
Associations between polygenic lifespan score and diseases of UK Biobank subjects and their kin. Logistic regression was performed on standardised polygenic survival score (all variants) and 21 disease traits reported by 24,059 Scottish and 29,815 English/Welsh out-of-sample individuals about themselves and their kin. Displayed here are inverse-variance meta-analysed estimates of the diseases for which multiple sources of data were available (i.e. parents and/or siblings; see Fig. S8 for all associations). “Cancer” is only in subjects, whilst the specific subtypes are analysed for kin. The left panel shows disease estimates for each kin separately; the right panel shows the combined estimate, with standard errors adjusted for correlation between family members. Diseases have been ordered by magnitude of effect size (combined estimate). Beta – log odds reduction ratio of disease per standard deviation of polygenic survival score, where a negative beta indicates a deleterious effect of score on disease prevalence (lifetime so far), and positive beta indicates a protective effect on disease. Effect sizes for first degree relative have been doubled. Cancer – Binary cancer phenotype (any cancer, yes / no).

**Fig. S8:**
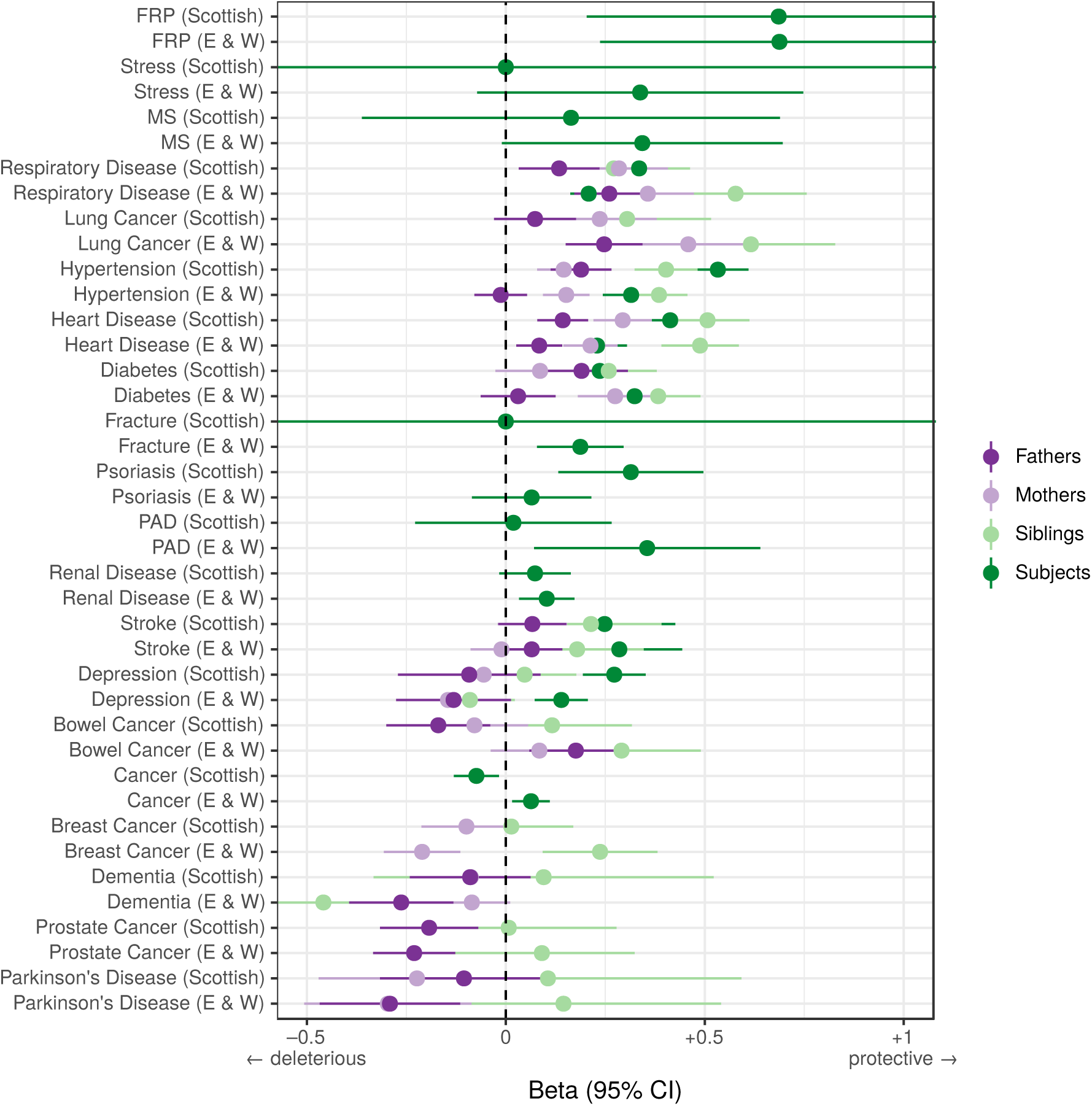
Associations between polygenic survival score and diseases of individuals and their kin from Scottish and English/Welsh subsamples of UK Biobank. Logistic regression was performed on standardised polygenic survival score (all variants) and 21 traits reported by 24,059 Scottish and 29,815 English/Welsh out-of-sample individuals about themselves and their kin. Diseases have been ordered by magnitude of effect size (meta-analysed between cohorts and kin). Beta – log odds reduction ratio of disease per standard deviation of polygenic survival score, where a negative beta indicates a deleterious effect of score on disease prevalence (lifetime so far), and positive beta indicates a protective effect on disease. Effect sizes for first degree relative have been doubled. Cancer – Binary cancer phenotype (yes / no), FRP – Female Reproductive Problems, MS – Multiple Sclerosis, PAD – Peripheral Artery Disease.

**Fig. S9:**
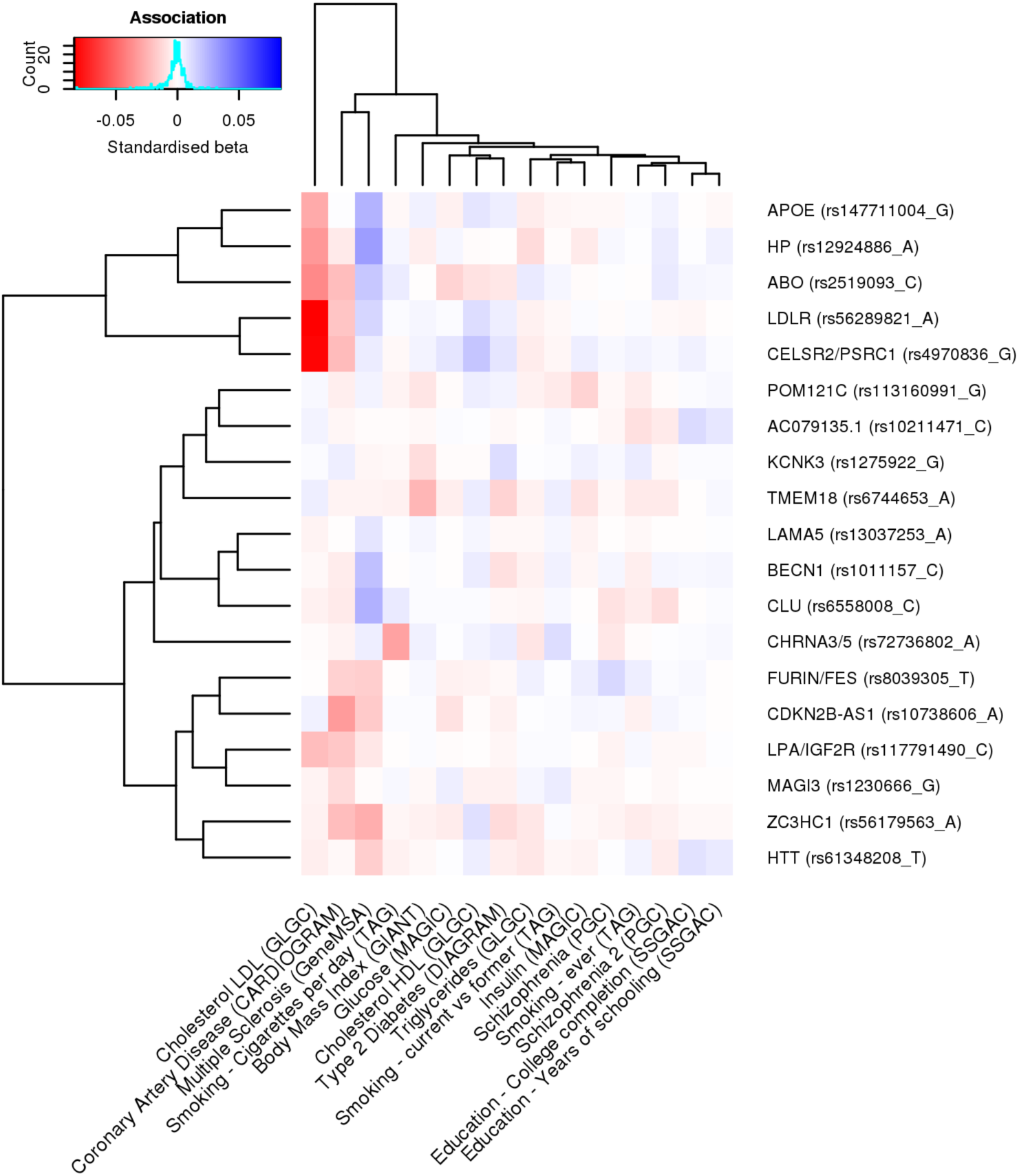
Heat map of the effect of 19 SNPs significant at 5e-8 on the risk factors in the iGWAS. We looked up the effects of lifespan protecting alleles identified by iGWAS in the consortium GWAMA for all risk factors significantly associated with lifespan in univariate analysis (for studies tested see Methods). We kept all traits univariately associated with lifespan to allow for the presence of potentially correlated traits, not significant in the multivariate analysis. In the iGWAS analysis, Z-scores (estimated effect divided by standard error) are used, but for comparison purposes, standardised betas (Z-score divided by square root of the sample size) were calculated for each risk factor at every SNP and represented in this figure. Both SNPs and traits were clustered for similarity. For example, we can see that almost all iGWAS alleles identified as protective for lifespan are exhibiting negative standardized betas in the coronary artery disease (CAD) association study, confirming the hypothesis that CAD is negatively affecting lifespan. We can also notice that some SNPs are strongly associated with some risk factors (APOE and LDR with lipids traits or CDKN2B-AS1 with CAD) and likely influence lifespan through their effect on these traits. However, some other SNPs (KCNK3 and HTT for example) are showing moderate effects on several risk factors and are probably affecting lifespan through pleiotropic effects.

**Table S1:** Descriptive statistics of the cohorts and lives analysed. Summary statistics for the 1,012,240 parental lifespans passing phenotypic QC (most notably, parent age > 40). In practice, fewer lives than these were analysed for some SNPs, as a SNP may not have passed QC in all cohorts (in particular LifeGen MAF > 1%). Ancestries in UK Biobank are self-declared, except in the case of Gen. British. Gen. British – Participants identified as genomically British by UK Biobank, based on their genomic profile. D+R - Discovery and replication cohorts combined. LifeGen – A consortium of 26 population cohorts of European Ancestry, with UK Biobank lives removed

| Study | Cohort | Ancestry | Parent | Dead | Count | Age |  |  |
| --- | --- | --- | --- | --- | --- | --- | --- | --- |
|  |  |  |  |  |  | Max | Mean | Min |
| UK Biobank | Discovery | Gen.British | Father | FALSE | 74,320 | 103 | 77.97 | 60 |
| UK Biobank | Discovery | Gen.British | Father | TRUE | 237,940 | 106 | 71.51 | 40 |
| UK Biobank | Discovery | Gen.British | Mother | FALSE | 129,357 | 105 | 78.60 | 60 |
| UK Biobank | Discovery | Gen.British | Mother | TRUE | 193,588 | 107 | 75.14 | 40 |
| UK Biobank | Replication | British | Father | FALSE | 4,087 | 106 | 78.55 | 60 |
| UK Biobank | Replication | British | Father | TRUE | 12,035 | 102 | 71.64 | 40 |
| UK Biobank | Replication | British | Mother | FALSE | 7,430 | 105 | 78.36 | 60 |
| UK Biobank | Replication | British | Mother | TRUE | 9,605 | 104 | 74.35 | 40 |
| UK Biobank | Replication | Irish | Father | FALSE | 1,799 | 100 | 77.56 | 60 |
| UK Biobank | Replication | Irish | Father | TRUE | 7,955 | 101 | 71.20 | 40 |
| UK Biobank | Replication | Irish | Mother | FALSE | 3,546 | 102 | 78.32 | 60 |
| UK Biobank | Replication | Irish | Mother | TRUE | 6,564 | 103 | 74.08 | 40 |
| UK Biobank | Replication | European | Father | FALSE | 523 | 101 | 76.28 | 60 |
| UK Biobank | Replication | European | Father | TRUE | 1,073 | 103 | 70.29 | 40 |
| UK Biobank | Replication | European | Mother | FALSE | 947 | 96 | 74.46 | 60 |
| UK Biobank | Replication | European | Mother | TRUE | 852 | 101 | 71.94 | 40 |
| LifeGen | Replication | European | Father | FALSE | 83,298 | 117 | 57.48 | 41 |
| LifeGen | Replication | European | Father | TRUE | 77,163 | 109 | 71.06 | 41 |
| LifeGen | Replication | European | Mother | FALSE | 97,794 | 117 | 59.55 | 41 |
| LifeGen | Replication | European | Mother | TRUE | 62,364 | 108 | 75.27 | 41 |
| Total | D+R | European | Father | FALSE | 164,027 | 117 | 67.57 | 41 |
| Total | D+R | European | Father | TRUE | 336,166 | 109 | 71.40 | 40 |
| Total | D+R | European | Mother | FALSE | 239,074 | 117 | 70.78 | 41 |
| Total | D+R | European | Mother | TRUE | 272,973 | 108 | 75.10 | 40 |
| Grand Total | D+R | European | Parents | ALL | 1,012,240 | 117 | 71.63 | 40 |

**Table S2:** Fourteen regions associate with lifespan at genome-wide significance in discovery and seven nominally replicate (P < 0.05) in a European population sample. At or near – Gene, cluster of genes, or cytogenetic band in close proximity to lead SNP; Chr – Chromosome, Position – Base-pair position on chromosome (build GRCh37); A1 – the effect allele, increasing lifespan; Freq1-Frequency of the A1 allele in the discovery population; Years – Years of lifespan gained for carrying one copy of the Al allele; Betal – the log_e_(protection ratio) for carrying one copy of A1 under additive dosage model which multiplied observed offspring genotype on parent effect by 2; SE – Standard Error; CES – Common effect size assumption of A1 across sexes; PDES – Potentially different effect size assumption of A1 across sexes; P – P value for the Wald test of association between imputed dosage and Cox model residual (replication P values are one sided for CES and direction agnostic for PDES). Loci reaching nominal significance (P < 0.05) in the replication cohort are bolded. Non-significant replication P values are greyed out.

| rsID | At or near | Chr | Position | A1 | Freq1 | Discovery |  |  |  |  | Replication |  |  |  |  |
| --- | --- | --- | --- | --- | --- | --- | --- | --- | --- | --- | --- | --- | --- | --- | --- |
|  |  |  |  |  |  | Beta1 | SE | Years | CES P | PDES P | Beta1 | SE | Years | CES P | PDES P |
| rs7528419 | CELSR2/PSRC1 | 1 | 109817192 | G | 0.222 | 0.0314 | 0.0055 | 0.31441 | 1.4E-08 | 2.6E-08 | 0.0012 | 0.0840 | 0.0121 | 0.4426 | 0.0565 |
| **rs34967069** | **HLA-DQA1** | **6** | **32591248** | **T** | **0.053** | **0.0577** | **0.0117** | **0.57693** | **7.5E-07** | **3.3E-08** | **0.0529** | **0.1669** | **0.5293** | **0.0008** | **0.0087** |
| **rs118039278** | **LPA** | **6** | **160985526** | **G** | **0.919** | **0.0737** | **0.0085** | **0.73662** | **4.0E-18** | **3.7E-18** | **0.0857** | **0.1548** | **0.8566** | **1.6E-08** | **1.3E-07** |
| rs10225529 | ARPC1 | 7 | 98963155 | C | 0.104 | 0.0419 | 0.0076 | 0.41888 | 3.2E-08 | 2.4E-07 | -0.0048 | 0.1141 | -0.0481 | 0.3368 | 0.6272 |
| rs7844965 | CLU | 8 | 27442064 | A | 0.232 | 0.0328 | 0.0055 | 0.32802 | 2.2E-09 | 4.4E-09 | 0.0057 | 0.0840 | 0.0569 | 0.7510 | 0.3500 |
| **rs1556516** | **CDKN2B-AS1** | **9** | **22100176** | **G** | **0.503** | **0.0246** | **0.0046** | **0.24638** | **9.1E-08** | **8.0E-10** | **0.0262** | **0.0703** | **0.2617** | **9.9E-05** | **0.0014** |
| rs10908903 | GADD45G | 9 | 92228559 | T | 0.532 | 0.0256 | 0.0047 | 0.2564 | 3.7E-08 | 1.3E-06 | 0.0065 | 0.0710 | 0.0653 | 0.1790 | 0.5779 |
| **rs11065979** | **ATXN2/BRAP** | **12** | **112059557** | **C** | **0.563** | **0.0338** | **0.0047** | **0.33797** | **3.9E-13** | **1.8E-13** | **0.0137** | **0.0731** | **0.1366** | **0.0307** | 0.1652 |
| rs143498116 | 13q21.33 | 13 | 71286100 | A | 0.994 | 0.1052 | 0.0320 | 1.05173 | 0.0010 | 2.4E-08 | 0.1209 | 0.9451 | 1.2094 | 0.1004 | 0.4474 |
| **rs72738786** | **CHRNA3/5** | **15** | **78828086** | **G** | **0.666** | **0.0474** | **0.0049** | **0.47408** | **3.0E-22** | **6.0E-26** | **0.0302** | **0.0747** | **0.3021** | **2.6E-05** | **0.0001** |
| **rs7177338** | **FURIN/FES** | **15** | **91428636** | **A** | **0.525** | **0.0266** | **0.0046** | **0.2658** | **9.1E-09** | **6.9E-08** | **0.0204** | **0.0724** | **0.2040** | **0.0024** | **0.0247** |
| **rs429358** | **APOE** | **19** | **45411941** | **T** | **0.844** | **0.1114** | **0.0063** | **1.11421** | **9.4E-69** | **8.8E-71** | **0.0890** | **0.1074** | **0.8897** | **5.9E-17** | **5.5E-16** |
| rs6108784 | C20orf187 | 20 | 10964366 | T | 0.585 | 0.0185 | 0.0047 | 0.18466 | 8.5E-05 | 3.3E-08 | 0.0086 | 0.0727 | 0.0857 | 0.1193 | 0.4176 |
| rs6011779 | CHRNA4 | 20 | 61984317 | T | 0.81 | 0.0328 | 0.0059 | 0.32835 | 2.3E-08 | 6.2E-07 | -0.0044 | 0.0999 | -0.0438 | 0.3307 | 0.8935 |

**Table S3:** Five candidate lifespan regions replicate nominally (P < 0.05) in our discovery+replication meta-analysis sample or European population sample. At or near – Gene, cluster of genes, or cytogenetic band in close proximity to lead SNP; Chr – Chromosome, Position – Base-pair position on chromosome (build GRCh37); A1 – the effect allele, increasing lifespan in discovery; Freq1-Frequency of the A1 allele in the replication sample, or if missing, the discovery sample; Sex – sex of the individuals or their parents used in the discovery and replication; Beta1 – the log,(protection ratio) for carrying one copy of A1 under additive dosage model, inferred for discovery (see Methods); SE – Standard Error, calculated from reported P value and inferred effect estimates for discovery, assuming a two-sided test; Years – Years of lifespan gained for carrying one copy of the A1 allele; P – P value reported by original study for discovery, one-sided P value for the Wald test association between imputed dosage and cox model residual for the replication. For discovery, except Pilling et al’s (6) SNPs, where we re-calculated effects directly from individual UKBB data ourselves, effects sizes have been converted to a common scale to enable comparison. Ref-reference ID of original study that identified the candidate SNP; Sample – independent sample used to replicate the results (a = UK Biobank Discovery+Replication Meta-analysis, b = LifeGen excluding UK Biobank). Loci showing nominal replication (P < 0.05) are bolded. Note, rs151091095 near USP2-AS1 is a proxy (r^2^ = 1.00) for rs139137459, the SNP reported by Pilling et al; rs113946246 near ANKRD20A9P is a proxy (r^2^ = 0.97) for rs2440012, the SNP reported by Zeng et al; no proxies could be found for 13:31871514_T_G.

| rsID | At or near | Chr | Position | A1 | Freq1 | Sex | Discovery |  |  |  | Study | Replication |  |  |  | Sample |
| --- | --- | --- | --- | --- | --- | --- | --- | --- | --- | --- | --- | --- | --- | --- | --- | --- |
|  |  |  |  |  |  |  | Beta1 | SE | Years | P |  | Beta1 | SE | Years | P |  |
| rs3800231 | FOXO3A | 6 | 108998266 | A | 0.285 | Both | 0.0499 | 0.0223 | 0.4989 | 0.0250 | Flachsbar et al. 2009 | 0.0174 | 0.0043 | 0.1745 | 2.2E-05 | a |
| rs2149954 | 5q33.3/EBF1 | 5 | 157820602 | T | 0.359 | Both | 0.0249 | 0.0044 | 0.2494 | 1.7E-08 | Deelen et al. 2014 | 0.0085 | 0.0040 | 0.0853 | 0.0167 | a |
| rs2069837 | IL6 | 7 | 22768027 | A | 0.067 | Both | 0.0716 | 0.0119 | 0.7160 | 1.8E-09 | Zeng et al. 2016 | -0.0074 | 0.0076 | -0.0743 | 0.8362 | a |
| rs113946246 | ANKRD20A9P | 13 | 19429318 | T | 0.822 | Both | 0.0735 | 0.0134 | 0.7351 | 3.7E-08 | Zeng et al. 2016 | 0.0008 | 0.0055 | 0.0078 | 0.4440 | a |
| rs6873545 | d3-GHR | 5 | 42631264 | C | 0.269 | Male | 0.1553 | 0.0603 | 1.5533 | 0.0100 | Ben-Avraham et al. 2017 | -0.0008 | 0.0056 | -0.0084 | 0.5597 | a |
| rs3764814 | USP42 | 7 | 6189780 | C | 0.929 | Both | 0.0565 | 0.0072 | 0.5650 | 5.0E-15 | Sebastiani et al. 2017 | 0.0046 | 0.0075 | 0.0463 | 0.2681 | a |
| rs7976168 | TMT2C | 12 | 83438559 | A | 0.667 | Both | 0.0282 | 0.0048 | 0.2820 | 4.0E-09 | Sebastiani et al. 2017 | 0.0049 | 0.0041 | 0.0494 | 0.1140 | a |
| rs146254978 | FGPT/TNNI3K | 1 | 74867799 | C | 0.059 | Both | 0.0393 | 0.0171 | 0.3929 | 0.0210 | Pilling et al. 2017 | 0.0051 | 0.0555 | 0.0507 | 0.4636 | b |
| rs1627804 | BEND3 | 6 | 107400428 | C | 0.666 | Both | 0.0185 | 0.0047 | 0.1854 | 1.1E-04 | Pilling et al. 2017 | 0.0092 | 0.0085 | 0.0921 | 0.1382 | b |
| rs151091095 | USP2-AS1 | 11 | 119289932 | G | 0.013 | Both | 0.0817 | 0.0255 | 0.8172 | 0.0013 | Pilling et al. 2017 | 0.1265 | 0.1004 | 1.2652 | 0.1038 | b |
| rs61978928 | PROX2 | 14 | 75321714 | C | 0.323 | Both | 0.0267 | 0.0048 | 0.2673 | 2.0E-08 | Pilling et al. 2017 | -0.0051 | 0.0085 | -0.0509 | 0.7258 | b |
| rs28926173 | MC2R | 18 | 13886719 | A | 0.050 | Both | 0.0462 | 0.0103 | 0.4622 | 6.4E-06 | Pilling et al. 2017 | 0.0376 | 0.0231 | 0.3761 | 0.0521 | b |
| rs3131621 | MICA/MICB | 6 | 31425499 | A | 0.682 | Male | 0.0283 | 0.0051 | 0.2828 | 3.6E-08 | Pilling et al. 2017 | 0.0203 | 0.0154 | 0.2032 | 0.0930 | b |
| rs13262617 | TOX | 8 | 59838133 | G | 0.035 | Male | 0.0824 | 0.0147 | 0.8242 | 3.1E-08 | Pilling et al. 2017 | 0.0230 | 0.0303 | 0.2305 | 0.2233 | b |
| rs61905747 | ZW10 | 11 | 113639842 | A | 0.806 | Male | 0.0365 | 0.0063 | 0.3654 | 5.5E-09 | Pilling et al. 2017 | 0.0298 | 0.0139 | 0.2977 | 0.0161 | b |
| rs74011415 | SEMA6D | 15 | 47660194 | G | 0.853 | Male | 0.0439 | 0.0077 | 0.4385 | 1.4E-08 | Pilling et al. 2017 | 0.0105 | 0.0167 | 0.1049 | 0.2646 | b |
| rs12461964 | EGLN2/CYP2A6 | 19 | 41341229 | A | 0.504 | Male | 0.0288 | 0.0050 | 0.2879 | 8.2E-09 | Pilling et al. 2017 | 0.0041 | 0.0139 | 0.0408 | 0.3844 | b |
| rs74444983 | EXOC3L2/MARK4 | 19 | 45745607 | T | 0.735 | Male | 0.0327 | 0.0057 | 0.3265 | 9.1E-09 | Pilling et al. 2017 | 0.0164 | 0.0130 | 0.1638 | 0.1032 | b |
| rs3130507 | PSORS1C3 | 6 | 31147476 | G | 0.750 | Female | 0.0400 | 0.0063 | 0.3998 | 2.1E-10 | Pilling et al. 2017 | 0.0292 | 0.0175 | 0.2919 | 0.0480 | b |
| 13:31871514_T_G | B3GALT1 | 13 | 31871514 | G | 0.005 | Female | 0.2532 | 0.0464 | 2.5324 | 4.7E-08 | Pilling et al. 2017 | NA | NA | NA | NA | b |
| rs61949650 | 13q21.31 | 13 | 64836488 | C | 0.066 | Female | 0.0649 | 0.0117 | 0.6492 | 2.9E-08 | Pilling et al. 2017 | 0.0404 | 0.0239 | 0.4037 | 0.0455 | b |

**Table S4:** LD-score regression intercepts for GWAS results. Regression intercepts (standard error) of the GWAS summary statistics as calculated by LD-score regression, using LD scores from on average 457,407 SNPs from the UK Biobank array. CES – Results under the assumption of common effect sizes across sexes, PDES – Results under the assumption of potentially different effect sizes across sexes.

| Cohort | CES | PDES |
| --- | --- | --- |
| Discovery | 1.0351 (0.0049) | 1.0256 (0.0046) |
| Replication | 1.056 (0.0073) | 1.0134 (0.0045) |
| Meta | 1.0542 (0.0057) | 1.0307 (0.0049) |

**Table S5:** Predicted candidate genes using SMR-HEIDI test and two blood eQTL datasets. All loci reaching genome-wide significance in discovery cohort GWAS, combined cohort GWAS, or iGWAS (Table 1), plus candidate loci d3-GHR, 5q33.3/EBF1, and FOXO3 were tested against eQTL data. Only genes that pass 5% false discovery rate threshold for the SMR test and P > 0.05 threshold for HEIDI test are listed for their corresponding loci. Westra and CAGE refer to the two eQTL studies in the blood tissue. Corresponding probes also pass the P > 0.05 threshold for the HEIDI test, i. e. the expressions of the particular genes possibly share causal variants with the lifespan GWAS signals. Chr - chromosome. At or near – nearby gene or cluster of genes to lead variant. SMR genes – genes prioritised by SMR within the given locus.

| rsID | Chr | Position | At or near | SMR Prioritized Genes | Westra |  | CAGE |  |
| --- | --- | --- | --- | --- | --- | --- | --- | --- |
|  |  |  |  |  | P SMR | P HEIDI | P SMR | P HEIDI |
| rs7528419 | 1 | 109817192 | CELSR2/PSRC1 | PSRC1 |  |  | 7.28E-08 | 2.98E-01 |
| rs10225529 | 7 | 98963155 | ARPC1 | ARPC1B | 1.71E-05 | 2.97E-01 |  |  |
| rs11065979 | 12 | 112059557 | ATXN2/BRAP | SH2B3 | 3.76E-04 | 6.02E-02 |  |  |
| rs72738786 | 15 | 78828086 | CHRNA3/5 | PSMA4 | 8.19E-10 | 2.24E-01 | 1.30E-05 | 1.08E-01 |
|  |  |  |  | PSMA4 | 3.06E-08 | 5.59E-01 |  |  |
| rs7177338 | 15 | 91428636 | FURIN/FES | FURIN/FES | 1.83E-06 | 5.50E-02 | 1.84E-04 | 6.21E-02 |
| rs28971796 | 4 | 49151982 | CHW43 | OCIAD1 | 8.51E-04 | 1.99E-01 |  |  |
| rs1011157 | 17 | 40960253 | BECN1 | BECN1 |  |  | 1.16E-06 | 1.55E-01 |
|  |  |  |  | ATP6V0A1 |  |  | 1.01E-04 | 9.47E-01 |
| rs142158911 | 19 | 11190534 | LDLR | KANK2 | 7.17E-04 | 5.66E-01 |  |  |
| rs3800231 | 6 | 108998266 | FOXO3 | SESN1 | 8.17E-04 | 2.64E-01 |  |  |

**Table S6:**
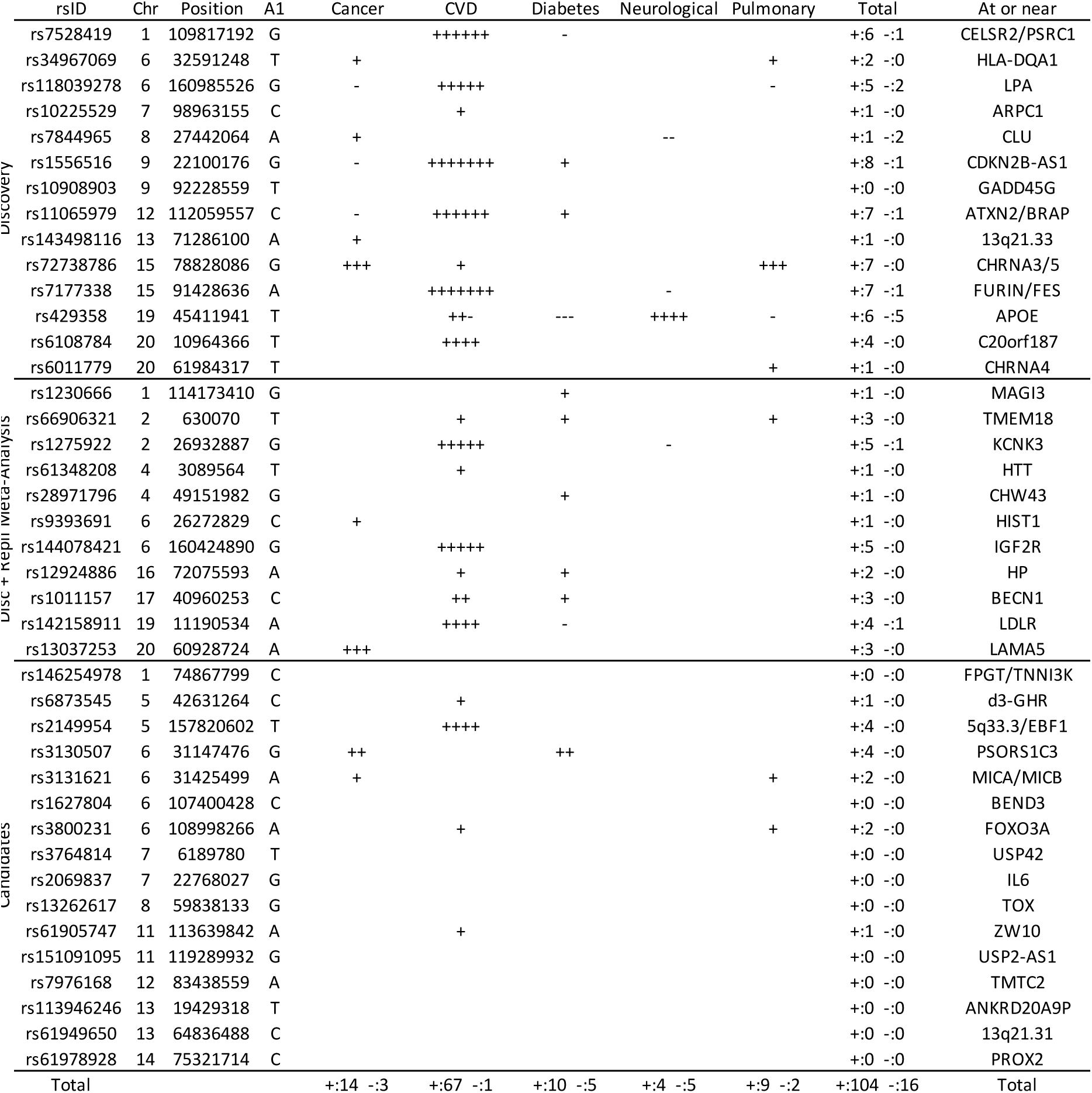
Significant associations of lifespan-protective variants (genome-wide significant in discovery or discovery+replication meta-analysis) and candidate variants with protection from 5 major disease categories in UK Biobank. Reported associations have been identified in 325,292 UK Biobank subjects, their siblings, or parents, at FDR 5%. Protective (+) and deleterious (−) associations of longevity increasing variant with a disease are listed within diseases categories. Chr – Chromosome; Position – Base-pair position (GRCh37); A1 – the effect allele, increasing lifespan; Count of protective and deleterious with: Cancer – cancer incidence, CVD – cardiovascular disease incidence, Diabetes – Type 2 diabetes incidence, Neurological – neurological disease incidence, Pulmonary – pulmonary disease incidence. Except for LAMA5 and HLA-DQA1, loci associating protectively with cancer specifically affect lung cancer. For the list of individual associations, see Table S17.

**Table S7:** Count of associations of lead SNPs with traits in PhenoScanner by broad disease category. Associations of lead lifespan SNPs, i.e. those identified in the discovery sample (top) and/or the combined discovery and replication sample (middle), and candidate lifespan SNPs (bottom), were retrieved from the PhenoScanner database. Trait associations include those with SNPs in high linkage disequilibrium with lead variants (r^2^ > 0.8) and were only reported if they passed a FDR 5% significance threshold. Grouping by broad disease categories was done as follows: CVD – Cardiovascular diseases and risk factors, such as myocardial infarction, aortic valve calcification, hypertension, and cholesterol and triglyceride levels (15 traits). IMMUNE – autoimmune and chronic inflammation disorders, such as type 1 diabetes, rheumatoid arthritis, multiple sclerosis, inflammatory bowel diseases, and autoimmune liver and thyroid disease (15 traits). PULMONARY – pulmonary function and disease (exc. cancer): asthma, chronic pulmonary obstructive disorder, respiratory function and airflow obstruction (4 traits). DIABETES – type 2 diabetes and risk factors including glucose, HbA1c, and insulin levels (4 traits). OBESITY – Anthropometric measures such as BMI, body fat percentage, waist/hip circumference, weight, and obesity (7 traits). NEURO – Neurological disorders, such as Alzheimer’s, Parkinson’s, and Huntington’s disease, as well as depression, smoking addiction, and neuroticism (14 traits). See Table S18 for a full list of traits and associations.

| Gene | CANCER | CVD | OBESITY | DIABETES | NEURO | IMMUNE | PULMONARY | TOTAL |
| --- | --- | --- | --- | --- | --- | --- | --- | --- |
| CELSR2/PSRC1 | 0 | 10 | 3 | 2 | 1 | 0 | 0 | 16 |
| HLA-DQA1 | 0 | 0 | 0 | 0 | 2 | 1 | 0 | 3 |
| LPA | 0 | 6 | 0 | 0 | 0 | 0 | 0 | 6 |
| ARPC1 | 0 | 0 | 1 | 1 | 0 | 0 | 0 | 2 |
| CLU | 0 | 1 | 1 | 1 | 1 | 0 | 0 | 4 |
| CDKN2B-AS1 | 0 | 5 | 0 | 1 | 0 | 0 | 0 | 6 |
| GADD45G | 0 | 2 | 3 | 0 | 0 | 0 | 0 | 5 |
| ATXN2/BRAP | 0 | 7 | 6 | 2 | 1 | 13 | 1 | 30 |
| 13q21.33 | 0 | 0 | 0 | 0 | 0 | 0 | 0 | 0 |
| CHRNA3/5 | 1 | 1 | 3 | 0 | 4 | 0 | 3 | 12 |
| FURIN/FES | 0 | 3 | 1 | 0 | 4 | 0 | 0 | 8 |
| APOE | 0 | 4 | 0 | 0 | 3 | 0 | 0 | 7 |
| C20orf187 | 0 | 2 | 1 | 0 | 1 | 0 | 0 | 4 |
| CHRNA4 | 0 | 0 | 0 | 0 | 0 | 0 | 0 | 0 |
| MAGI3 | 0 | 1 | 0 | 0 | 0 | 6 | 0 | 7 |
| TMEM18 | 0 | 1 | 7 | 1 | 0 | 0 | 0 | 9 |
| KCNK3 | 0 | 2 | 5 | 0 | 0 | 0 | 0 | 7 |
| HTT | 0 | 2 | 1 | 0 | 2 | 2 | 0 | 7 |
| CHW43 | 0 | 0 | 0 | 0 | 0 | 0 | 0 | 0 |
| HIST1 | 0 | 3 | 3 | 0 | 3 | 2 | 0 | 11 |
| IGF2R | 0 | 4 | 0 | 0 | 0 | 1 | 0 | 5 |
| HP | 0 | 5 | 2 | 0 | 1 | 0 | 0 | 8 |
| BECN1 | 0 | 0 | 0 | 0 | 0 | 0 | 0 | 0 |
| LDLR | 0 | 7 | 0 | 0 | 0 | 0 | 0 | 7 |
| LAMA5 | 0 | 0 | 0 | 0 | 0 | 0 | 0 | 0 |
| FPGT/TNNI3K | 0 | 0 | 0 | 0 | 0 | 0 | 0 | 0 |
| d3-GHR | 0 | 0 | 1 | 0 | 0 | 0 | 0 | 1 |
| 5q33.3/EBF1 | 0 | 3 | 0 | 0 | 0 | 0 | 0 | 3 |
| MICA/MICB | 0 | 2 | 1 | 0 | 1 | 5 | 1 | 10 |
| PSORS1C3 | 0 | 0 | 0 | 0 | 1 | 2 | 0 | 3 |
| BEND3 | 1 | 0 | 0 | 0 | 0 | 0 | 0 | 1 |
| FOXO3A | 0 | 2 | 6 | 0 | 3 | 0 | 1 | 12 |
| USP42 | 0 | 0 | 0 | 0 | 0 | 0 | 0 | 0 |
| IL6 | 0 | 0 | 0 | 0 | 1 | 1 | 0 | 2 |
| TOX | 0 | 2 | 0 | 0 | 0 | 0 | 0 | 2 |
| ZW10 | 0 | 0 | 1 | 0 | 0 | 0 | 0 | 1 |
| USP2-AS1 | 0 | 0 | 0 | 0 | 0 | 0 | 0 | 0 |
| TMTC2 | 0 | 0 | 1 | 0 | 0 | 0 | 0 | 1 |
| ANKRD20A9P | 0 | 0 | 0 | 0 | 0 | 0 | 0 | 0 |
| 13q21.31 | 0 | 0 | 0 | 0 | 0 | 0 | 0 | 0 |
| PROX2 | 0 | 2 | 0 | 0 | 2 | 0 | 0 | 4 |
| SEMA6D | 0 | 0 | 0 | 0 | 0 | 0 | 0 | 0 |
| MC2R | 0 | 1 | 0 | 0 | 0 | 0 | 0 | 1 |
| EGLN2/CYP2A6 | 0 | 0 | 1 | 0 | 2 | 0 | 0 | 3 |
| EXOC3L2/MARK4 | 0 | 2 | 0 | 0 | 0 | 0 | 0 | 2 |
| TOTAL | 2 | 80 | 48 | 8 | 33 | 33 | 6 | 210 |

**Table S8:** Gene sets identified as enriched by VEGAS (FDR 5%), and corresponding results from DEPICT and PASCAL. Pathway ID – Gene ontology identifier or VEGAS ID number of the pathway; VEGAS Name – Name of the pathway in VEGAS; VEGAS Q – Empirical P value from VEGAS adjusted for multiple testing using Benjamini-Hochberg correction; Nominal P – Uncorrected P value obtained for enrichment of the pathway using each method; DEPICT GW: DEPICT analysis run on genome-wide significant variants (P < 5×10-8); DEPICT Sug – DEPICT analysis run on variants passing suggestive significance (P < 1×10-5); PASCAL – PASCAL analysis using dichotomised gene set from DEPICT. Grey boxes indicate there was no gene set in DEPICT matching the VEGAS gene set. Green boxes highlight nominally significant P values. P values remaining significant after correcting for multiple comparisons with VEGAS are listed in bold.

| Pathway ID | Vegas Name | VEGAS Q | Nominal P |  |  |
| --- | --- | --- | --- | --- | --- |
|  |  |  | DEPICT GW | DEPICT Sug | PASCAL |
| GO:0032994 | protein-lipid complex | 0.0024 | 0.4007 | 0.1129 | 0.0179 |
| GO:0034358 | plasma lipoprotein particle | 0.0024 | 0.4007 | 0.1129 | 0.0177 |
| HSA-174824 | lipoprotein metabolism | 0.0024 | 0.3872 | 0.0640 | 0.0829 |
| GO:1990777 | lipoprotein particle | 0.0024 |  |  |  |
| GO:0034362 | low-density lipoprotein particle | 0.0039 | 0.3886 | 0.1151 | 0.0367 |
| GO:0099572 | postsynaptic specialization | 0.0065 |  |  |  |
| GO:0043691 | reverse cholesterol transport | 0.0070 | 0.5256 | 0.0265 | 0.1451 |
| GO:0014069 | postsynaptic density | 0.0097 | 0.9199 | 0.9629 | **0.0014** |
| GO:0030301 | cholesterol transport | 0.0117 | 0.4263 | 0.1479 | 0.8495 |
| GO:0070328 | triglyceride homeostasis | 0.0117 | 0.4761 | 0.7302 | 0.7190 |
| GO:0071813 | lipoprotein particle binding | 0.0123 | 0.1612 | 0.1106 | 0.0206 |
| GO:0015918 | sterol transport | 0.0123 | 0.4783 | 0.1716 | 0.8408 |
| GO:0034185 | apolipoprotein binding | 0.0123 | 0.0413 | 0.0557 | 0.0107 |
| GO:0034364 | high-density lipoprotein particle | 0.0123 | 0.3213 | 0.1674 | 0.0115 |
| GO:0055090 | acylglycerol homeostasis | 0.0123 |  |  |  |
| GO:0060076 | excitatory synapse | 0.0123 | 0.8615 | 0.9116 | 0.0271 |
| GO:0060627 | regulation of vesicle-mediated transport | 0.0123 | 0.4993 | 0.8919 | 0.8416 |
| GO:0071814 | protein-lipid complex binding | 0.0123 | 0.1612 | 0.1106 | 0.0209 |
| GO:0098794 | postsynapse | 0.0123 |  |  |  |
| GO:0034369 | plasma lipoprotein particle remodeling | 0.0130 | 0.7773 | 0.1237 | 0.0727 |
| GO:0097006 | regulation of plasma lipoprotein particle levels | 0.0130 | 0.5828 | 0.0579 | 0.0458 |
| GO:0015850 | organic hydroxy compound transport | 0.0146 | 0.5181 | 0.6117 | 0.3329 |
| GO:0034368 | protein-lipid complex remodeling | 0.0161 | 0.7773 | 0.1237 | 0.0728 |
| GO:0042632 | cholesterol homeostasis | 0.0170 | 0.1629 | 0.1931 | 0.0943 |
| GO:0055092 | sterol homeostasis | 0.0187 | 0.1629 | 0.1931 | 0.0956 |
| GO:0034367 | macromolecular complex remodeling | 0.0216 | 0.7773 | 0.1237 | 0.0726 |
| GO:1902991 | regulation of amyloid precursor protein catabolic process | 0.0216 |  |  |  |
| GO:0048261 | negative regulation of receptor-mediated endocytosis | 0.0285 |  |  |  |
| 138068 | signaling events mediated by stem cell factor receptor (c-kit) | 0.0302 |  |  |  |
| GO:0042157 | lipoprotein metabolic process | 0.0357 | 0.9639 | 0.2099 | 0.3124 |
| GO:0030425 | dendrite | 0.0377 | 0.5113 | 0.9234 | **0.0002** |
| GO:0006897 | endocytosis | 0.0426 | 0.7422 | 0.6595 | 0.2804 |
| GO:0071827 | plasma lipoprotein particle organization | 0.0443 | 0.6279 | 0.0703 | 0.0973 |

**Table S9:** Bayesian GWAS - Multivariate effect estimates for the 16 traits chosen by the AIC based stepwise model selection. The multivariate MR identified 16 traits (58 tested, see McDaid et al, 2017 for an exhaustive list) with significant causal effect on lifespan and used the effect estimates to create the prior assumption of the expected effect size of each variant on lifespan, in the (Bayesian) iGWAS. Effect Estimate – the estimated effect of standardized trait on standardized lifespan, in multivariate model. SE – the standard error of the estimated effect, in multivariate model. P – the P value (two sided) from MR, for testing association between standardized trait and standardized lifespan, in multivariate model.

| Trait | Causal Effect Estimate | SE | P |
| --- | --- | --- | --- |
| Cholesterol LDL (GLGC) | -0.139 | 0.007 | 1.25E-81 |
| Education level, years of schooling (SSGAC) | 0.218 | 0.012 | 3.62E-75 |
| Body mass index (GIANT) | -0.146 | 0.012 | 1.55E-31 |
| Coronary Artery Disease (CARDIoGRAM) | -0.253 | 0.030 | 1.19E-16 |
| Cholesterol HDL (GLGC) | 0.048 | 0.008 | 4.18E-10 |
| Smoking, cigarettes per day (TAG) | -0.437 | 0.055 | 2.31E-15 |
| Triglycerides (GLGC) | -0.029 | 0.009 | 2.05E-03 |
| Type 2 diabetes (DIAGRAM) | -0.080 | 0.021 | 1.81E-04 |
| Smoking, current vs former (TAG) | 0.215 | 0.078 | 5.86E-03 |
| Schizophrenia (PGC) | -0.031 | 0.011 | 3.74E-03 |
| Glucose (MAGIC) | -0.070 | 0.025 | 5.88E-03 |
| Smoking, ever (TAG) | -0.227 | 0.071 | 1.30E-03 |
| Insulin (MAGIC) | -0.191 | 0.066 | 3.62E-03 |
| Multiple Sclerosis (GeneMSA) | -0.083 | 0.038 | 2.78E-02 |
| Schizophrenia 2 (PGC) | -0.127 | 0.061 | 3.94E-02 |
| Education level, college completion (SSGAC) | 0.080 | 0.041 | 4.89E-02 |

**Table S10: 82 SNPs significantly associated with lifespan at 1% FDR and the SNP’s associations with risk factors.** [see Supplementary Information Excel File] Bayesian iGWAS was performed using observed association results from combined GWAS (discovery and replication sample) and prior was based on 16 traits selected by AIC based stepwise model selection. Bayes Factor were calculated to compare effect estimates observed in the conventional GWAS to the prior effect computed. Empirical P values were assigned using a permutation approach and further corrected for multiple testing using Benjamini-Hochberg correction. Chr – Chromosome, Position – Base-pair position on chromosome (GRCh37), A1 – Effect Allele, Freq1 – Frequency of the A1 allele (from conventional GWAS), Beta1(from conventional GWAS), SE – Standard Error of Beta1, Years – Years of lifespan gained for carrying one copy of the A1 allele (from conventional GWAS), P – P value (from conventional GWAS), PriorEffect – Prior effect estimate calculated from the summary statistics data for the 16 risk factors identified, PriorSE – Standard Error of the prior effect estimate, LogBF – Log of the observed Bayes Factor, P_BF – Empirical P value from a permutation approach for the log Bayes Factor. Final columns show the P value of each SNP in the studies used to calculate the prior, if the P value is significant after Bonferroni multiple testing correction (P < 3.81×10^-5^, 82*16 tests) the cell is shaded green. Counts of these significant associations by SNP/trait are shown in the final column/row.

**Table S11: Replication of lead SNPs associating with lifespan using published longevity GWaS** [See Supplementary Information Excel File] At or near – gene, cluster of genes, or cytogenetic band near lead SNP; Proxy – the rsID of the nearest (r2) SNP reported by Deelen et al.; Chr – Chromosome; Position – Base-pair position (GRCh37); A1 – the effect allele, A0 – the reference allele, Freq1 – the frequency of A1 allele; Beta1 – the log hazard ratio (in self) for a carrier of 1 copy of A1; SE – standard error; P – P value for test of association between proxy and lifespan (for IVM replication this is one sided); Discovery – the combined GWAS of UKBB genomically British, UKBB other and LifeGen; Replication – the GWAMAs of Deelen et al (15), Broer et al (10) and Walter et al (28), recalibrated (using APOE) to log hazard ratios and then combined using inverse-variance meta-analysis; Alpha – the ratio of effect size in replication to discovery (note as this was calibrated on APOE, that result was necessarily 1).

**Table S12: Predicted causal genes using SMR-HEIDI test and expression QTLs** [see Supplementary Information Excel File] 48 tissues of the GTEx project were analysed by taking the significant eQTL signals. Only genes that pass 5% false discovery rate threshold for the SMR test and P > 0.05 threshold for HEIDI test are listed for their corresponding loci. Chr – chromosome, Position – Base-pair position (GRCh37), At or near – Nearest gene, cluster of genes, or cytogenetic band to lead SNP.

**Table S13: Predicted causal CpG probes using SMR-HEIDI and methylation QTLs** [see Supplementary Information Excel File] Methylation QTLs from blood tissue were analysed. Only CpG probes that pass 5% false discovery rate threshold for the SMR test and P > 0.05 threshold for HEIDI test are listed for their corresponding loci. Chr – chromosome, (Probe) Position – Base-pair position (GRCh37), At or near – Nearest gene, cluster of genes, or cytogenetic band to lead SNP.

**Table S14:** Evidence of allelic heterogeneity of the lifespan loci via identification of secondary associations using SOJO. SOJO maps additional variants of each locus besides the top variant by implementing a LASSO regression across the locus. R-squared (Top variant): the captured narrow-sense heritability by the top variant of each locus. R-squared (SOJO): out-of-sample prediction R-squared achieved in the replication cohort, i.e. the captured narrow-sense heritability of each locus by the polygenic score across multiple variants within the same locus; R-squared is on a non-intuitive scale here, as the phenotype is martingale residuals in the replication cohort, where a substantial proportion of the parents are still alive. It is therefore much lower than the proportion of lifespan variance explained for a set of subjects that are all dead. Nonetheless, the ratios of R-squared, which allow allelic heterogeneity to be assessed, remain valid. At or near – the gene, cluster of genes, or cytogenetic band in close proximity to the lead variant; N SOJO – The number of variants selected by SOJO, maximum is set to be 30; Ratio – The ratio of out of sample R-squared between using SOJO variants and top variant. A large ratio together with large N SOJO indicate there is higher allelic heterogeneity at the locus.

| Lead Variant | At or near | R-squared<br>(Lead Variant) | R-squared<br>(SOJO) | N SOJO | Ratio |
| --- | --- | --- | --- | --- | --- |
| rs7528419 | CELSR2/PSRC1 | 6.65E-08 | 4.55E-06 | 30 | 68.50 |
| rs34967069 | HLA-DQA1 | 6.33E-06 | 3.37E-05 | 29 | 5.32 |
| rs118039278 | LPA | 7.07E-05 | 1.89E-04 | 15 | 2.67 |
| rs10225529 | ARPC1 | 5.26E-07 | 2.04E-05 | 23 | 38.80 |
| rs7844965 | CLU | 1.42E-06 | 4.02E-06 | 11 | 2.84 |
| rs1556516 | CDKN2B-AS1 | 4.54E-05 | 4.61E-05 | 2 | 1.01 |
| rs10908903 | GADD45G | 2.70E-06 | 3.32E-06 | 22 | 1.23 |
| rs11065979 | ATXN2/BRAP | 1.07E-05 | 1.14E-05 | 2 | 1.05 |
| rs143498116 | 13q21.33 | 1.65E-06 | 2.55E-05 | 21 | 15.50 |
| rs72738786 | CHRNA3/5 | 5.27E-05 | 7.29E-05 | 30 | 1.39 |
| rs7177338 | FURIN/FES | 2.48E-05 | 8.21E-05 | 29 | 3.30 |
| rs429358 | APOE | 2.04E-04 | 2.63E-04 | 30 | 1.29 |
| rs6108784 | C20orf187 | 4.54E-06 | 2.33E-05 | 11 | 5.12 |
| rs6011779 | CHRNA4 | 5.09E-07 | 3.23E-06 | 8 | 6.36 |
| rs1230666 | MAGI3 | 1.66E-05 | 1.74E-05 | 2 | 1.05 |
| rs66906321 | TMEM18 | 5.66E-09 | 2.40E-05 | 23 | 4235.00 |
| rs1275922 | KCNK3 | 3.25E-05 | 4.17E-05 | 30 | 1.28 |
| rs61348208 | HTT | 3.39E-05 | 4.52E-05 | 3 | 1.33 |
| rs28971796 | CHW43 | 5.69E-06 | 6.02E-06 | 2 | 1.06 |
| rs9393691 | HIST1 | 1.43E-05 | 4.38E-05 | 18 | 3.07 |
| rs144078421 | IGF2R | 9.78E-06 | 5.52E-05 | 17 | 5.63 |
| rs12924886 | HP | 6.14E-05 | 1.34E-04 | 3 | 2.18 |
| rs1011157 | BECN1 | 9.60E-06 | 4.37E-05 | 30 | 4.56 |
| rs142158911 | LDLR | 5.05E-05 | 5.65E-05 | 6 | 1.12 |
| rs13037253 | LAMA5 | 1.16E-05 | 1.18E-05 | 2 | 1.01 |
| rs10211471 | AC079135.1 | 3.63E-05 | 5.33E-05 | 5 | 1.47 |
| rs113160991 | POM121C | 1.22E-05 | 1.23E-05 | 20 | 1.01 |
| rs56179563 | ZC3HC1 | 5.27E-05 | 6.64E-05 | 30 | 1.26 |
| rs2519093 | ABO | 1.66E-05 | 1.66E-05 | 1 | 1.00 |
| Total |  | 7.90E-04 | 1.41E-03 | 455 | 1.79 |

**Table S15: Detailed results of the fine-mapping analysis by SOJO** [see Supplementary Information Excel File] SOJO maps additional variants of each locus besides the top variant by implementing a LASSO regression across the locus. This table expands the results of Table S14. Each additional variant at each locus is reported. R-squared is on a non-intuitive scale here, as the phenotype is martingale residuals in the replication cohort, where a substantial proportion of the parents are still alive. It is therefore much lower than the proportion of lifespan variance explained for a set of subjects that are all dead. Nonetheless, the ratios of R-squared, which allow allelic heterogeneity to be assessed, remain valid. R-squared (Top variant) – the captured narrow-sense heritability by the top variant of each locus. R-squared (SOJO) – out-of-sample prediction R-squared achieved in the replication cohort, i.e. the captured narrow-sense heritability of each locus by the polygenic score across multiple variants within the same locus. RA – reference allele. EA(F) – effective allele (frequency). r – LD correlation with the top variant in each locus.

**Table S16: Grouping of UK Biobank disease codes into diseases and major disease categories** [see Supplementary Information Excel File] UKBB phenotypes included 29 self-reported non-cancer disease fields for the participants and each of their parents, which included 474 integer-value coded diseases. These 474 diseases were aggregated and meta-analysed into four major mortality-increasing disease groups, namely CVD, diabetes, neurological and pulmonary disorders. Cancer was the fifth major disease group and was coded as either the occurrence or absence of cancers instances throughout the participant’s lifetime. Codes not shown were excluded from the analysis.

**Table S17: Full list of associations of lead SNPs with subject, sibling, and parental diseases in UK Biobank** [see Supplementary Information Excel File] Disease associations have been identified in 325,292 UK Biobank subjects, their siblings, or parents, at FDR 5%. At or near – Gene, cluster of genes, or cytogenetic band in close proximity to lead variant. A1 – Longevity allele of lead SNP. Trait – Disease reported by UK Biobank subject (kin). N – Number of individuals tested. Cases – Number of reported individuals or kin carrying the disease. Beta – log OR of NOT carrying the disease (i.e. positive beta indicates the longevity SNP protects from disease). SE – Standard Error. P – Two-sided P value. Q – Benjamin-Hochberg adjusted P value

**Table S18: Full list of associations of lead SNPs with traits in PhenoScanner by broad disease category** [see Supplementary Information Excel File] Associations of lead lifespan SNPs identified in the discovery sample and/or the combined discovery and replication sample, and candidate lifespan SNPs, were retrieved from the PhenoScanner database. Trait associations include those with SNPs in high linkage disequilibrium with lead variants (r^2^Ϡ.8) and were only reported if they passed a FDR 5% significance threshold. Gene – Nearest gene or cluster of genes to lead variant, rsID – SNP identifier of lead or proxy SNP, Alleles – Effect allele and non-effect allele, matched to the lead SNP alleles, r2 – correlation coefficient between lead and proxy SNP, Trait – trait name reported by PhenoScanner, P – P value of association, Q – Benjamini-Hochberg FDR adjusted P value, PMID – PubMed identification number of study reporting the association, Category – Disease category the association has been assigned to.

**Table S19: List of genome-wide significant disease variants, their association with disease in UK Biobank and their lifespan variance explained** [see Supplementary Information Excel File] Genome-wide significant disease SNPs from the GWAS catalog are listed with the amount of lifespan variance explained (LVE), with disease-protective alleles signed positively when increasing lifespan and signed negatively when decreasing lifespan. SNPs with limited evidence of an effect on lifespan are greyed out: an FDR cut-off of 1.55% is applied simultaneously across all diseases, allowing for 1 false positive among all significant SNPs. Secondary pleiotropic SNPs (i.e. those associating strongly with another one of the diseases, as assessed by PheWAS in UK Biobank) are coloured, as less relevant to the disease in question. Of these, turquoise SNPs show one or more alternative disease associations in the same direction and at least twice as strong (double Z statistic) as the principal disease, while brown SNPs show one or more significant associations with alternative disease in the opposite direction that explains the negative association of the disease-protective SNP with lifespan. At or near – Gene, cluster of genes, or cytogenetic band in close proximity to lead variant. Chr – Chromosome. Position – Base-pair position on chromosome (build GRCh37). A1 – Allele protecting from disease or disease risk factors. Freq1 – Frequency of the disease-protective allele in the discovery+replication sample. Years – Years of lifespan gained for carrying one copy of the A1 allele. P – P value for association with lifespan under CES assumption (left), P value for genome-wide significant association with disease as reported in the GWAS catalog (right). Q – Benjamini-Hochberg FDR-corrected P value for association with lifespan. LVE – Lifespan variance explained, signed positively when A1 increases lifespan and negative when A1 decreases lifespan. Pleiotropic – SNP shows evidence of pleiotropy, see definition above. Trait – Disease trait reported in GWAS catalog. Beta1 – log OR for having the reported disease, or unit increase in risk factors associated with disease, per copy of A1 allele. PMID – PubMed identification number of the study reporting the disease association. Z estimates – Z statistic for association with disease in unrelated, Gen. British UK Biobank samples. Missing statistics indicate the SNP is not present in the CES meta-analysis summary statistics and its LVE has been imputed from the closest proxy (min. r^2^>0.9) or proxies if equally close.

**Table S20: Sex and age stratified effects on survival for 49 lifespan increasing variants** [see Supplementary Information Excel File] At or near – Gene, cluster of genes, or cytogenetic band in close proximity to lead variant. Variant – rsID, longevity allele. Parent – Parent. Age range – Lower limit to upper limit of age in analysis. N – Number of lives used for the analysis (e.g. a parent aged 55 contributed to analysis of 40-50 and 50-60, but not 60-70). Deaths – Number of deaths within the age range. Beta – log_e_(protection ratio) for 1 copy of effect allele in self in the age band (i.e. 2 x observed due to kin cohort method). SE – Standard error. Z – Test statistic for test of H_0_ P – P value of two sided test of association.

**Table S21: Effect sizes of sex and age moderators within fixed-effects with moderators’ model of longevity alleles for 49 SNPs** [see Supplementary Information Excel File] At or near – Gene, cluster of genes, or cytogenetic band in close proximity to lead variant. Variant – rsID, longevity allele. Beta – Moderator effect estimate of sex (categorical variable, being male) or age (ordinal variable, mean age in age band) on lead SNP effect on lifespan. SE – Standard error. P – P value for association of SNP lifespan effect size with age or sex. Q – Benjamini-Hochberg FDR-corrected P value. Bolded lines contain sex or age-specific effects passing a 5% FDR threshold.

**Table S22: Cell types enriched for lifespan heritability identified by stratified LD-score regression** [see Supplementary Information Excel File] Name – Default tissue or cell-type names from stratified LD-score regression data. Beta – regression coefficient fitting baseline model and cell-type specific LD scores. SE – Standard Error. P – Two-sided P value for regression coefficient. Q – Benjamini-Hochberg FDR corrected P value.

**Table S23: Putative lifespan pathways highlighted by VEGAS2Pathway gene set enrichment analysis** [see Supplementary Information Excel File] Pathway – Reference number and name of gene set. nGenesMapped – Number of genes in the pathway tagged by SNPs. nGenesUsed – Number of genes after pruning. nSamples – Number of permutations used to calculate observed P value. ObservedP – Unadjusted P value for pathway enrichment. empiricalP – Observed P value adjusted for pathway size. Q – Benjamini-Hochberg-adjusted empirical P value. Genes – Genes present in the gene set

**Table S24:** eQTL SNPs associated with lifespan are for genes whose expression varies with age. We identified SNPs in our GWAS (discovery plus replication combined CES) that were also eQTLs i.e. associated with the expression of at least one gene in a dataset provided to us by the eQTLGen Consortium. A total of 2967 eQTLs after distance pruning (500kb) were present, of which 500 were associated with genes differentially expressed with age(101). We used Fisher’s exact test to determine, amongst the set of eQTLs, if SNPs which were associated with lifespan (at varying thresholds of statistical significance) were enriched for SNPs associated with genes whose expression is age-related. Threshold – P value threshold for lifespan association. #eQTLs (All) – number of independent eQTLs passing the significance threshold. #eQTLs (Ageing) – number of independent eQTLs for genes differentially expressed with age passing the significance threshold. OR – Odds ratio. P – P value for Fisher’s exact test.

| Threshold | # eQTLs |  | OR | P |
| --- | --- | --- | --- | --- |
|  | All | Ageing |  |  |
| 5E-8 | 16 | 7 | 3.45 | 0.0176 |
| 5E-7 | 28 | 10 | 2.47 | 0.0266 |
| 5E-6 | 63 | 21 | 2.25 | 0.0046 |
| 5E-5 | 206 | 63 | 2.07 | 1.4E-05 |
| 5E-4 | 654 | 168 | 1.78 | 1.7E-07 |
| 0.005 | 1685 | 386 | 2.34 | 1.1E-14 |
| 0.05 | 2340 | 475 | 3.38 | 6.8E-11 |
| 1 | 2967 | 500 | - | - |

**Table S25:** Polygenic survival scores in independent samples are most predictive when including all markers. A polygenic risk score was made for each subject using GWAS results that did not include the subject sets under consideration. Parent survival information (age and alive/dead status) was used to test the association between survival and several polygenic risk scores with different P value thresholds. Sample – Out-of-sample subsets of UK Biobank individuals used for PGRS association. N – Number of reported parental lifespans by sample individuals. Deaths – Number of reported parental deaths by sample individuals. Threshold – Criteria for SNPs to be included in the polygenic score. Beta – Log_e_(protection ratio) per standard deviation of polygenic score, doubled to reflect the effect of the score on offspring survival. SE – standard error of the effect estimate. Mean Years – Mean years of life gained per standard deviation in PGRS. P – P value of the predicted effect of the polygenic score on lifespan.

| Sample | Parent | N | Deaths | Threshold | Beta | SE | Mean Years | P |
| --- | --- | --- | --- | --- | --- | --- | --- | --- |
| Scottish | Fathers | 23,071 | 18,255 | P < 5E-8 | 0.08 | 0.01 | 0.81 | 5.2E-08 |
|  |  |  |  | P < 1E-6 | 0.08 | 0.01 | 0.78 | 1.5E-07 |
|  |  |  |  | P < 1E-4 | 0.08 | 0.01 | 0.78 | 1.5E-07 |
|  |  |  |  | P < 0.05 | 0.08 | 0.01 | 0.82 | 3.9E-08 |
|  |  |  |  | P < 1 | 0.10 | 0.02 | 0.99 | 4.4E-11 |
|  | Mothers | 23,865 | 14,941 | P < 5E-8 | 0.08 | 0.02 | 0.84 | 3.1E-07 |
|  |  |  |  | P < 1E-6 | 0.09 | 0.02 | 0.91 | 2.7E-08 |
|  |  |  |  | P < 1E-4 | 0.11 | 0.02 | 1.12 | 1.3E-11 |
|  |  |  |  | P < 0.05 | 0.10 | 0.02 | 1.02 | 5.8E-10 |
|  |  |  |  | P < 1 | 0.12 | 0.02 | 1.19 | 5.8E-13 |
| English & Welsh (out-of-sample subset) | Fathers | 28,532 | 21,699 | P < 5E-8 | 0.07 | 0.01 | 0.73 | 8.3E-08 |
|  |  |  |  | P < 1E-6 | 0.08 | 0.01 | 0.82 | 1.7E-09 |
|  |  |  |  | P < 1E-4 | 0.08 | 0.01 | 0.84 | 6.8E-10 |
|  |  |  |  | P < 0.05 | 0.09 | 0.01 | 0.94 | 7.0E-12 |
|  |  |  |  | P < 1 | 0.12 | 0.01 | 1.21 | 2.9E-18 |
|  | Mothers | 29,538 | 17,648 | P < 5E-8 | 0.07 | 0.02 | 0.74 | 1.1E-06 |
|  |  |  |  | P < 1E-6 | 0.07 | 0.02 | 0.70 | 4.0E-06 |
|  |  |  |  | P < 1E-4 | 0.09 | 0.02 | 0.89 | 3.8E-09 |
|  |  |  |  | P < 0.05 | 0.12 | 0.02 | 1.18 | 8.2E-15 |
|  |  |  |  | P < 1 | 0.15 | 0.02 | 1.48 | 1.9E-22 |

**Table S26:**
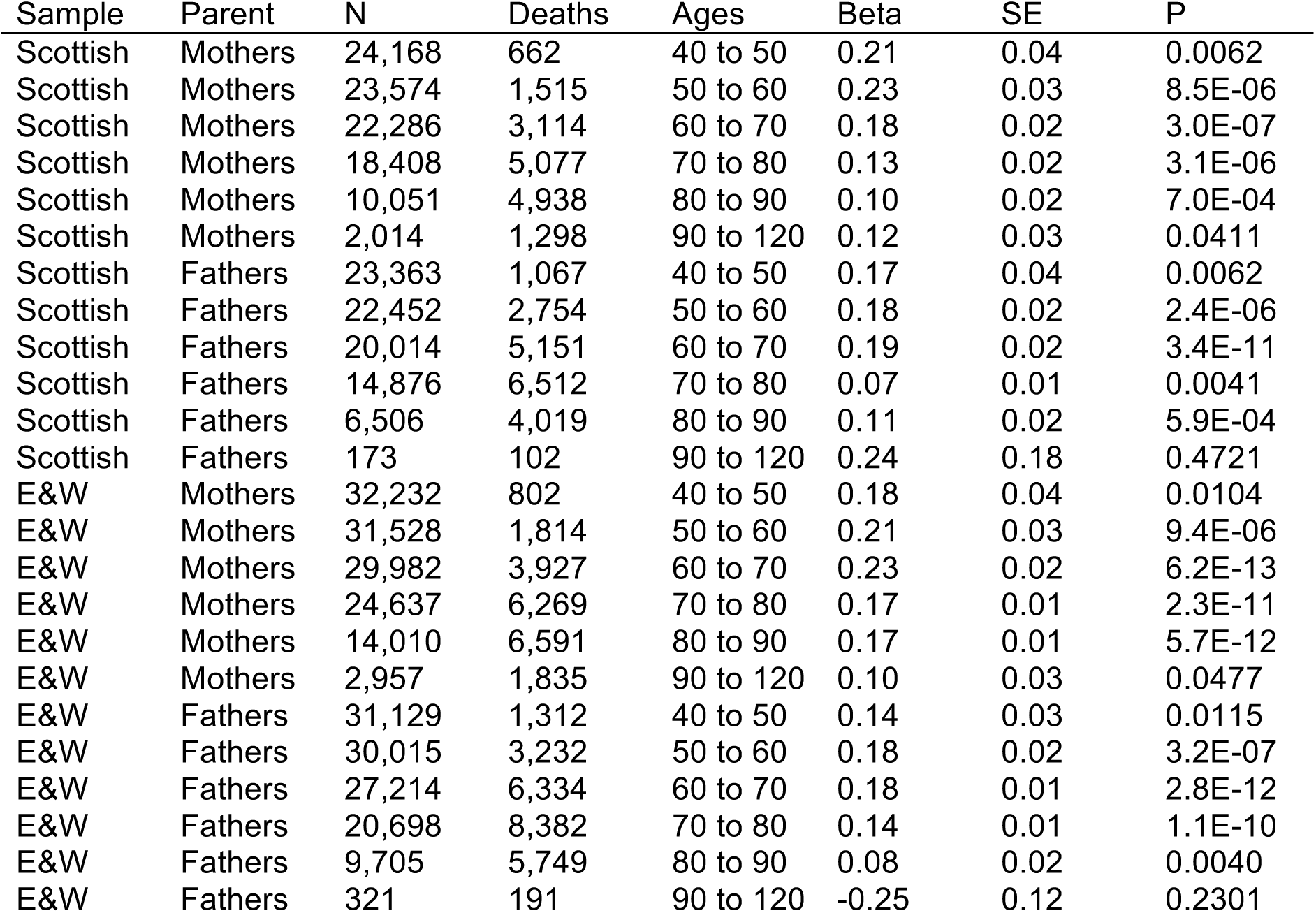
Sex and age-stratified association of polygenic score on lifespan. A polygenic risk score was made for each subject using GWAS results that did not include the subject sets under consideration. Parent survival information (age and alive/dead status) was stratified by sex and age. Sample – Out-of-sample subsets of UK Biobank individuals used for PGRS association (E& W: English and Welsh). N – Number of parental lifespans reported by sample individuals and used for the analysis (e.g. a parent aged 55 contributed to analysis of 40-50 and 50-60, but not 60-70). Deaths – Number of parental deaths within the age range reported by sample individuals. Ages – Lower limit to upper limit of age in analysis. Beta – log_e_(protection ratio) for 1 standard deviation in polygenic score in self in the age band (i.e. 2 x observed due to kin cohort method). SE – Standard error. P–P value of two sided test of association.

**Table S27: Associations of polygenic score with diseases in UK Biobank** [see Supplementary Information Excel File] A polygenic risk score was made for each subject using GWAS results that did not include the subject sets under consideration. Disease associations have been identified in the subjects, their siblings, or parents, at FDR 5%. Sample — Out-of-sample subsets of UK Biobank individuals used for PGRS association (E&W: English and Welsh). Kin – Family member for which the disease was reported. Trait – Disease reported by UK Biobank subject. N – Number of individuals tested. Cases – Number of reported individuals or kin carrying the disease. Beta – log OR of NOT carrying the disease per standard deviation of PGRS (i.e. positive beta indicates the PGRS protects from disease). SE – Standard Error. P – Two-sided P value. Q – Benjamin-Hochberg adjusted P value

